# Targeted Amplification and Genetic Sequencing of the Severe Acute Respiratory Syndrome Coronavirus 2 Surface Glycoprotein

**DOI:** 10.1101/2023.07.28.551051

**Authors:** Matthew W. Keller, Lisa M. Keong, Benjamin L. Rambo-Martin, Norman Hassell, Kristine Lacek, Malania M. Wilson, Marie K. Kirby, Jimma Liddell, D. Collins Owuor, Mili Sheth, Joseph Madden, Justin S. Lee, Rebecca J. Kondor, David E. Wentworth, John R. Barnes

## Abstract

The SARS-CoV-2 spike protein is a highly immunogenic and mutable protein that is the target of vaccine prevention and antibody therapeutics. This makes the encoding S-gene an important sequencing target. The SARS-CoV-2 sequencing community overwhelmingly adopted tiling amplicon-based strategies for sequencing the entire genome. As the virus evolved, primer mismatches inevitably led to amplicon drop-out. Given the exposure of the spike protein to host antibodies, mutation occurred here most rapidly, leading to amplicon failure over the most insightful region of the genome. To mitigate this, we developed SpikeSeq, a targeted method to amplify and sequence the S-gene. We evaluated 20 distinct primer designs through iterative *in silico* and *in vitro* testing to select the optimal primer pairs and run conditions. Once selected, periodic *in silico* analysis monitor primer conservation as SARS-CoV-2 evolves. Despite being designed during the Beta wave, the selected primers remain > 99% conserved through Omicron as of 2023-04-14. To validate the final design, we compared SpikeSeq data and National SARS-CoV-2 Strain Surveillance whole-genome data for 321 matching samples. Consensus sequences for the two methods were highly identical (99.998%) across the S-gene. SpikeSeq can serve as a complement to whole-genome surveillance or be leveraged where only S-gene sequencing is of interest. While SpikeSeq is adaptable to other sequencing platforms, the Nanopore platform validated here is compatible with low to moderate throughputs, and its simplicity better enables users to achieve accurate results, even in low resource settings.

## Introduction

In December 2019 an outbreak of pneumonia of unknown cause began in Wuhan, China (1). This illness (COVID-19) was found to be caused by a novel betacoronavirus, severe acute respiratory syndrome coronavirus 2 (SARS-CoV-2, SC2) (2). The virus quickly spread around the world, and in March 2020, the World Health Organization officially declared COVID-19 a pandemic (3). As of May 2023, COVID-19 has caused roughly 677 million infections and 6.9 million deaths (4).

As the COVID-19 pandemic progressed, waves of new variants spread (5), and mutations within the surface glycoprotein (spike) accumulated (6, 7). The spike protein is key to the viral replication cycle as its binding to the human angiotensin-converting enzyme 2 (ACE2) receptor initiates cellular entry of the virus (8). It also bears clinical significance as it is the target of vaccine prevention (9) and antibody therapeutics (10). The continual evolution of SARS-CoV-2 to evade immune pressures has led to a plethora of spike mutations that have been deleterious to vaccine effectiveness (11) and antibody neutralization (12-14). Importantly, many of these mutations are located within the receptor-binding domain (RBD) (15) where 90% of neutralizing antibodies target SARS-CoV-2 (16). As such, the spike protein encoding S-gene is an important sequencing target, and complete and accurate data for the S-gene is paramount for high quality surveillance information.

Genomic tools, such as the widely used Artic SARS-CoV-2 primer set, have required numerous updates to remain effective against new variants (17-20) (https://github.com/artic-network/artic-ncov2019/tree/master/primer_schemes/nCoV-2019). It can be challenging for surveillance labs, which are likely operating at surge capacity, to examine available alternative methods and validate revisions to the method for use. When overlooked, these limitations can lead to overt sequencing gaps or areas of low coverage, usually within the S-gene (21). This issue has occurred multiple times during the COVID-19 pandemic with major variant transitions (Origin strain to Alpha, Alpha to Delta, and Delta to Omicron). But this problem is not limited to major variant shifts. Variability in sequencing protocols and the design of sequencing primers in highly mutable regions of the SARS-CoV-2 spike protein causes intermittent sequencing dropouts, even with moderate amounts of variation. This is complicated further by the variability of organization for countrywide sequencing. Some countries have centralized health systems with more direct control of sequencing protocols and communication. Whereas other countries contract out sequencing to private labs, which can yield higher data volume but have more variability in sequencing methods and directness of communication. This can lead to protocol issues that are very slow to address, causing blind spots to critical regions of the spike protein as evolution occurs. As an illustration, we examined global surveillance data across the SARS-CoV-2 RBD over different time periods (**Figure S01**). During the transition from Delta to Omicron, this critical region was missing a significant amount of data. Moreover, the shape of that missing data resembles the amplicon 76 dropout known to affect the Artic SARS-CoV-2 primer set during the emergence of Omicron (19). The coverage across this region has since improved, but this does illustrate the issues of using mutation-sensitive amplification methods through a highly mutable region of a highly mutable virus.

Efforts have been made to focus surveillance to the S-gene; however, these methods have serious limitations. One such effort is to use eight overlapping amplicons (342-979 bp) and sanger sequencing to bring SARS-CoV-2 surveillance to low resource areas (22). Unfortunately, the need for eight RT-PCRs per sample and the use of sanger sequencing is costly, labor intensive, and seriously limits throughput potential. In an effort to improve the throughput of S-gene only sequencing, a modified version of Artic V3 SARS-CoV-2 primer set, HiSpike, was developed (23). HiSpike retains many of the limitations of the Artic SARS-CoV-2 protocol, most notably, the use of small amplicons (~400 bp) and the need for many primers to bind within the spike coding region. Using many primers to generate many small overlapping amplicons is not ideally suited to the surveillance of a rapidly evolving RNA virus and will likely again lead to sequencing dropouts due to primer mismatches. Indeed, multiple primers from both studies have conservation issues.

Because of these challenges, it was critical to develop a robust method for obtaining rapid sequence information, specifically for the S-gene. For this purpose, we developed SpikeSeq, a targeted method to amplify and sequence the S-gene (**Figure 1**). SpikeSeq uses four carefully selected and highly conserved primers (**Table 1**) to produce two overlapping amplicons that yield full coverage of the S-gene (**Figure 2**).

**Table 1:**
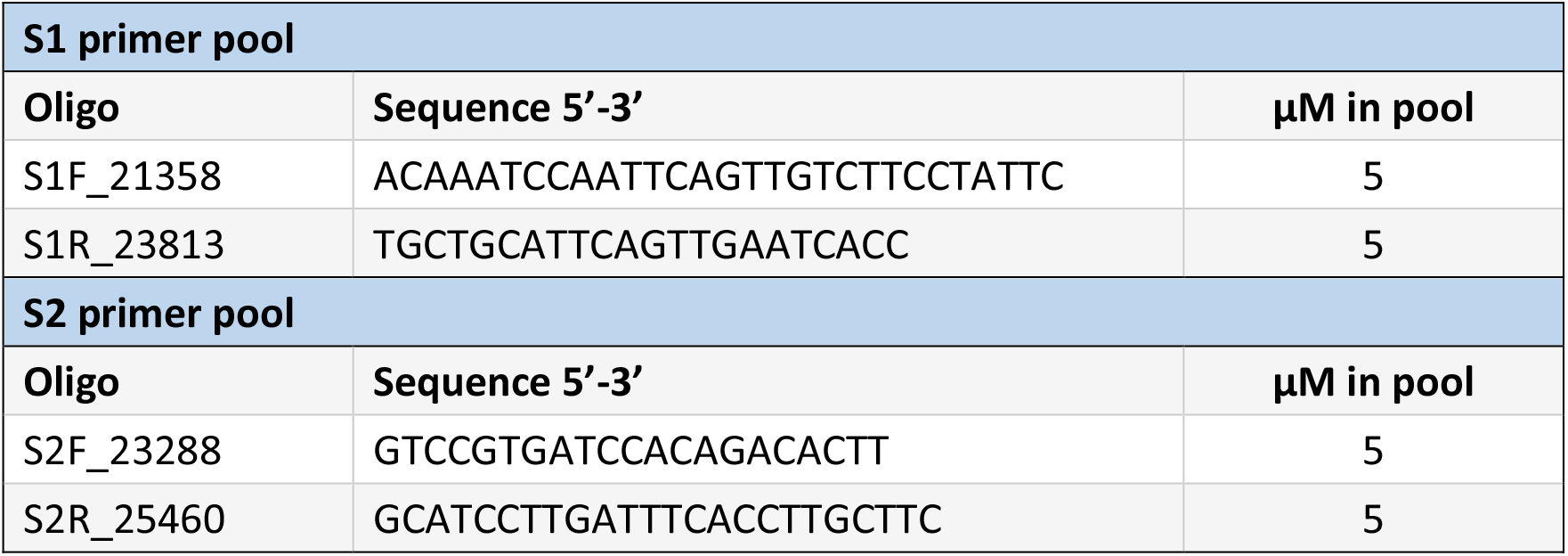
Primers. SpikeSeq primer sequences and working stock concentration.

**Figure 1:**
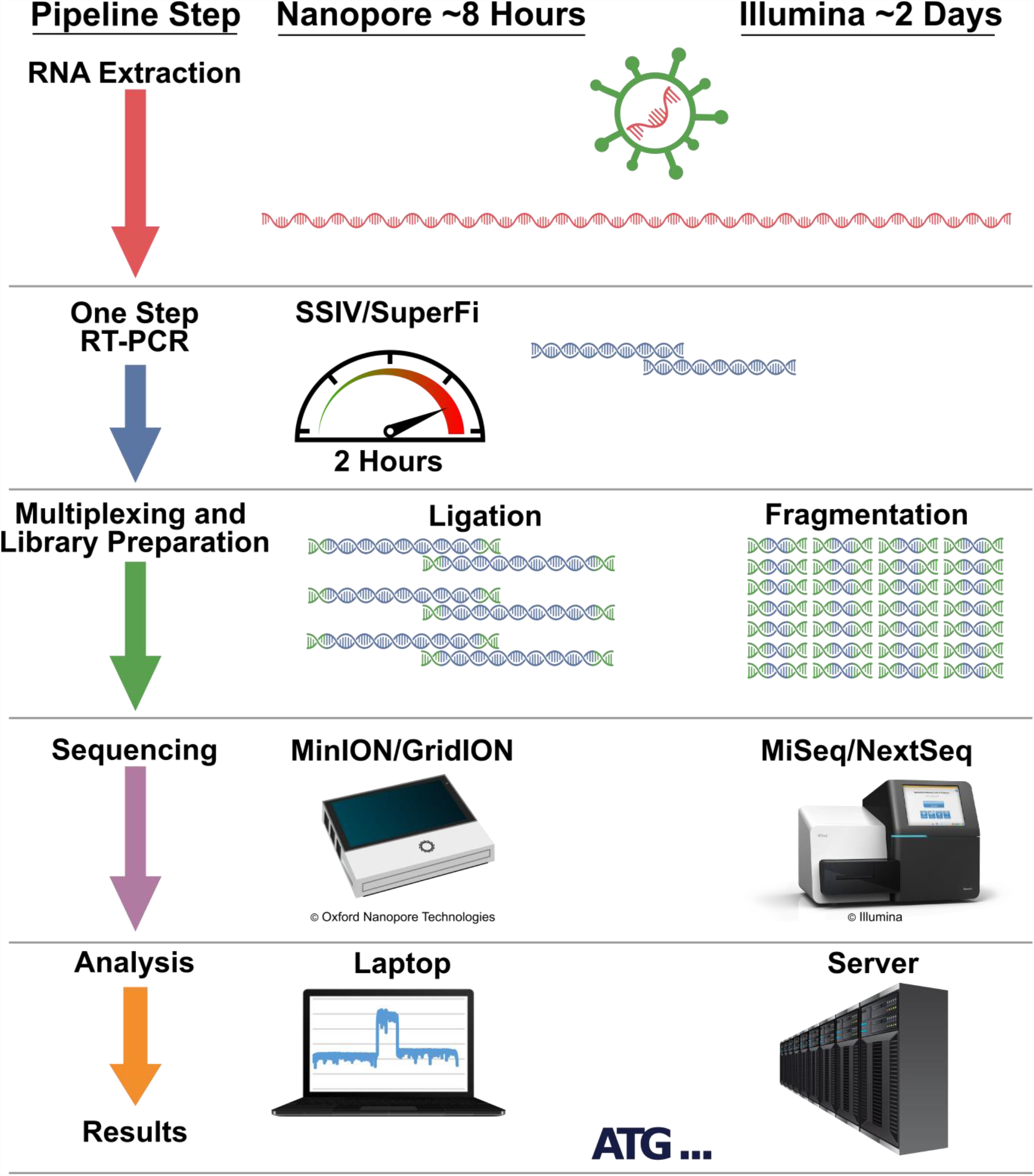
SpikeSeq Workflow. SpikeSeq is presented here as RT-PCR amplification and Nanopore sequencing. The workflow is designed to be flexible as the amplicons can be diverted to other sequencing platforms.

**Figure 2:**
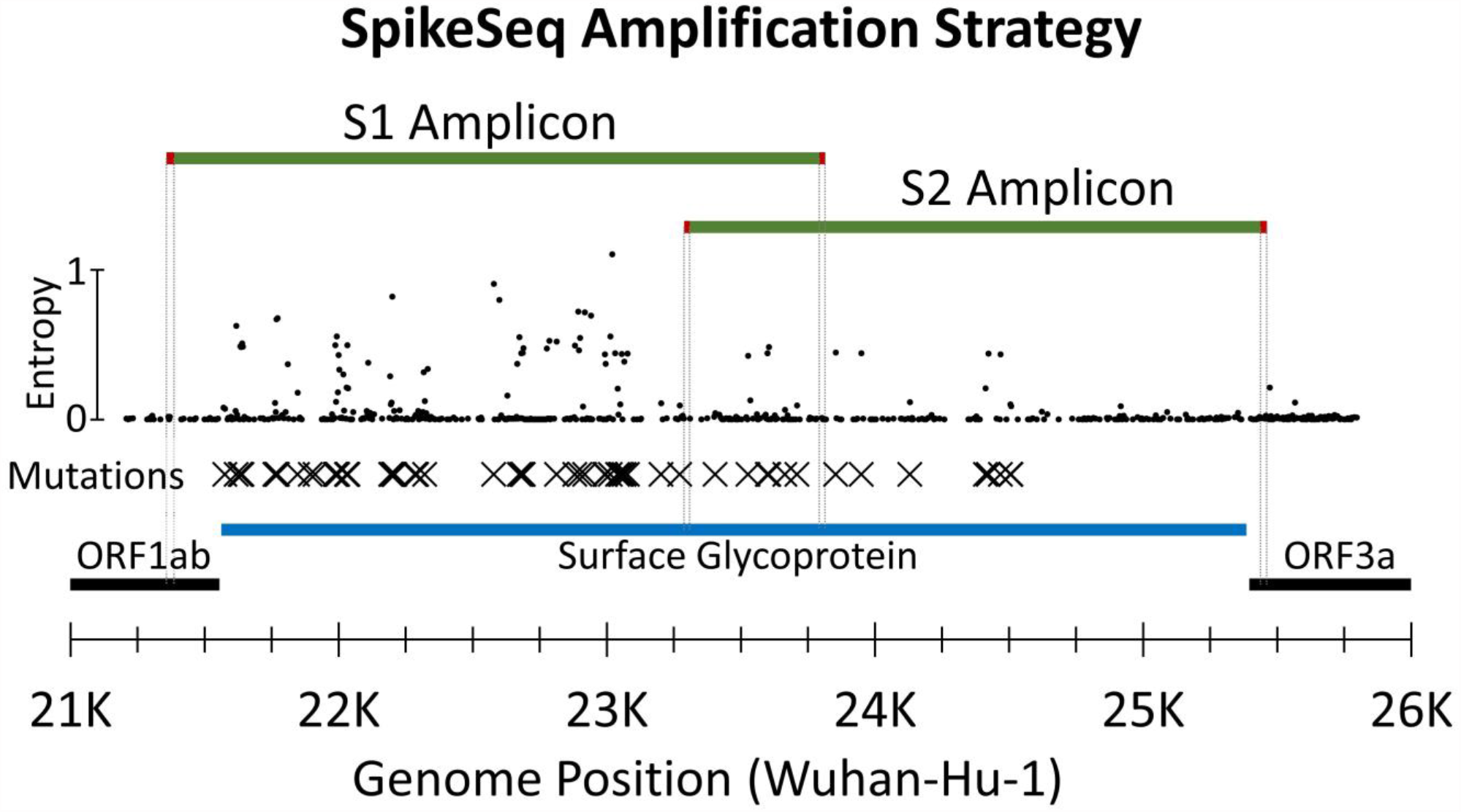
SpikeSeq Amplification Strategy. SpikeSeq amplicons are green with primer locations highlited in red and traced down to the ORFs. Diversity (entropy) across the region is plotted with small circles. Detected amino acid mutations are maked with Xs. SARS-CoV-2 ORFs are black with the Surface Glycoprotein ORF highlighted blue. Separate one-step RT-PCRs generate overlapping S1 and S2 amplicons that are 2.2 kb and 2.5 kb respectively. These amplicons extend beyond the coding region and overlap across the S1-S2 subunit cleavage site.

## Results

### Primer Selection and Validation

We used the conservation of all available SARS-CoV-2 sequences to identify (**Table S01**) and evaluate (**Table S02**) candidate primers. We identified three candidates for each of the 4 needed primers (S1F, S1R, S2F, and S2R) with additional candidates for S2R where SARS-CoV-2 (Wuhan-hu-1, NC_045512.2) and SARS-CoV-1 (NC_004718.3) shared identity. We eliminated those with < 95% conservation for all available SARS-CoV-2 sequences. By testing candidate primer combinations across an annealing temperature gradient (**Table S03; Figures S02-S03**), we were able to simultaneously eliminate possible combinations with poor performance and select 60°C as the annealing temperature. Finally, a limit of detection assay (**Tables S03-S04; Figure S04**) was used to select S1F_21358, S1R_23813, S2F_23288, and S2R_25460 as the final primers (**Table 1; Seq S01**).

We periodically monitor the conservation of these primers, and as of 2023-07-28, the selected primers remain highly conserved against SARS-CoV-2 using three months of US data, three months of global data, and all global data (**Figures S05-S16**). SpikeSeq primers also show some conservation against related coronaviruses (**Seq S02; Figure S17**). If a particular subvariant is of concern, we can perform a more focused conservation analysis. Such was the case with Omicron XBB, XBB.1.5, and derivatives. Analysis against those subvariants, as of 2023-01-18, demonstrated that our primers remained conserved (**Table S05**).

Our design results in an amplification strategy where two overlapping amplicons, each in their own RT-PCR reaction, span the entire gene. The four selected primers, which will generally be known as S1F, S1R, S2F, and S2R, avoid mutations and regions of high diversity (**Figure 2**).

### SpikeSeq Runs

We performed 14 Nanopore sequencing runs to validate and characterize SpikeSeq. A summary of these runs is available in the supplemental materials (**Table S06**).

### Sensitivity and Specificity

The limit of detection (LOD) via MinION flow cell sequencing was ~ 100 copies/μL and Ct 30 (**Table S07; Figure S18**). Via Flongle flow cell sequencing, the LOD was ~ 100 copies/μL and Ct 27 (**Table S08; Figure S19**).

No reads from the 84 NTCs mapped to SARS-CoV-2 (**Tables S09-S10; Figure S20-S21**).

### SpikeSeq Validation

We tested 377 samples via SpikeSeq for a pairwise comparison to National SARS-CoV-2 Strain Surveillance (NS3) whole-genome data. Of those, 321 samples passed SpikeSeq (**Seq S04**) and whole-genome sequencing (**Seq S05**) to be carried forward for further analysis. The S-gene consensus sequences were highly identical with 1,225,156 identities out of 1,225,185 positions (99.998% identical). Analyzing SpikeSeq data via Nextclade or Pangolin is limited by some group defining mutation residing outside of the S-gene. Still, with the widespread use of these analytical tools, we wanted to characterize the Nextclade results of SpikeSeq derived S-gene sequences in comparison to whole-genome sequences. Of the 281 samples that had a variant assignment (e.g., Delta or Omicron), Nextclade assignment of these variants was 100% concordant between SpikeSeq and whole-genome data. These assignments included the variants Alpha, Beta, Gamma, Delta, Epsilon, Eta, Iota, Lambda, Mu, and Omicron (24, 25). Identical clades were assigned for 91% of samples. As expected, clades with identical S-gene sequences, such as clades 21A (Delta) and 21J (Delta), were often conflated. Clades 21K (Omicron) and 21L (Omicron) were accurately assigned due to the S-gene diversity between those clades. Identical Nextclade_pango lineages were assigned for 72% of samples. Similar to clade identification, the resolution of SpikeSeq lineage identification is limited by a great number of named lineages and their identification being based off mutations outside the S-gene. (**Table S11**)

Importantly, SpikeSeq identified 4,428 mutations which includes all 4,422 spike protein mutations identified by whole-genome sequencing. For six samples, SpikeSeq identified one additional mutation each (**Table S11**). Further investigation of raw read data confirmed that these additional mutations were due to minor subpopulations at > 20% frequency amplified at variable proportions due to separate rounds of PCR between SpikeSeq and NS3. In any case, correctly identifying all 4,422 presumably true spike protein mutations reflects a high degree of accuracy that is more than sufficient for surveillance purposes.

A subset of 277 clinical specimens from the NS3 project were used for additional characterization of SpikeSeq. By comparing Ct values to SpikeSeq coverage results (**Table S12**), we found that 98-99% of samples with a Ct value less than 25 (n = 217) passed the coverage threshold of requiring ≥ 50x coverage at every position. For samples with Ct values between 25 and 30 (n = 44), 89% of samples passed. And for samples with Ct values over 30 (n = 16), 81% of samples passed (**Figure S22**).

For this subset of 277 clinical specimens, we split the three Nanopore libraries for loading on standard MinION flow cells (FLO-MIN106) for 72 hours and disposable Flongle flow cells (FLO-FLG001) for 24 hours. These flow cell types are known to have disparate sequencing yields, and indeed the Flongle flow cells produced just ~1% of the average coverage compared to the MinION flow cells. However, the coverage thresholds for SpikeSeq only requires a full assembly be made and ≥ 50x coverage at every position. Using those requirements, MinION flow cell sequencing passed 267/277 samples (96%), and Flongle flow cell sequencing passed 241/277 samples (87%). In other words, ~ 1% of the average coverage from Flongle flow cells passed 90% (241/267) as many samples with respect to MinION flow cells (**Table S13; Figure S23**).

For this same subset of 277 clinical specimens, we diverted a portion of the spike amplicons to Illumina sequencing, and 251 samples passed both techniques. We compared consensus level identity between spike amplicons sequenced via Nanopore (MIN) to those same amplicons sequenced via Illumina (ILL; **Seq S06**). For 251 samples, consensus sequences were highly identical with 958,236 identities out of 958,239 positions (99.9997% identical). This was expanded to a three-way comparison that includes the corresponding NS3 generated S-gene sequences (NS3; **Table S14**). The MINvNS3 consensus sequences were 99.9972% (958,212/958,239) identical, and the ILLvNS3 consensus sequences were 99.9969% (958,210/958,240) identical. All 251 samples had 100% identity between at least two of the three methods, and 20 samples had discrepant results. Because at least two of the methods always agreed, the discrepant results always appeared in pairs, were of identical magnitude, and shared a common method. For example, the three-way blast results of sample 3002648260 for MINvILL is 100% identical whereas MINvNS3 and ILLvNS3 are both 99.974% identical. This indicates the discrepancy lies with the NS3 derived S-gene consensus sample 3002648260. Of the 20 samples with discrepant results, 18 are due to discrepancies with the NS3 derived S-gene consensus, and 2 are due to discrepancies with the Illumina sequenced spike amplicons. This distribution of discrepancies is expected as the NS3 samples were independently amplified, processed, and analyzed. Ultimately though, these discrepancies are very minor and more than acceptable for surveillance purposes.

### Phylogenetics

We visualized the nextclade results in auspice to generate a tanglegram (**Figure S24**) of matching samples (n = 321) that passed both SpikeSeq and whole-genome sequencing. As detailed in **Table S11**, variant assignment was 100% concordant and clade assignment was highly concordant with ambiguities appearing for different clades with identical S-gene sequences.

## Discussion

We have developed and validated a robust method for amplifying and sequencing the SARS-CoV-2 surface glycoprotein. The length of the S-gene necessitated internal primers and separate overlapping RT-PCRs. Still, we were able to limit the total number of primers to four and the number of primers within the coding region of the S-gene to two. With so few primers required, we were able to evaluate many candidates for conservation and efficacy. We also ensured the primer binding sites avoided any major structural/functional elements. This design and process of primer selection gives SpikeSeq its best odds at avoiding mutations that might affect primer annealing. Indeed, despite being originally designed during the beta wave, the selected primers have remained highly conserved through Omicron as of 2023-04-14.

We validated this method against hundreds of clinical specimens collected for genomic surveillance by the NS3 project. We compared SpikeSeq data to the available whole-genome data to confirm that SpikeSeq accurately represents the S-gene amino acid mutations. Having only one amplicon in each reaction and two amplicons total, QC via electrophoresis can immediately reveal dropouts. This, combined with strict coverage requirements, ensures only complete and high quality S-gene sequencing data is reported.

As the pandemic progressed, the methods used by NS3 required several revisions, and occasionally, relied upon SpikeSeq for sequence completion (**Figure S25**) or confirmation of recombination (26). The speed of SpikeSeq has also proven useful during the floods of high priority samples associated with the appearance of new variants. In one case, SpikeSeq was used to confirm the presence of Omicron in the US Virgin Islands which allowed the territory to acquire antibody therapeutics best suited at the time for infection with the Omicron variant.

SpikeSeq can serve as a complement to whole-genome sequencing data with S-gene coverage gaps, be leveraged as a tool for projects in which only S-gene sequencing is of interest, and stand alone as a means of surveillance. SpikeSeq was evaluated and approved as a research use only method by the CDC Infectious Disease Test Review Board, and is currently being deployed to partner laboratories. In collaboration with The World Health Organization, we are hosting intensive week-long regional trainings where country representatives will gain hands-on experience and receive reagents, consumables, and a Mk1C sequencing device sufficient to perform a year of SARS-CoV-2 S-gene (and influenza A virus) surveillance. These regional trainings dedicate a great deal of time on foundational knowledge about the viruses themselves, the fundamentals of a quality surveillance effort, the importance of each step of data analysis and curation, and critically evaluating all step of the surveillance pipeline to ensure only quality data is submitted to public databases. While it is tempting to simply ship out point-and-click solutions, we want to develop a strong foundational knowledge about surveillance and data curation when training and equipping labs and countries new to next-generation sequencing (NGS) surveillance. This not only ensures the best use of resources, but gives those labs and countries the best chance at being successful in generating quality data and participating in this global surveillance effort. Moreover, because SpikeSeq targets a portion of the genome, a given amount of surveillance capacity could cover several times more samples compared to whole-genome sequencing on the same platform. The Nanopore platform used by SpikeSeq is compatible with low to moderate throughputs, and its simplicity better enables users to achieve accurate results, even in low resource settings. Finally, the relatively low capital expenditure makes this strategy an ideal starting point for public health laboratories new to NGS surveillance. As of April 2023, public health representatives from 59 countries have received this training with 23 more scheduled by the end of August 2023.

Whole-genome sequencing by a variety of methods will remain an integral part of SARS-CoV-2 surveillance, and we are not intending SpikeSeq to simply be a replacement. Whole-genome sequencing is the only way to properly assign phylogenetic relationships or monitor for amino acid mutations outside of the S-gene that can, for example, affect viral replication and pathogenesis (27). Moreover, quality whole-genome data is necessary to monitor primer conservation for any targeted amplification strategy.

SpikeSeq represents a refocusing on essential information needed from surveillance data. Whole-genome surveillance of SARS-CoV-2 has occasionally, and unfortunately, prioritized getting any result at the expense of sequence completeness and quality. As an example, eagerness to define new clades/lineages based on trivial differences has convoluted the classification of SARS-CoV-2 viruses and obscured the relationships between similar or disparate S-gene mutations that carry clinical significances. By focusing on the S-gene, imposing strict coverage and quality metrics, and applying lessons learned through surveillance of the diverse RNA influenza viruses, we hope to supplement SARS-CoV-2 surveillance with complete and quality reporting on the rapidly mutating S-gene.

## Materials and Methods

### Molecular Workflow

To amplify the S-gene, we produced overlapping amplicons (S1 and S2) via separate SuperScript™ IV One-Step RT-PCR System (Thermo Fischer Scientific, USA) reactions. The RT-PCR mixture contained: 4.25 μL nuclease-free water, 12.5 μL SSIV 2X reaction mix, 0.25 μL SSIV RT Mix, 5 μL S1 or S2 primer pairs, and 3 μL of RNA. The RT-PCR conditions are as follows: 10 minutes at 50°C; 2 minutes at 98°C; 40 cycles of 10 seconds at 98°C, 10 seconds at 60°C, and 1 minute 15 seconds at 72°C; a final elongation of 5 minutes at 72°C; and a hold at 4°C. Electrophoresis quality control was performed on individual RT-PCRs. After QC, corresponding S1 and S2 amplicons were combined, cleaned via SPRI beads (1x) with ethanol washes, and eluted into 15 μL of nuclease-free water.

Nanopore libraries were prepared using SQK-LSK109 and EXP-NBD196 and sequenced on GridION (Oxford Nanopore Technologies, UK) using FLO-MIN106 or FLO-FLG001 flow cells.

Laboratory procedures for RT-PCR and library preparation are available in the supplemental (**Text S01** and **Text S02**).

For Illumina sequencing, a portion of the cleaned amplicons were taken and prepared using the Nextera XT sample preparation kit. Since the SARS-CoV-2 S-gene amplicons are of a similar size to the influenza virus amplicons, they were processed via the standard influenza surveillance pipeline used by the CDC Genomics and Diagnostics Team (28, 29).

### Sequencing Data Analysis

During the sequencing run, we used the GridION MinKNOW to perform super-accuracy basecalling live (ont-guppy-for-gridion 5.0.17 or 5.1.13), to trim the barcodes, and to filter the reads. We trimmed primers using BBDuk (30), restricted the trimming using restrictleft=50 and restrictright=50, and referred to the primer sequences (**Seq S01**). We assembled reads using IRMA (https://wonder.cdc.gov/amd/flu/irma/irma.html) with the CoV-s-gene module (IRMA v.1.0.3 https://wonder.cdc.gov/amd/flu/irma/release_notes.html) and mapped to the S-gene reference (28). For a sample to pass SpikeSeq, it must meet coverage and quality metrics. Specifically, it must have a complete S-gene assembly, have at least 50x coverage at every position, and be free of frameshift mutations. Mutations were identified using Nextclade Web version 2.6.1 (https://clades.nextstrain.org; accessed September 30, 2022) SARS-CoV-2 without recombinants (24).

Analysis tools are available online https://cdcgov.github.io/MIRA (31).

### Primer Selection and Validation

We selected four primer target regions where S1F and S2R would lie outside of the S-gene coding region, and S1R and S2F would be on opposite sides of the S1/S2 cleavage site and avoid major structural elements. We identified multiple sets of candidate primers for each S1F, S1R, S2F, and S2R. For S2R, we also evaluated an area where SARS-CoV-2 (Wuhan-hu-1, NC_045512.2) and SARS-CoV-1 (NC_004718.3) shared identity (**Table S01**). During the Beta wave (March 2021), we evaluate the conservation of primer candidates against 476,466 SARS-CoV-2 genomes (**Table S02**). Twenty primer combinations were tested (**Table S03**). We initially screened the candidate primer pairs across a temperature gradient using RNA from B.1.351 (Beta) with a ct 25 as determined by the Flu SC2 multiplex assay (32). We used an LOD of B.1.351 (Beta) from ct 14-30 (846k to 16 copies/μL; **Table S04**) to finalize the primer selection. The presence of amplicons was determined using a QIAxcel HT fragment analyzer.

We monitored the conservation of the primers by downloading data from GISAID. Downloaded genomic data was aligned to the Wuhan-Hu-1 reference (NCBI accession MN908947.3) genome using SSW (33). Aligned genome primer regions were regularly compared for mismatches against each individual primer sequence. This information was used to highlight potential assay issues with new emerging variants. We downloaded diversity (entropy) data from Nextstrain (https://nextstrain.org/ncov/gisaid/global/6m; accessed March 6, 2023) (24).

### Sensitivity and Specificity

To measure the absolute limit of detection, we used a custom synthetic RNA fragment from Twist Bioscience (CA, USA) based on the Delta lineage virus hCoV-19/USA/CO-CDC-MMB09467199/2021. The sequence for this fragment (TwistDeltaFragment_4276451.fasta) is available in the supplemental materials (**Seq S03**). The 4,626-nucleotide fragment spans the S-gene and extends into neighboring genes. The synthetic fragment aliquots were delivered at 629,000 copies/μL as determined via manufacturer ddPCR. To measure the viral limit of detection, we used a propagated isolate of Delta SARS-CoV-2 and measured the Ct value of the serial dilutions using the Flu SC2 multiplex assay (32). The limit of detection was determined by the most dilute sample to pass coverage and quality thresholds for all the replicates.

We prepared RT-PCR master mixes for triplicate limit of detection assays using both synthetic and viral material. The dilution series were a 5-fold serial dilution through 7 steps with a water NTC as the 8^th^ step. LOD amplicons were split at the end-prep stages for sequencing on both MinION and flongle flow cells. For sequencing on MinION flow cells, we included 48 additional water NTCs. For sequencing on Flongle flow cells, we included 24 additional A549 RNA (Rp Ct 22) NTCs.

### SpikeSeq Validation

To validate this method, we tested a total of 377 specimens from the NS3 project. We started with a retrospective analysis of 277 clinical specimens that were collected from March to August 2021 and that capture the diversity of SARS-CoV-2 into the Delta wave. During the omicron wave, we continued SpikeSeq validation concurrently with NS3. These additional 100 samples were collected from November 2021 to January 2022. Of these 377 samples, 321 passed SpikeSeq (**Seq S04**) and whole-genome sequencing (**Seq S05**) to be carried forward for further analysis.

We compared matching samples (n = 321) that passed both SpikeSeq and whole-genome sequencing using ncbi-blast+/2.9.0 (34) and Nextclade Web version 2.6.1 (https://clades.nextstrain.org; accessed September 30, 2022) SARS-CoV-2 without recombinants (24). Using the output of Nextclade, we evaluated the concordance of variant, clade, and lineage assignment. We also compared the reported S-gene amino acid mutations for complete matches of corresponding samples and by counting individual mutations for corresponding samples (**Table S11**).

A subset of 277 samples through Delta was used to compare Ct values to coverage, Nanopore sequencing yield on two flow cell types (FLO-MIN106 versus FLO-FLG001), and Nanopore sequencing accuracy to Illumina sequencing. Each time the RNA was thawed, we tested it with the Flu SC2 multiplex assay (32) to determine the Ct value and amplified the S-gene using the methods presented here. For samples with an undefined Ct value (n =2), a Ct value of 40 was assigned. We then split the spike amplicons to both Illumina and Nanopore sequencing methods. For Nanopore sequencing, we prepared libraries using the methods described here and loaded both standard MinION flow cells (FLO-MIN106) for 72 hours and disposable Flongle flow cells (FLO-FLG001) for 24 hours.

All 277 samples from this subset (pass or fail) were used to assess the relative pass rates of standard MinION flow cells (FLO-MIN106) versus Ct value (**Table S12 and Figure S22**) and versus disposable Flongle flow cells (FLO-FLG001; **Table S13 and Figure S23**).

From that subset of 277 samples, 251 samples passed both Nanopore (FLO-MIN106) and Illumina sequencing of the SpikeSeq (**Seq S06**). For each of these 251 samples, we used ncbi-blast+/2.9.0 (34) to generate a three-way comparison between: SpikeSeq amplification and Nanopore sequencing (MIN), Illumina sequencing of those same amplicons (ILL), and NS3 surveillance results for the S-gene (NS3; **Table S14**).

### Phylogenetics

We compared matching samples (n = 321) that passed both SpikeSeq and whole-genome sequencing using Nextclade Web version 2.6.1 (https://clades.nextstrain.org; accessed September 30, 2022) SARS-CoV-2 without recombinants (24). From this analysis, we exported the phylogenetics and visualized them with Auspice (https://auspice.us; accessed October 6, 2022). We added a metadata sheet to label and highlight added sequences above the backbone sequences.

### Primer Kit Manufacturing

CDC Division of Scientific Resources manufactured primers for use in this study and distribution to public health laboratories. The Oligo Synthesis Laboratory synthesized the primers, purified via HPLC, and verified by mass spectrophotometry. Following initial synthesis and purification, we received three QC aliquots for limit of detection analysis and excess material for use in this study. The remaining material (5 mmol each primer) was then transferred to the Diagnostic Manufacturing Laboratory for stochiometric mixing of forward and reverse primers, dispensing, drying, and kit assembly. We received three aliquots for QC testing.

## Supplemental Material

Supplemental material for this article may be found at https://figshare.com/articles/dataset/Supplemental_Material/22762076.

Supplemental legends are available in the supplemental (**Text S03**).

## Acknowledgements

We thank the CDC/NCEZID/Division of Scientific Resources/Biotechnology Core Facility Branch/Oligo Synthesis Laboratory for synthesizing the primers used in this study.

We thank the CDC/NCEZID/Division of Scientific Resources/Reagent and Diagnostic Services Branch/Diagnostic Manufacturing Laboratory for manufacturing primer kits.

## Data availability

Corresponding SpikeSeq (Nanopore sequencing) S-gene consensus sequences and NS3 whole-genome consensus sequences are available in the supplemental materials (n = 321 each; **Seq S04-S05**). SpikeSeq amplification and Illumina sequencing derived S-gene consensus sequences (n = 251) are available in the supplemental materials (**Seq S06**). https://figshare.com/articles/dataset/Supplemental_Material/22762076

FASTQ reads (that BLAT matched to IRMA reference) are available online at NCBI under BioProject: PRJNA999712. The BioSamples (n=810) include the 321 primary validation samples (320 FLO-MIN106 and 1 FLO-FLG001), the 238 flongle yield replicates that passed, and 251 Illumina accuracy replicates that passed. https://www.ncbi.nlm.nih.gov/bioproject/PRJNA999712

## Tables and Figures

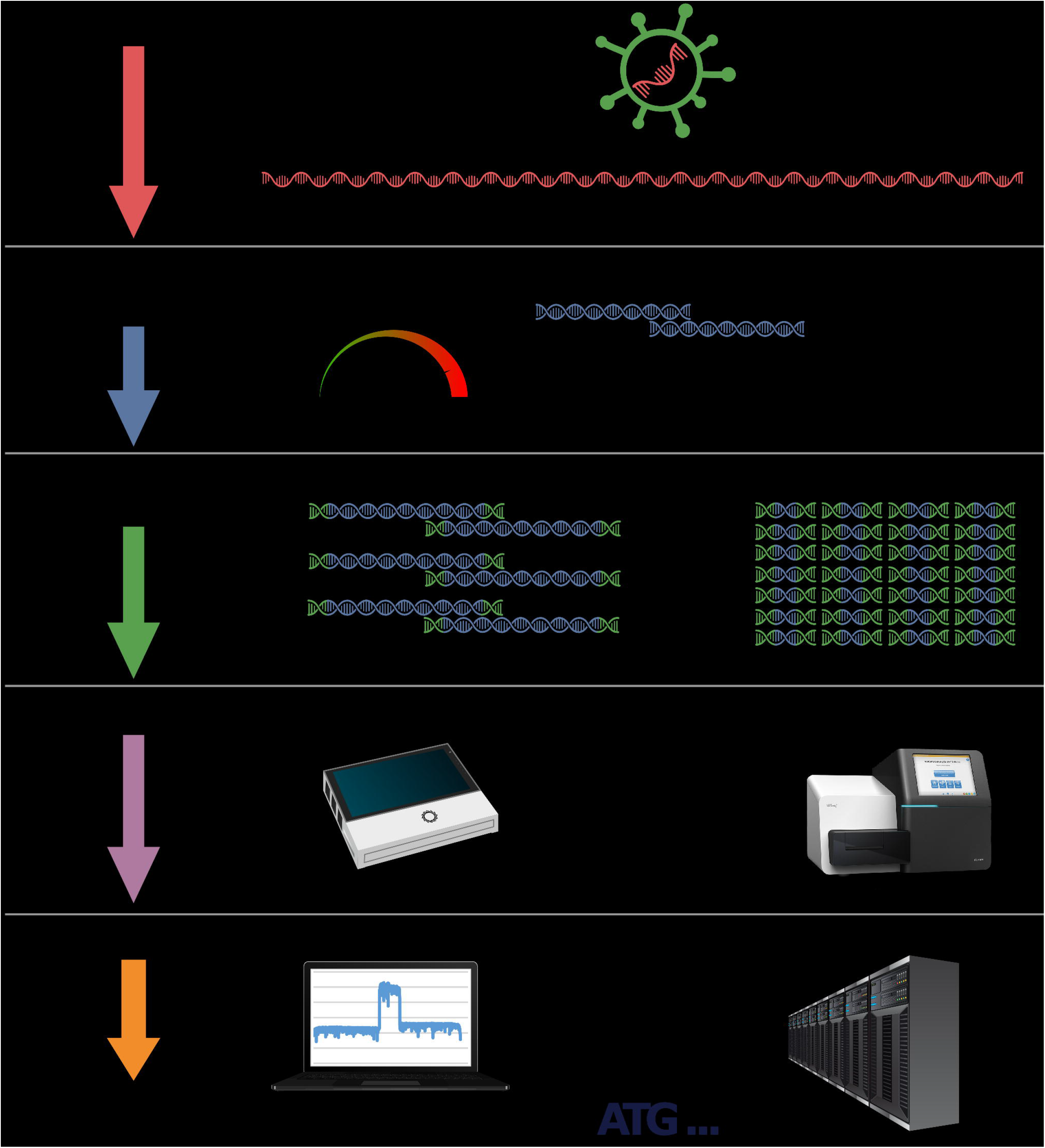

