## SupplementalMaterials for "Targeted Amplification and Genetic Sequencing of the Severe Acute Respiratory Syndrome Coronavirus 2 Surface Glycoprotein": Figure_S02_S1_TemperatureGradient_Fragments.pdf

| Report Overview | |
|---|---|
| Report Date: | 3/2/2021 2:26:19 PM |
| Experiment Name: | 2021-03-02_Matt1stepS1 |
| Cartridge ID: | C191210A09 |
| Instrument ID: | 30334 |

| Annealing Temperature (°C) | 68.0 | 67.0 | 65.4 | 62.9 | 59.9 | 57.5 | 55.9 | 55.0 |
|---|---|---|---|---|---|---|---|---|

Primer Combination S1-1

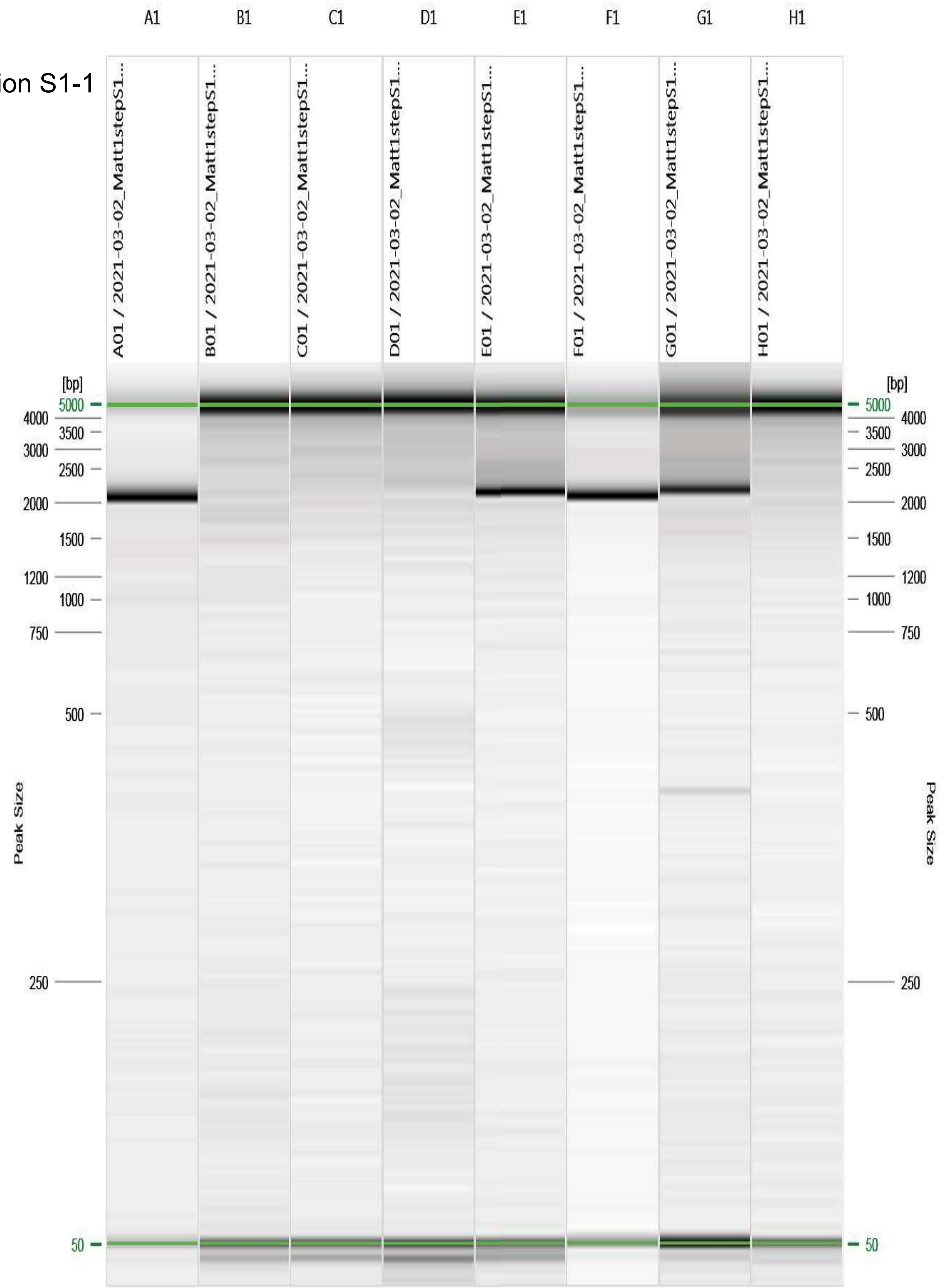


Figure: 1 - 2021-03-02_Matt1stepS1/R:1 E:1 #1

| Annealing Temperature (°C) | 68.0 | 67.0 | 65.4 | 62.9 | 59.9 | 57.5 | 55.9 | 55.0 |
|---|---|---|---|---|---|---|---|---|

Primer Combination S1-2

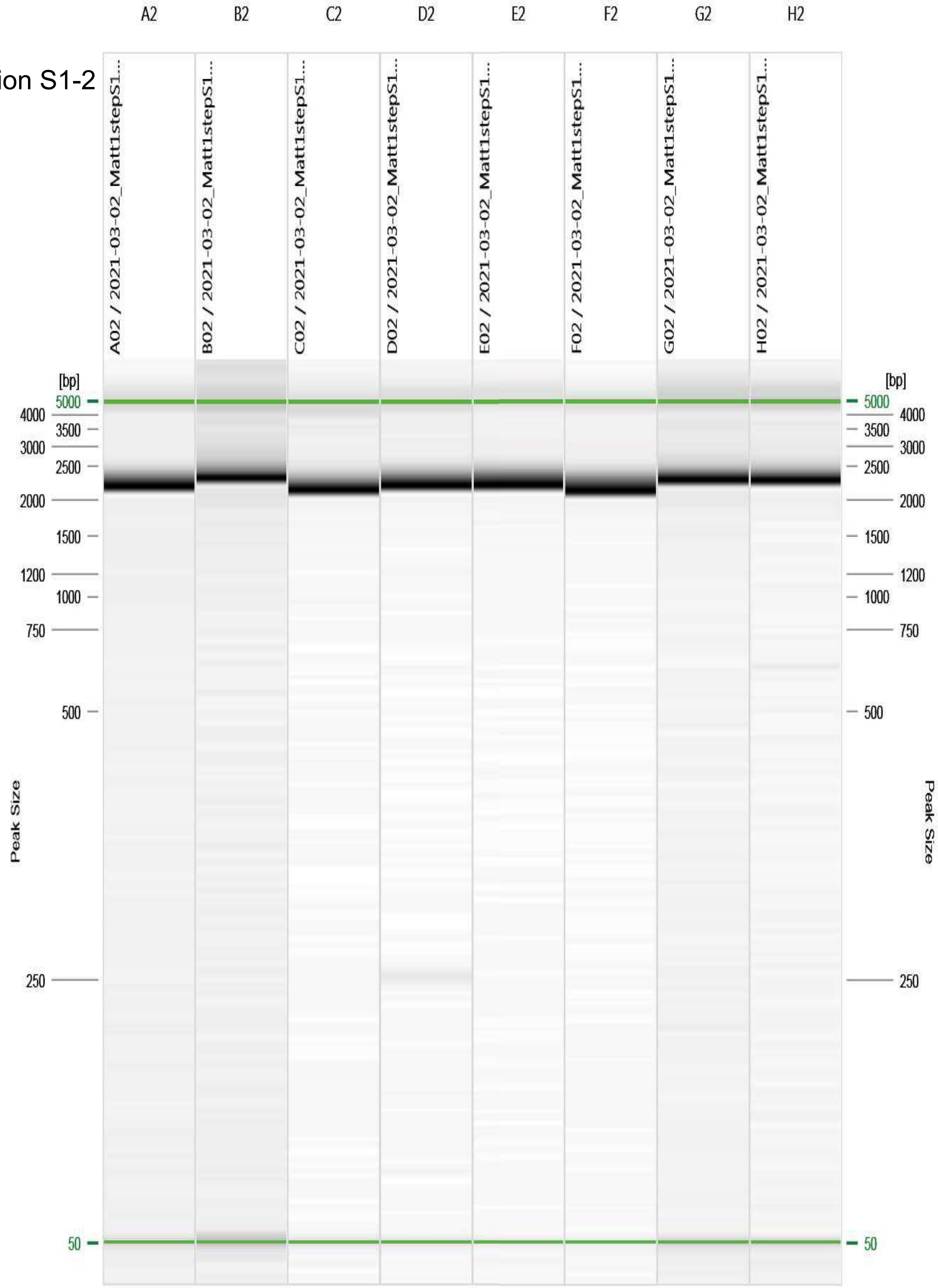


Figure: 2 - 2021-03-02_Matt1stepS1/R:1 E:1 #2

| Annealing Temperature (°C) | 68.0 | 67.0 | 65.4 | 62.9 | 59.9 | 57.5 | 55.9 | 55.0 |
|---|---|---|---|---|---|---|---|---|

Primer Combination S1-3

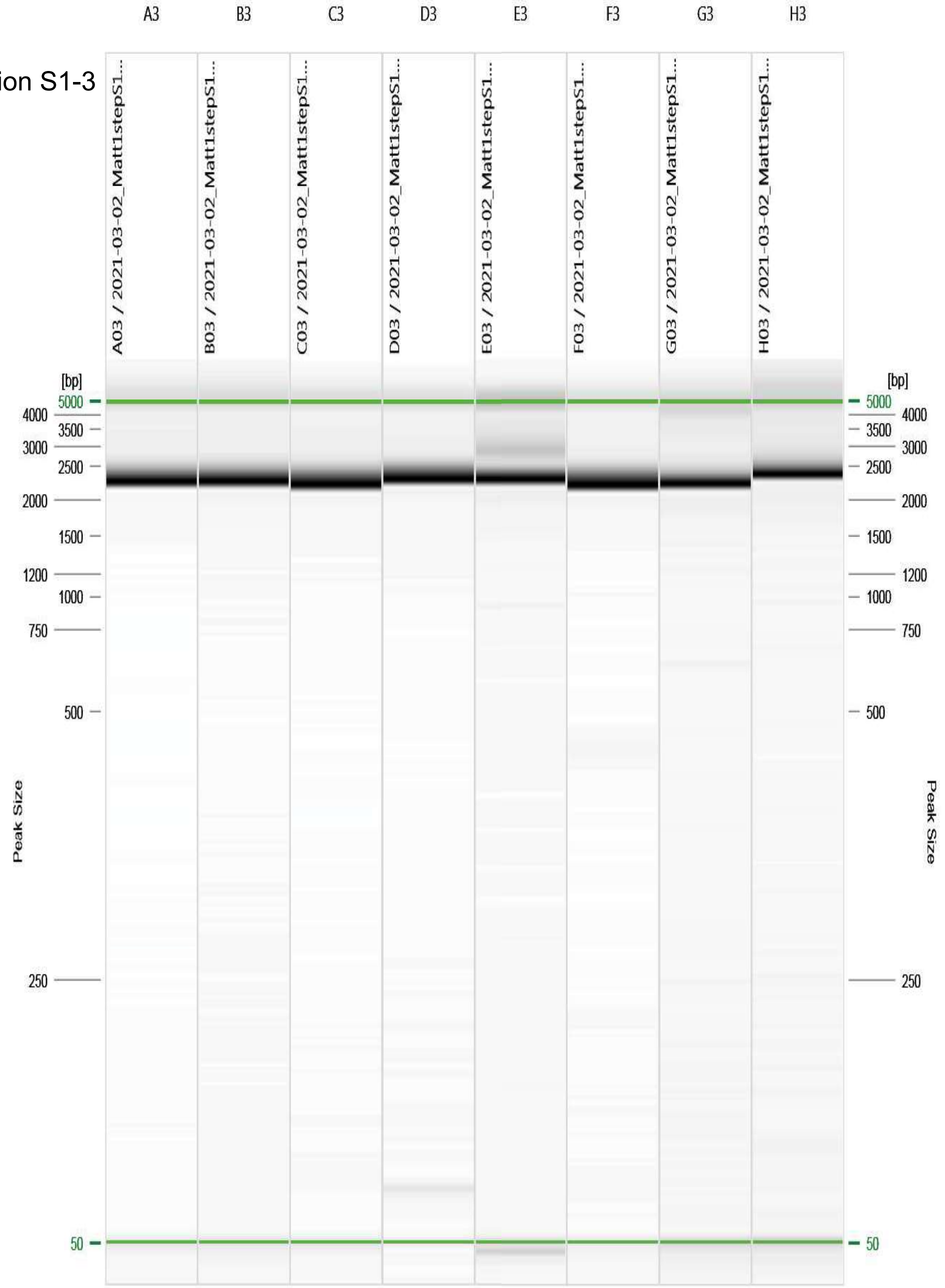


Figure: 3 - 2021-03-02_Matt1stepS1/R:1 E:1 #3

| | | | | | | | | |
|---|---|---|---|---|---|---|---|---|
| Annealing Temperature (°C) | 68.0 | 67.0 | 65.4 | 62.9 | 59.9 | 57.5 | 55.9 | 55.0 |

Primer Combination S1-4

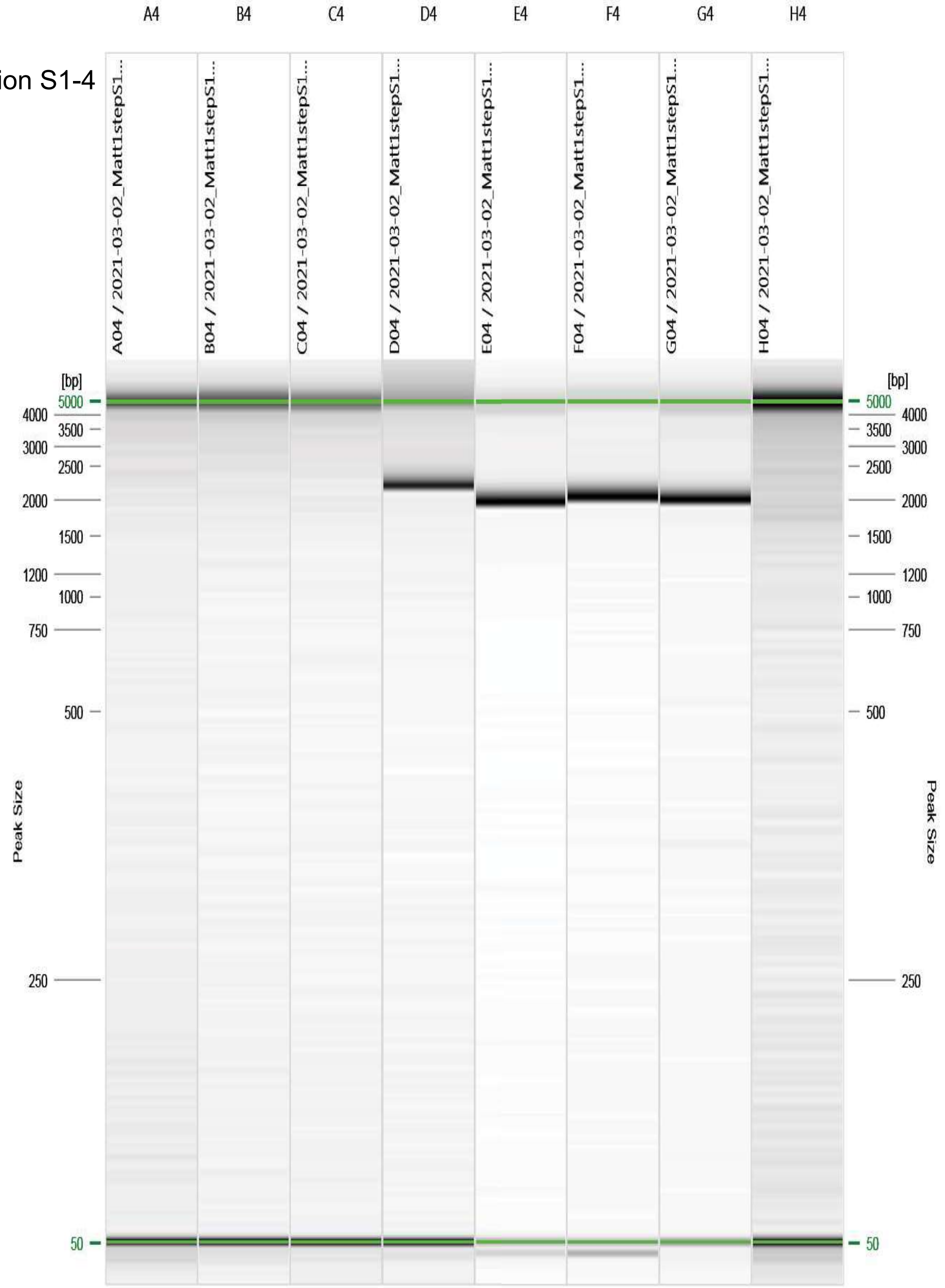


Figure: 4 - 2021-03-02_Matt1stepS1/R:1 E:1 #4

| Annealing Temperature (°C) | 68.0 | 67.0 | 65.4 | 62.9 | 59.9 | 57.5 | 55.9 | 55.0 |
|---|---|---|---|---|---|---|---|---|

Primer Combination S1-5

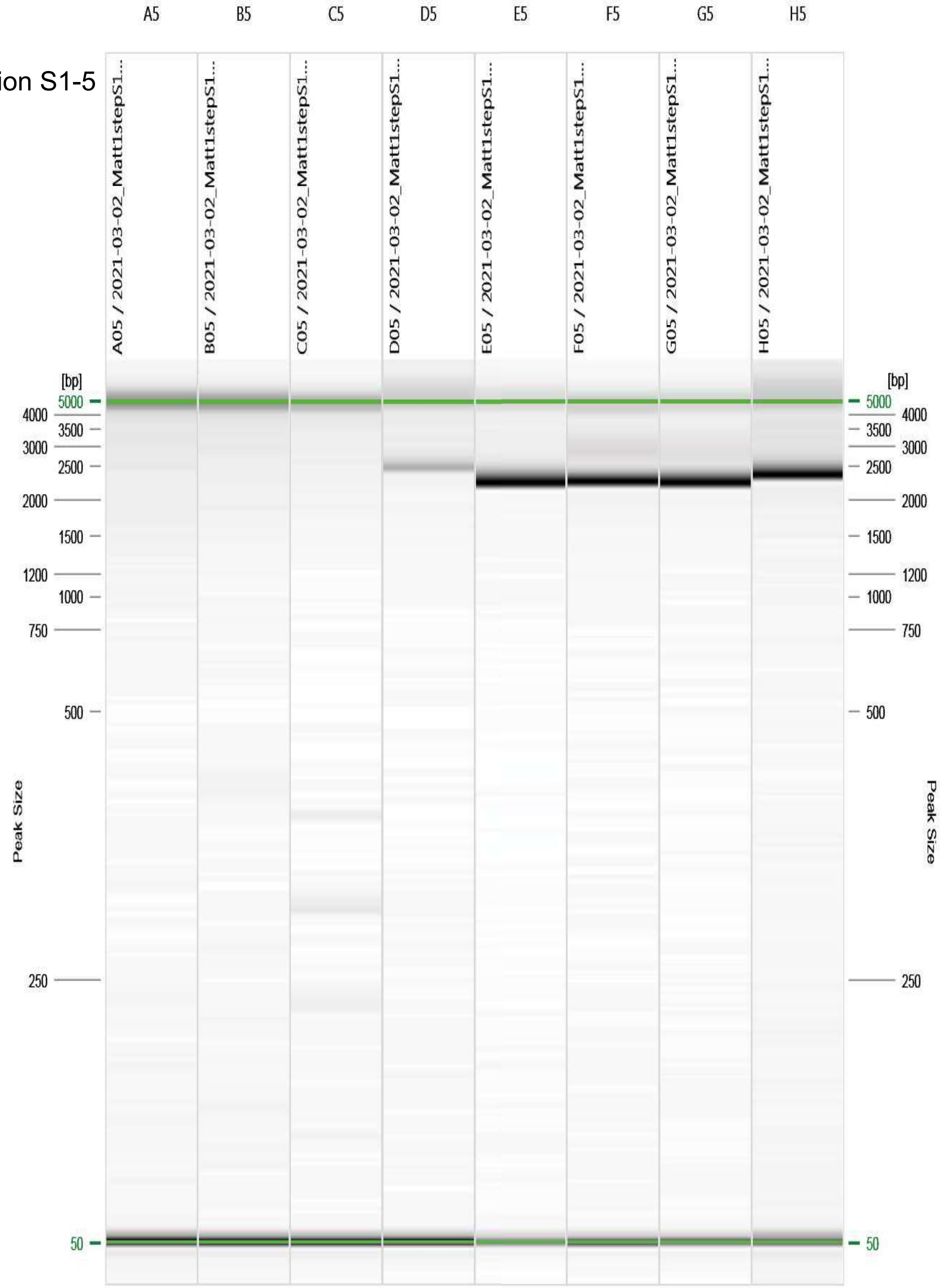


Figure: 5 - 2021-03-02_Matt1stepS1/R:1 E:1 #5

| Annealing Temperature (°C) | 68.0 | 67.0 | 65.4 | 62.9 | 59.9 | 57.5 | 55.9 | 55.0 |
|---|---|---|---|---|---|---|---|---|

Primer Combination S1-6

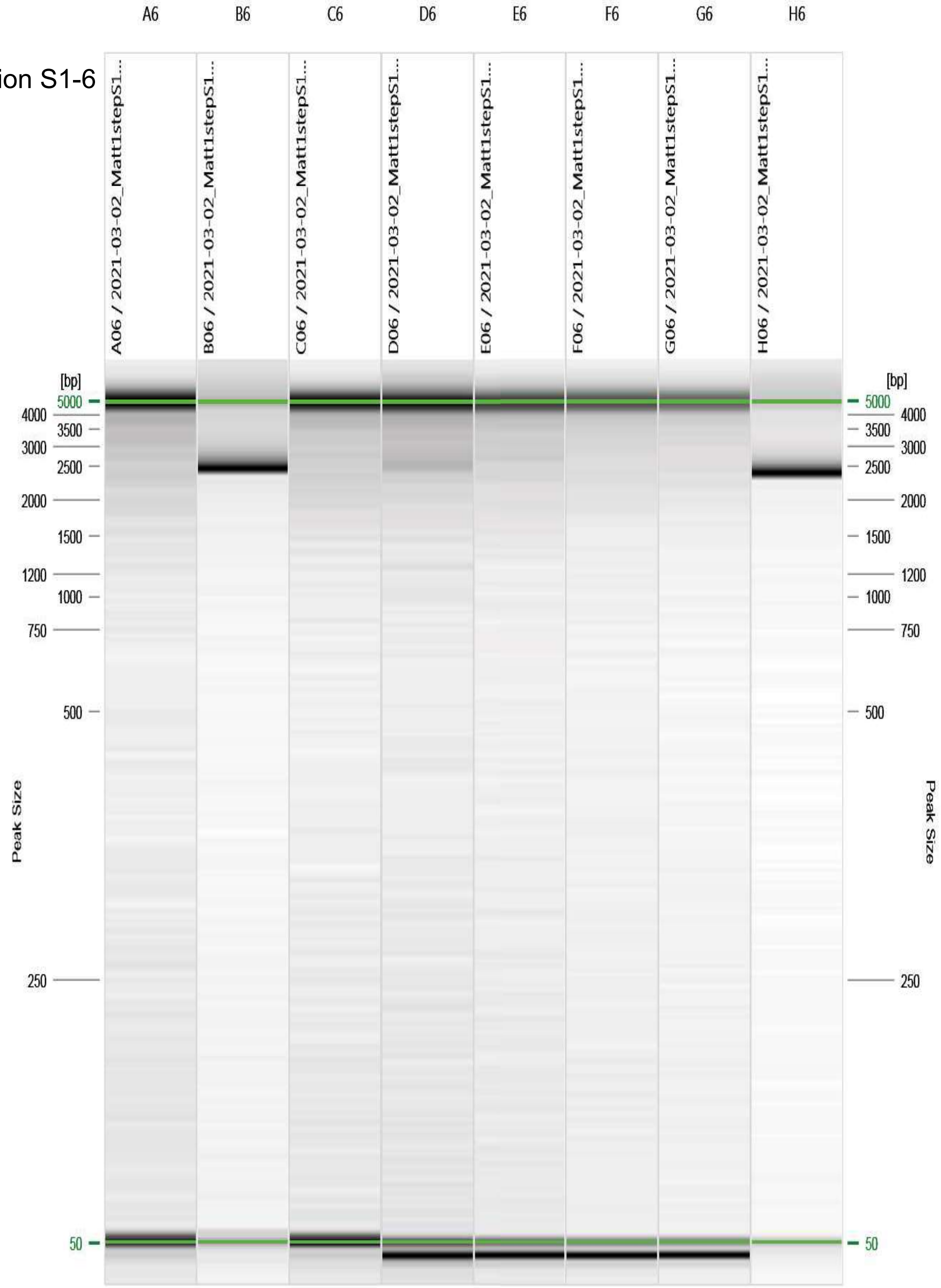


Figure: 6 - 2021-03-02_Matt1stepS1/R:1 E:1 #6

| Annealing Temperature (°C) | 68.0 | 67.0 | 65.4 | 62.9 | 59.9 | 57.5 | 55.9 | 55.0 |
|---|---|---|---|---|---|---|---|---|

Primer Combination S1-7

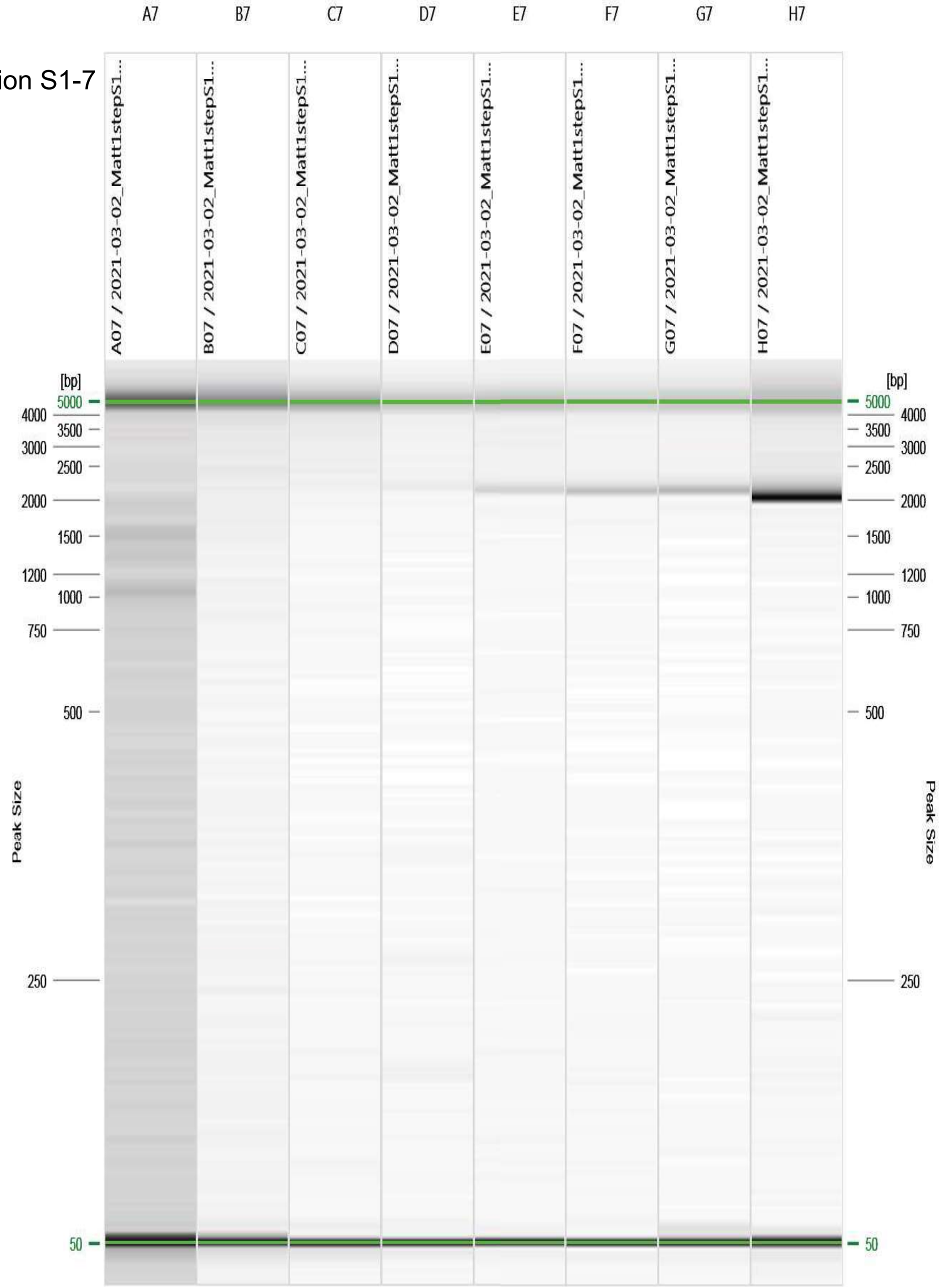


Figure: 7 - 2021-03-02_Matt1stepS1/R:1 E:1 #7

| Annealing Temperature (°C) | 68.0 | 67.0 | 65.4 | 62.9 | 59.9 | 57.5 | 55.9 | 55.0 |
|---|---|---|---|---|---|---|---|---|

Primer Combination S1-8

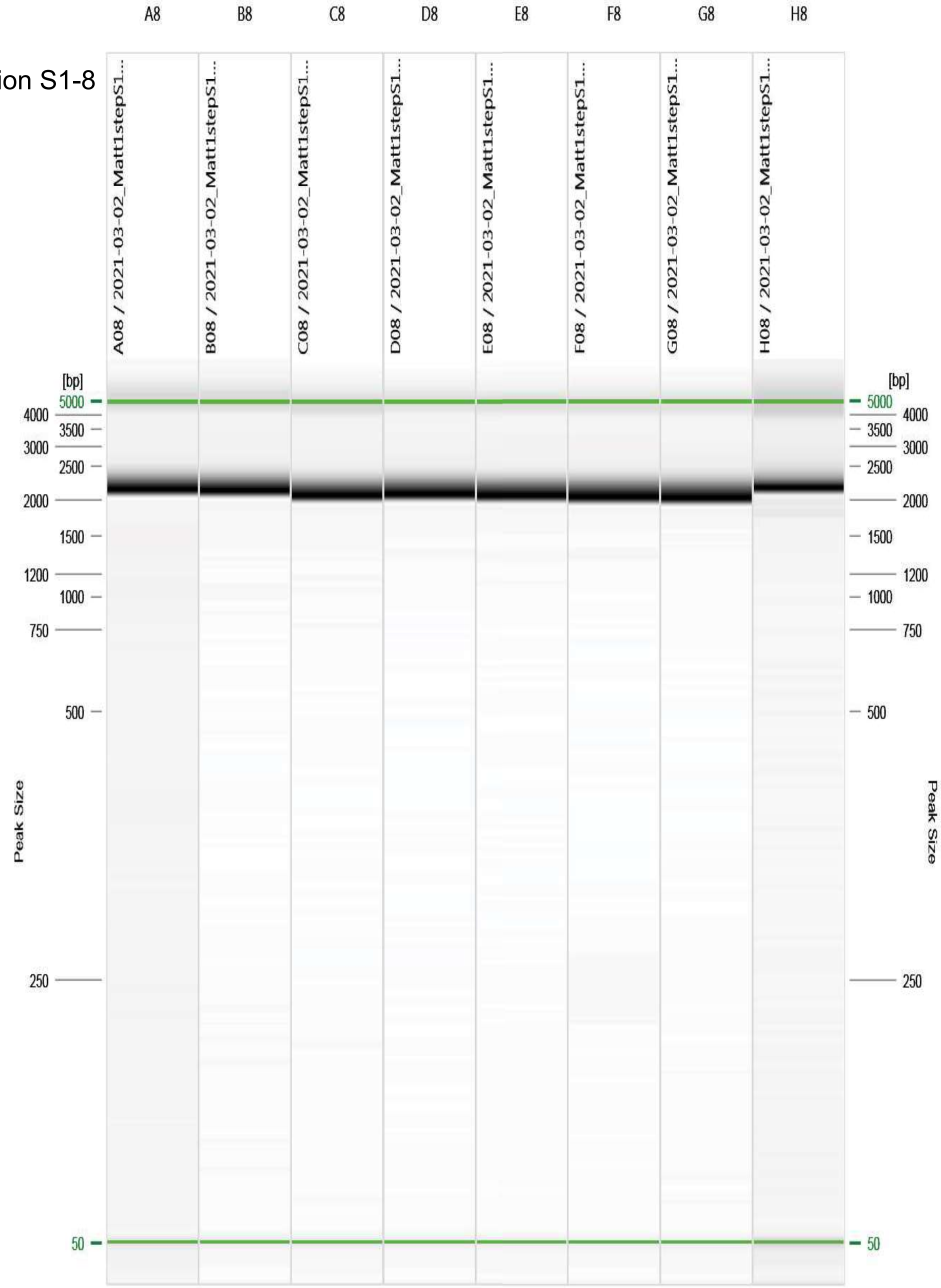


Figure: 8 - 2021-03-02_Matt1stepS1/R:1 E:1 #8

| Annealing Temperature (°C) | 68.0 | 67.0 | 65.4 | 62.9 | 59.9 | 57.5 | 55.9 | 55.0 |
|---|---|---|---|---|---|---|---|---|

Primer Combination S2-9

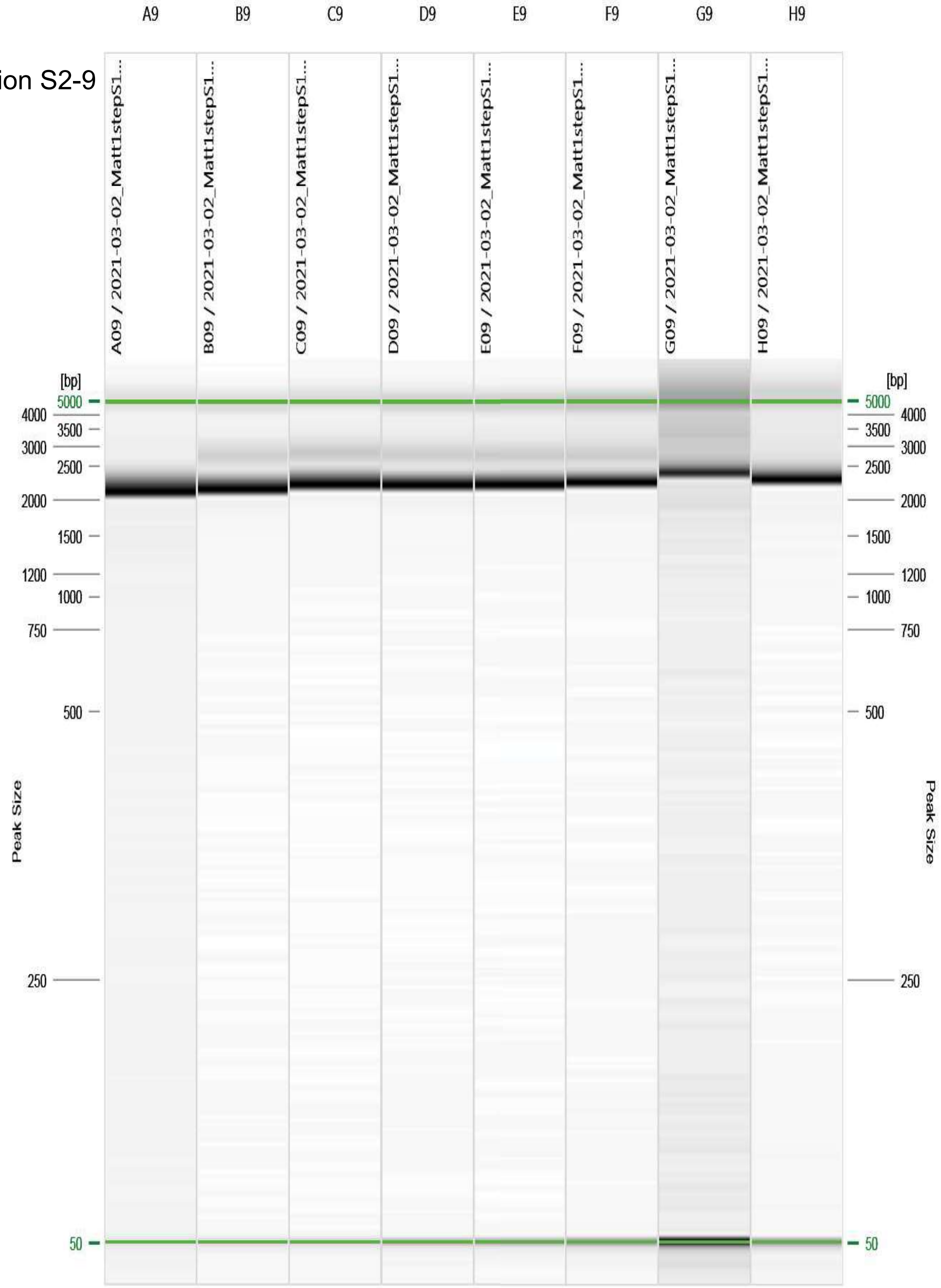


Figure: 9 - 2021-03-02_Matt1stepS1/R:1 E:1 #9
