## SupplementalMaterials for "Targeted Amplification and Genetic Sequencing of the Severe Acute Respiratory Syndrome Coronavirus 2 Surface Glycoprotein": Figure_S03_S2_TemperatureGradient_Fragments.pdf

| Report Overview | |
|---|---|
| Report Date: | 3/2/2021 4:40:02 PM |
| Experiment Name: | 2021-03-02_Matt1stepS2 |
| Cartridge ID: | C191210A09 |
| Instrument ID: | 30334 |

| Annealing Temperature (°C) | 68.0 | 67.0 | 65.4 | 62.9 | 59.9 | 57.5 | 55.9 | 55.0 |
|---|---|---|---|---|---|---|---|---|
| | A1 | B1 | C1 | D1 | E1 | F1 | G1 | H1 |

Primer Combination S2-1

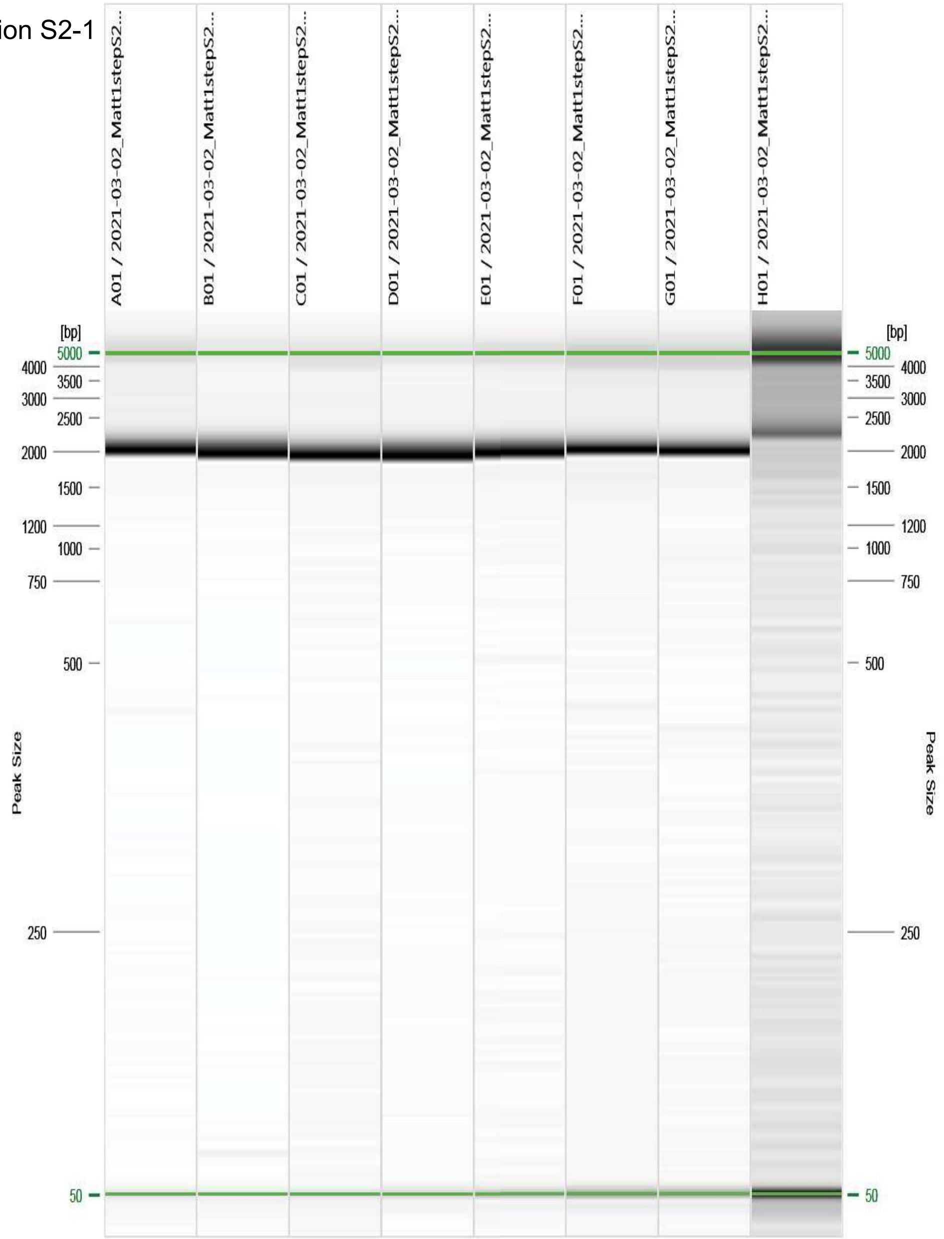


Figure: 1 - 2021-03-02_Matt1stepS2/R:1 E:1 #1

| Annealing Temperature (°C) | 68.0 | 67.0 | 65.4 | 62.9 | 59.9 | 57.5 | 55.9 | 55.0 |
|---|---|---|---|---|---|---|---|---|

Primer Combination S2-2

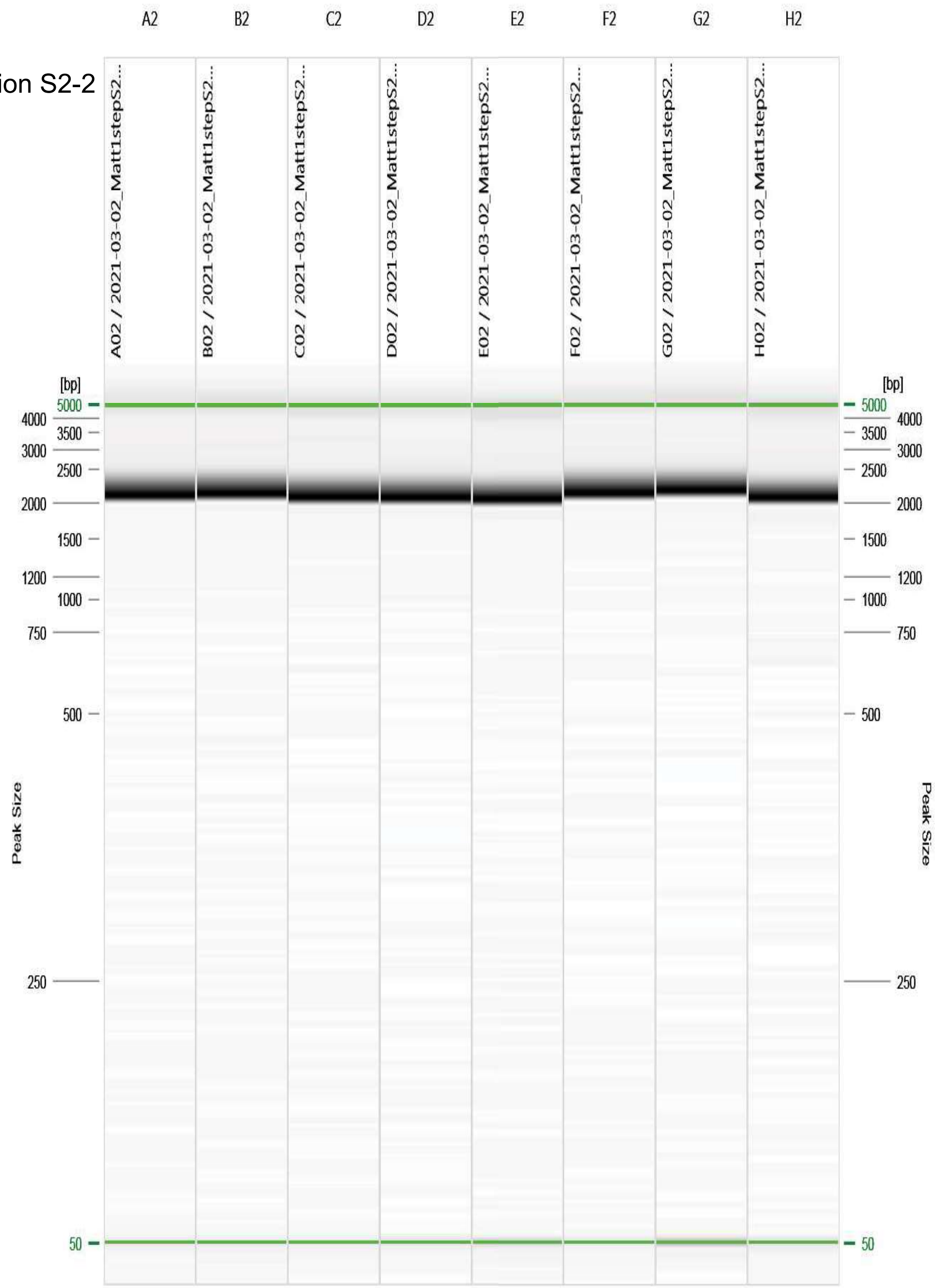


Figure: 2 - 2021-03-02_Matt1stepS2/R:1 E:1 #2

| | | | | | | | | |
|---|---|---|---|---|---|---|---|---|
| Annealing Temperature (°C) | 68.0 | 67.0 | 65.4 | 62.9 | 59.9 | 57.5 | 55.9 | 55.0 |

Primer Combination S2-3

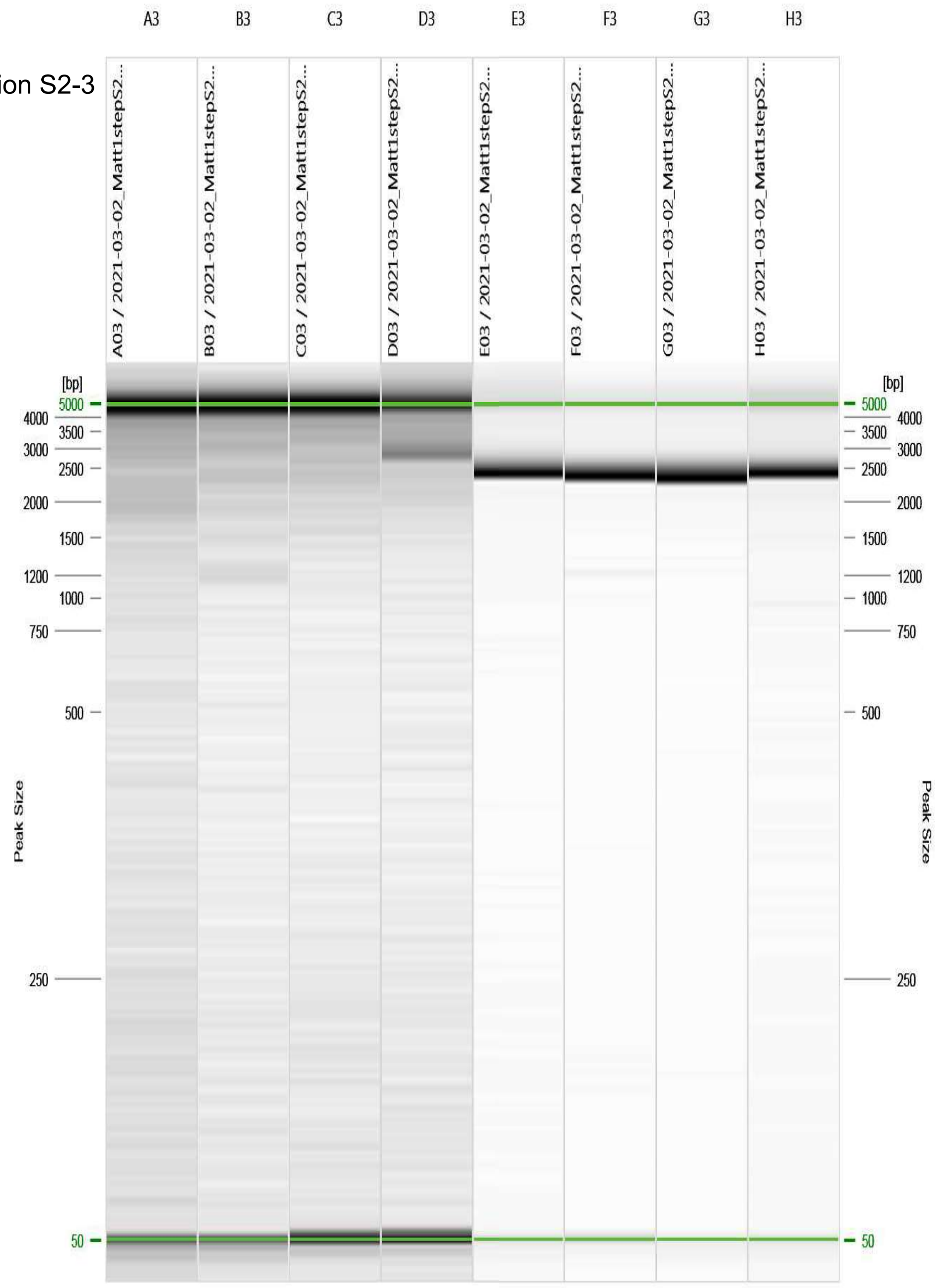


Figure: 3 - 2021-03-02_Matt1stepS2/R:1 E:1 #3

| Annealing Temperature (°C) | 68.0 | 67.0 | 65.4 | 62.9 | 59.9 | 57.5 | 55.9 | 55.0 |
|---|---|---|---|---|---|---|---|---|

Primer Combination S2-4

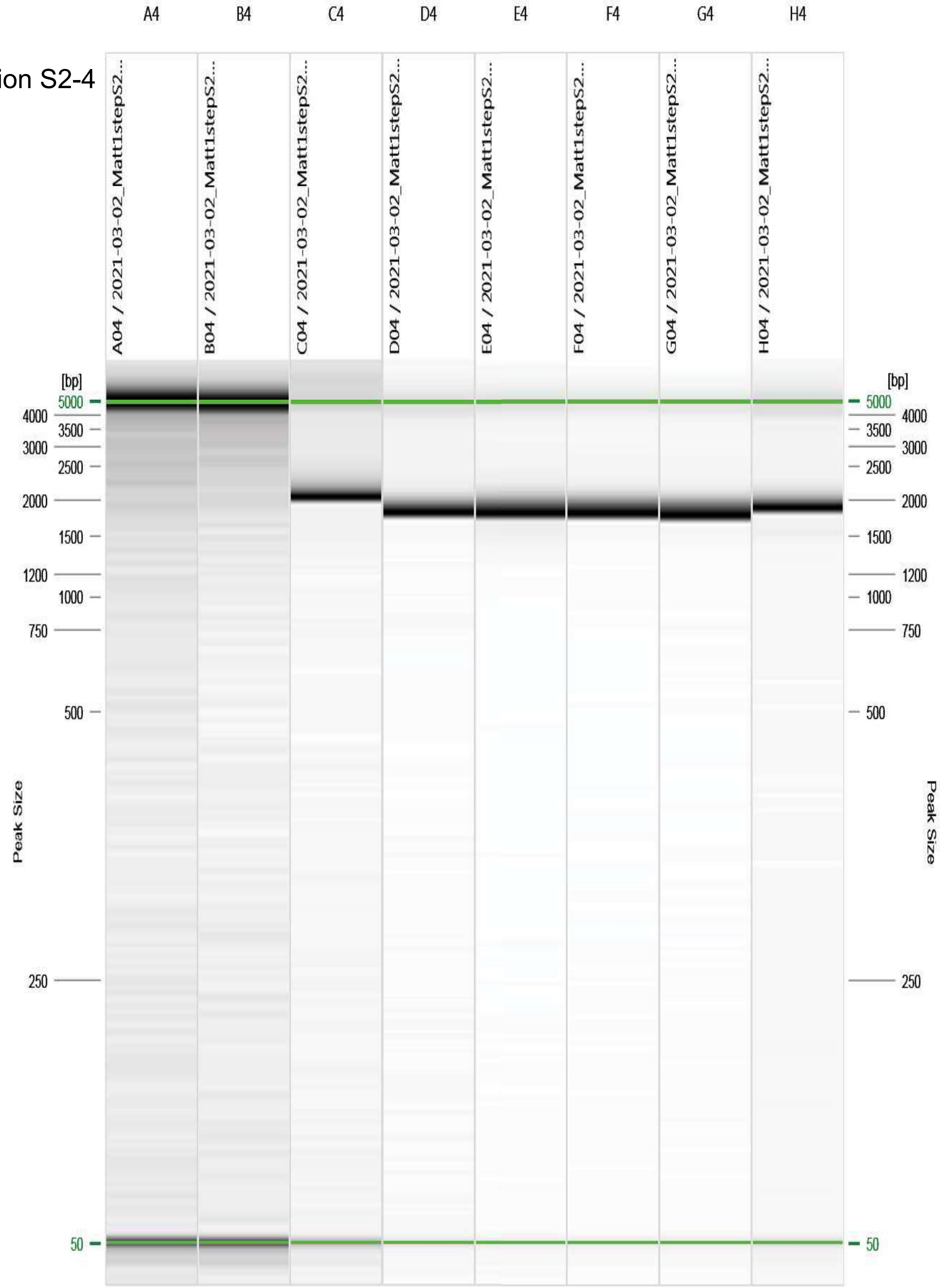


Figure: 4 - 2021-03-02_Matt1stepS2/R:1 E:1 #4

| Annealing Temperature (°C) | 68.0 | 67.0 | 65.4 | 62.9 | 59.9 | 57.5 | 55.9 | 55.0 |
|---|---|---|---|---|---|---|---|---|

Primer Combination S2-5

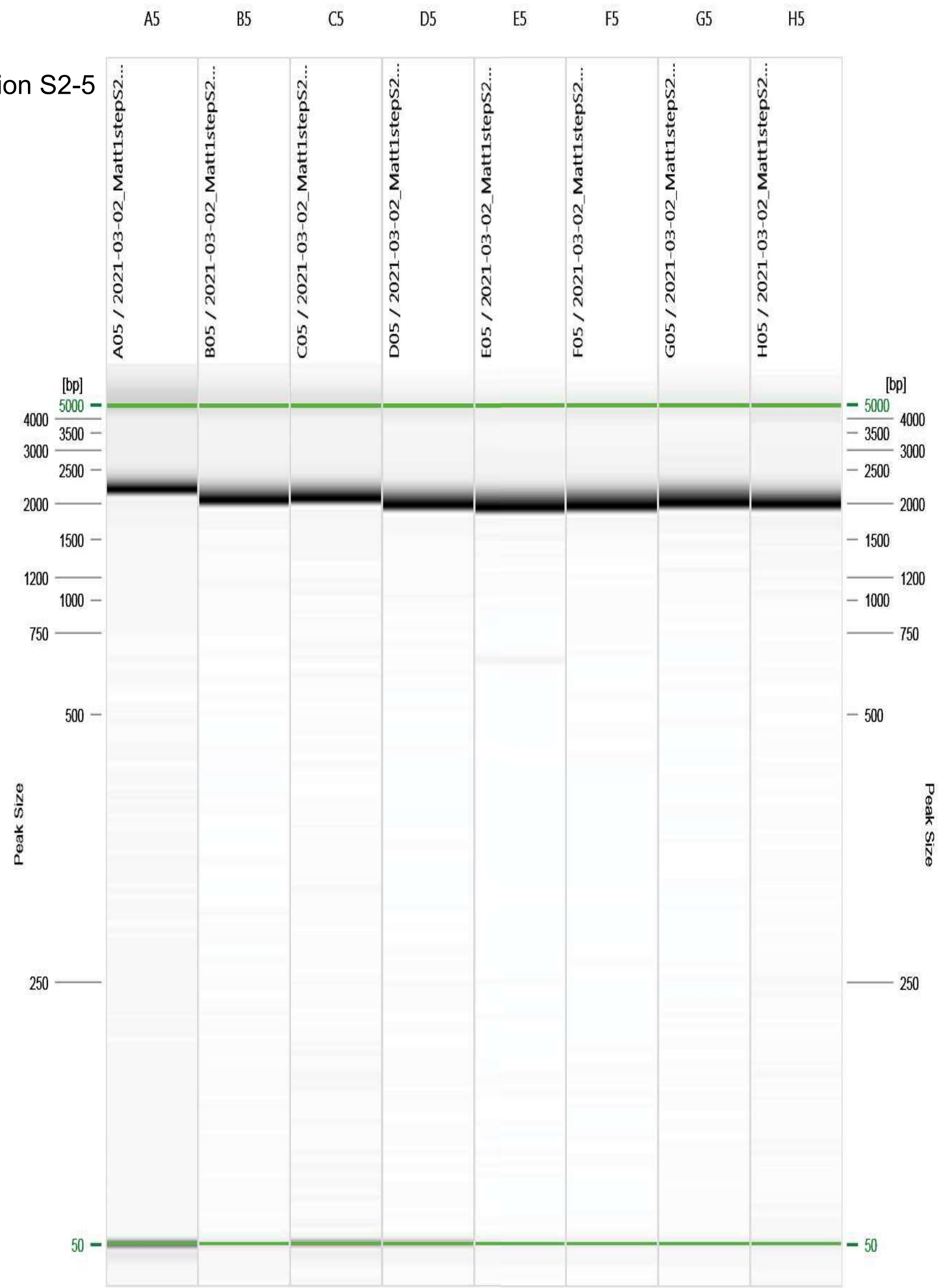


Figure: 5 - 2021-03-02_Matt1stepS2/R:1 E:1 #5

| Annealing Temperature (°C) | 68.0 | 67.0 | 65.4 | 62.9 | 59.9 | 57.5 | 55.9 | 55.0 |
|---|---|---|---|---|---|---|---|---|

Primer Combination S2-6

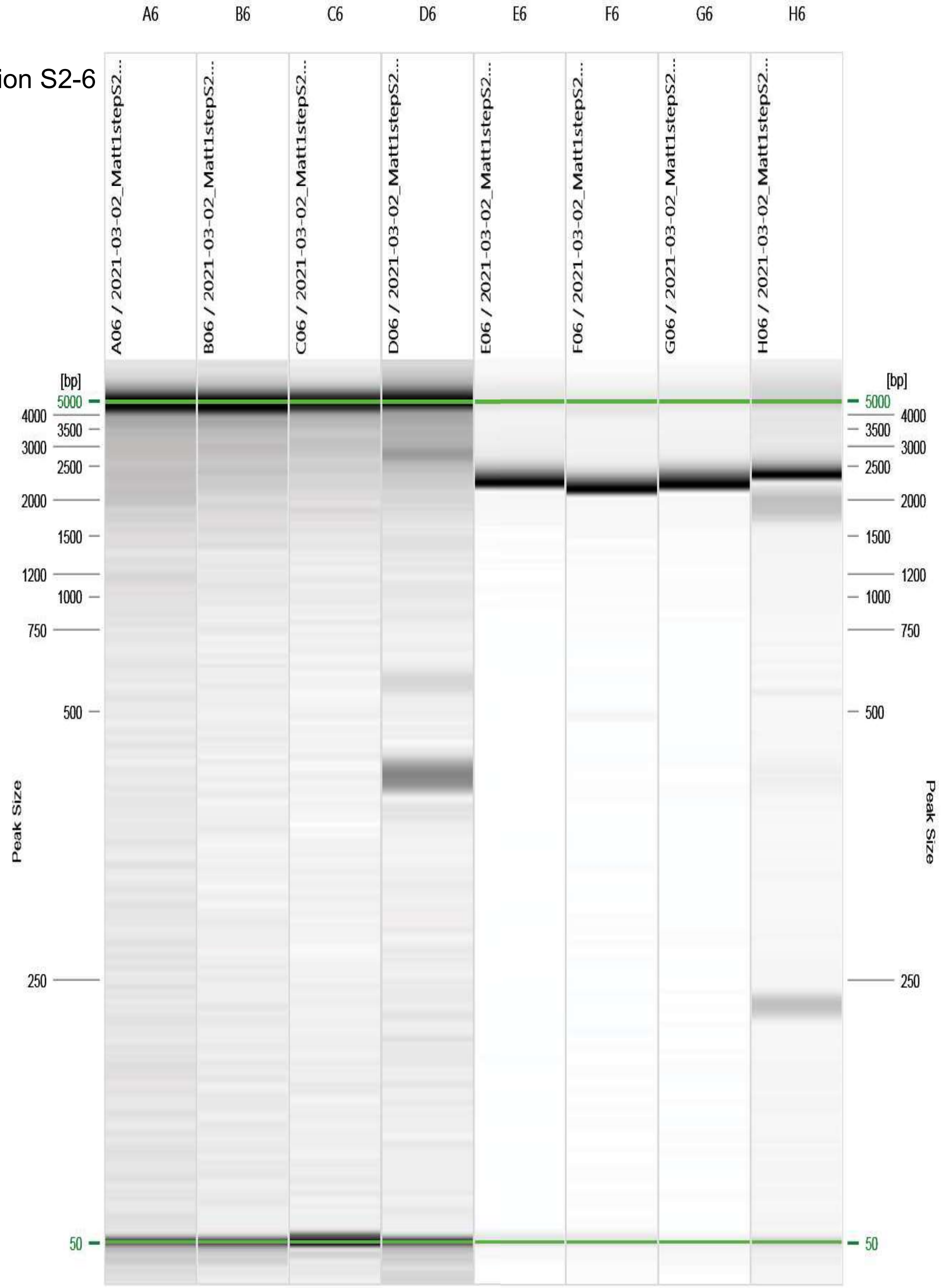


Figure: 6 - 2021-03-02_Matt1stepS2/R:1 E:1 #6

| Annealing Temperature (°C) | 68.0 | 67.0 | 65.4 | 62.9 | 59.9 | 57.5 | 55.9 | 55.0 |
|---|---|---|---|---|---|---|---|---|

Primer Combination S2-7

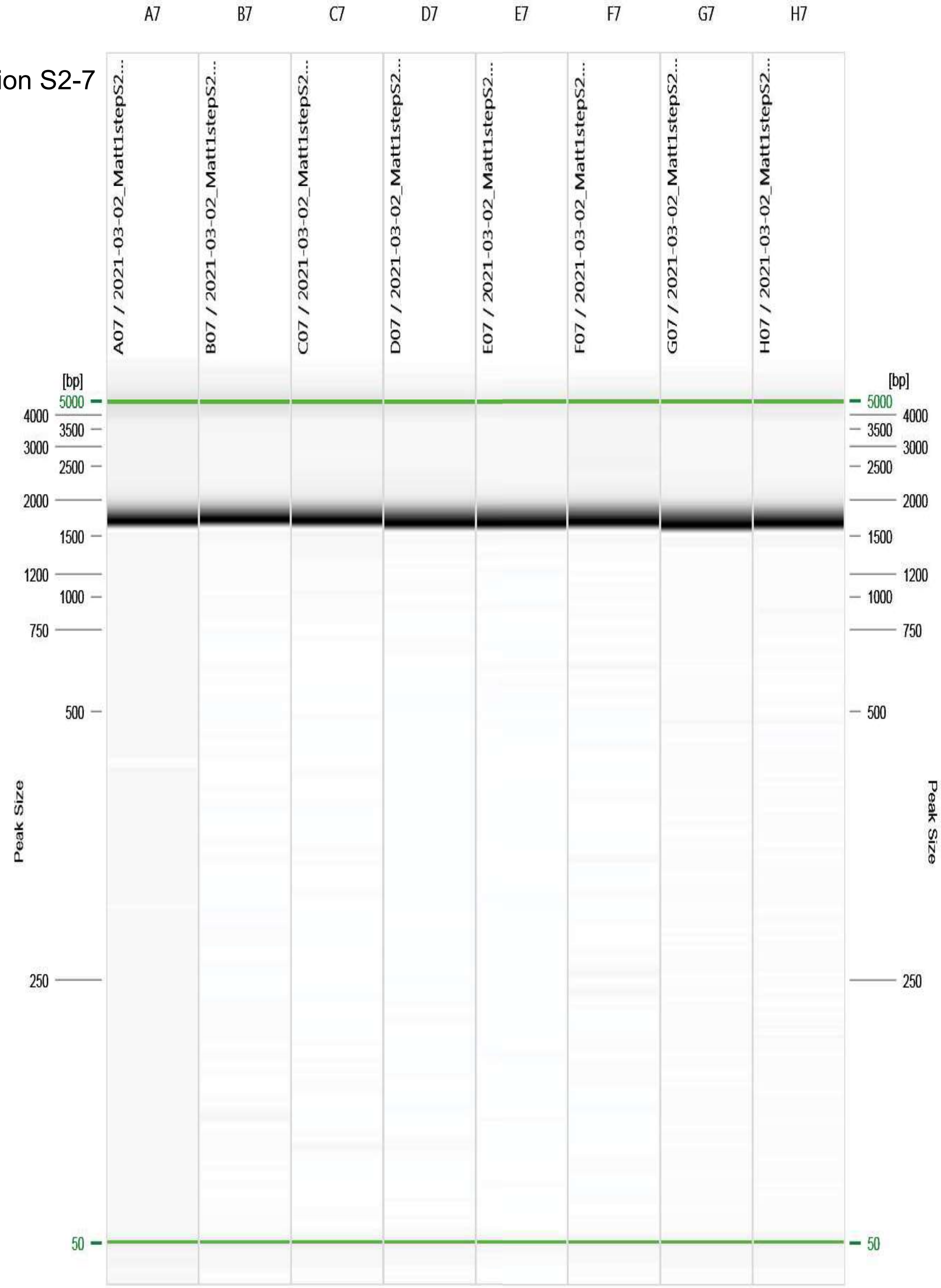


Figure: 7 - 2021-03-02_Matt1stepS2/R:1 E:1 #7

| Annealing Temperature (°C) | 68.0 | 67.0 | 65.4 | 62.9 | 59.9 | 57.5 | 55.9 | 55.0 |
|---|---|---|---|---|---|---|---|---|

Primer Combination S2-8

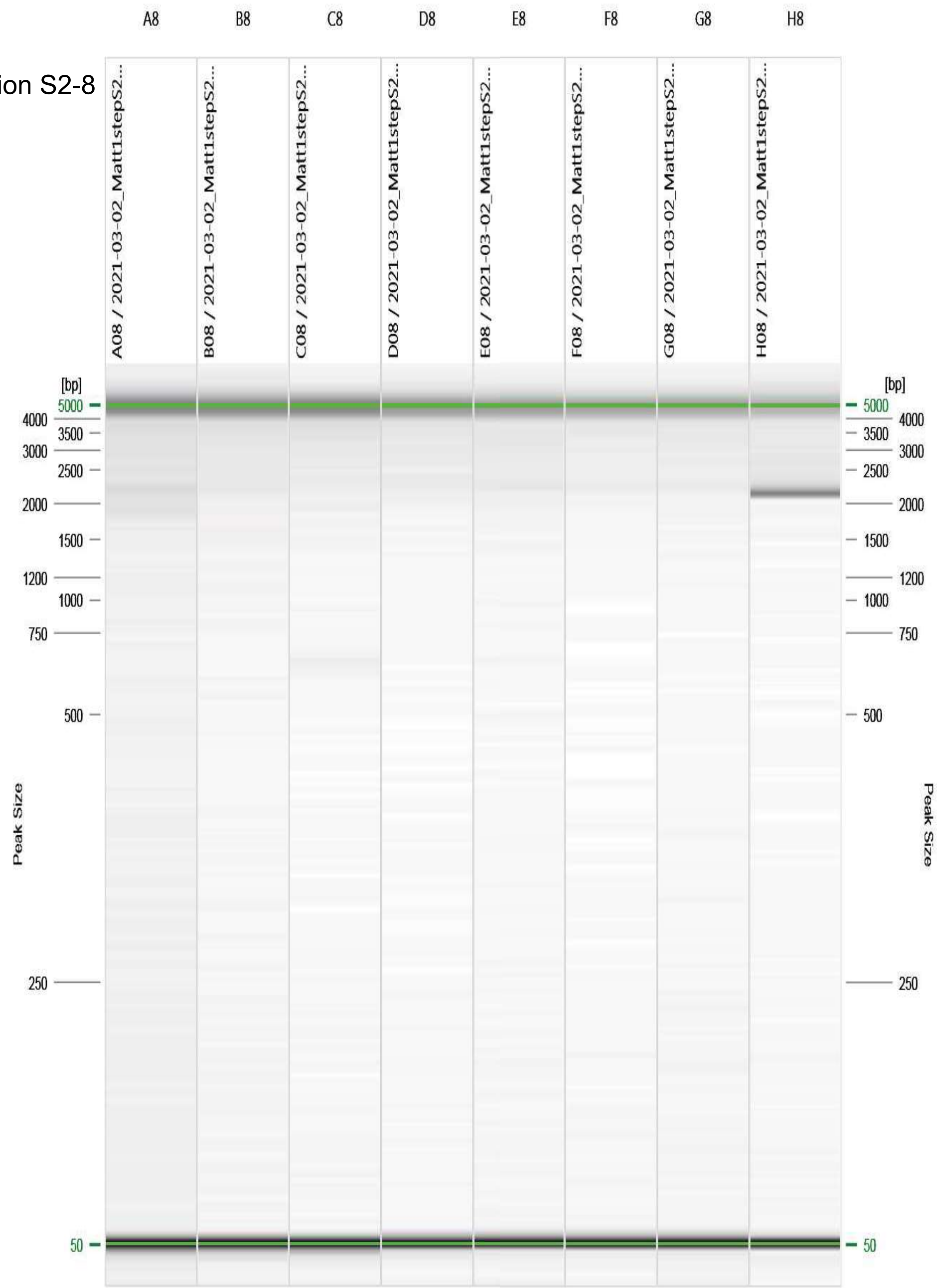


Figure: 8 - 2021-03-02_Matt1stepS2/R:1 E:1 #8

| Annealing Temperature (°C) | 68.0 | 67.0 | 65.4 | 62.9 | 59.9 | 57.5 | 55.9 | 55.0 |
|---|---|---|---|---|---|---|---|---|

Primer Combination S2-9

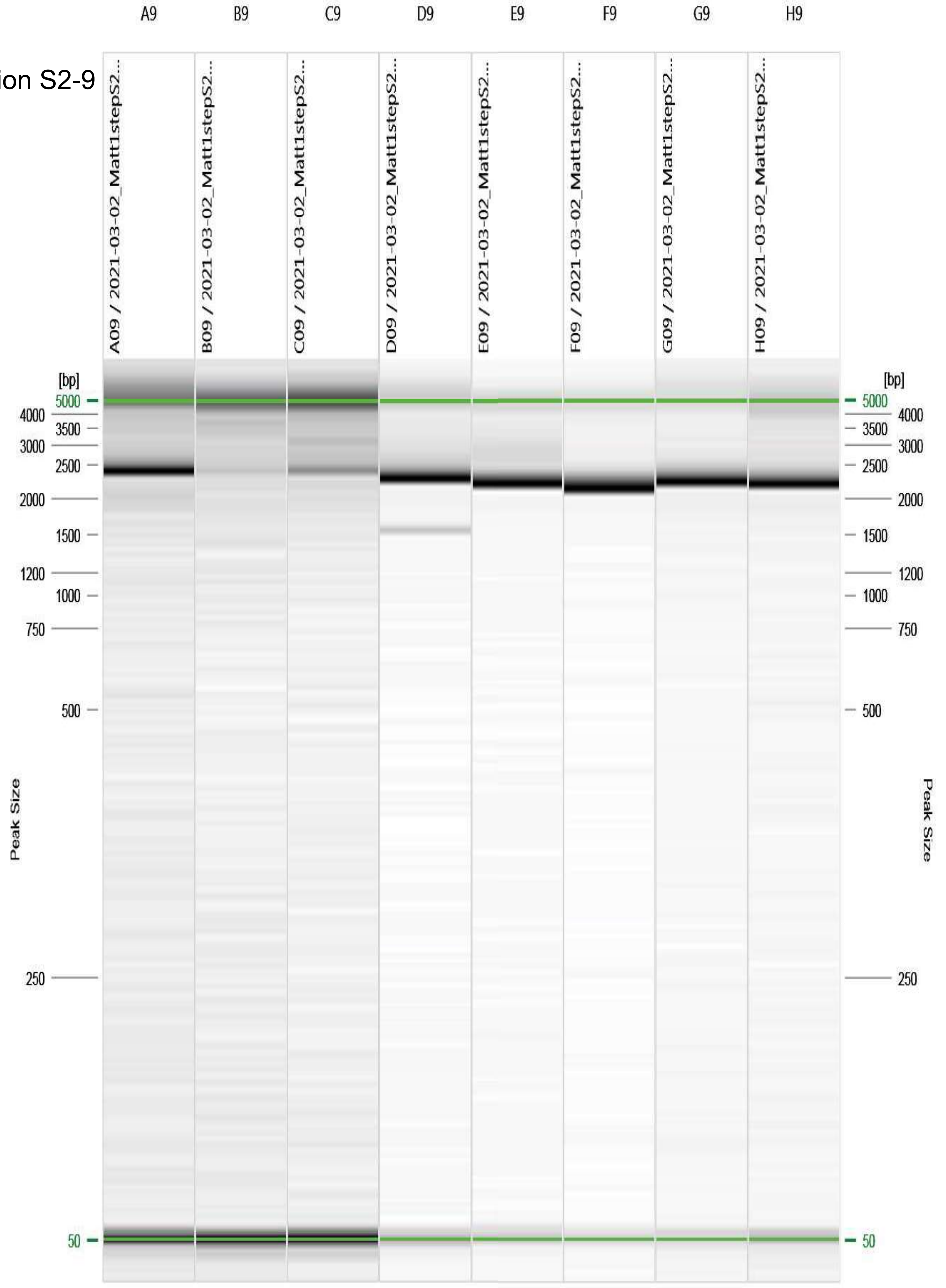


Figure: 9 - 2021-03-02_Matt1stepS2/R:1 E:1 #9
