## SupplementalMaterials for "Targeted Amplification and Genetic Sequencing of the Severe Acute Respiratory Syndrome Coronavirus 2 Surface Glycoprotein": Figure_S04_PrimerSelectionLOD_Fragments.pdf

| Report Overview | |
|---|---|
| Report Date: | 3/18/2021 3:32:01 PM |
| Experiment Name: | 2021-03-08_12PrimerLOD |
| Cartridge ID: | C191210A39 |
| Instrument ID: | 30334 |

| SARS-CoV-2 Abundance | ct14 | ct16 | ct17 | ct20 | ct22 | ct25 | ct27 | ct30 |
|---|---|---|---|---|---|---|---|---|

Primer Combination S1-1

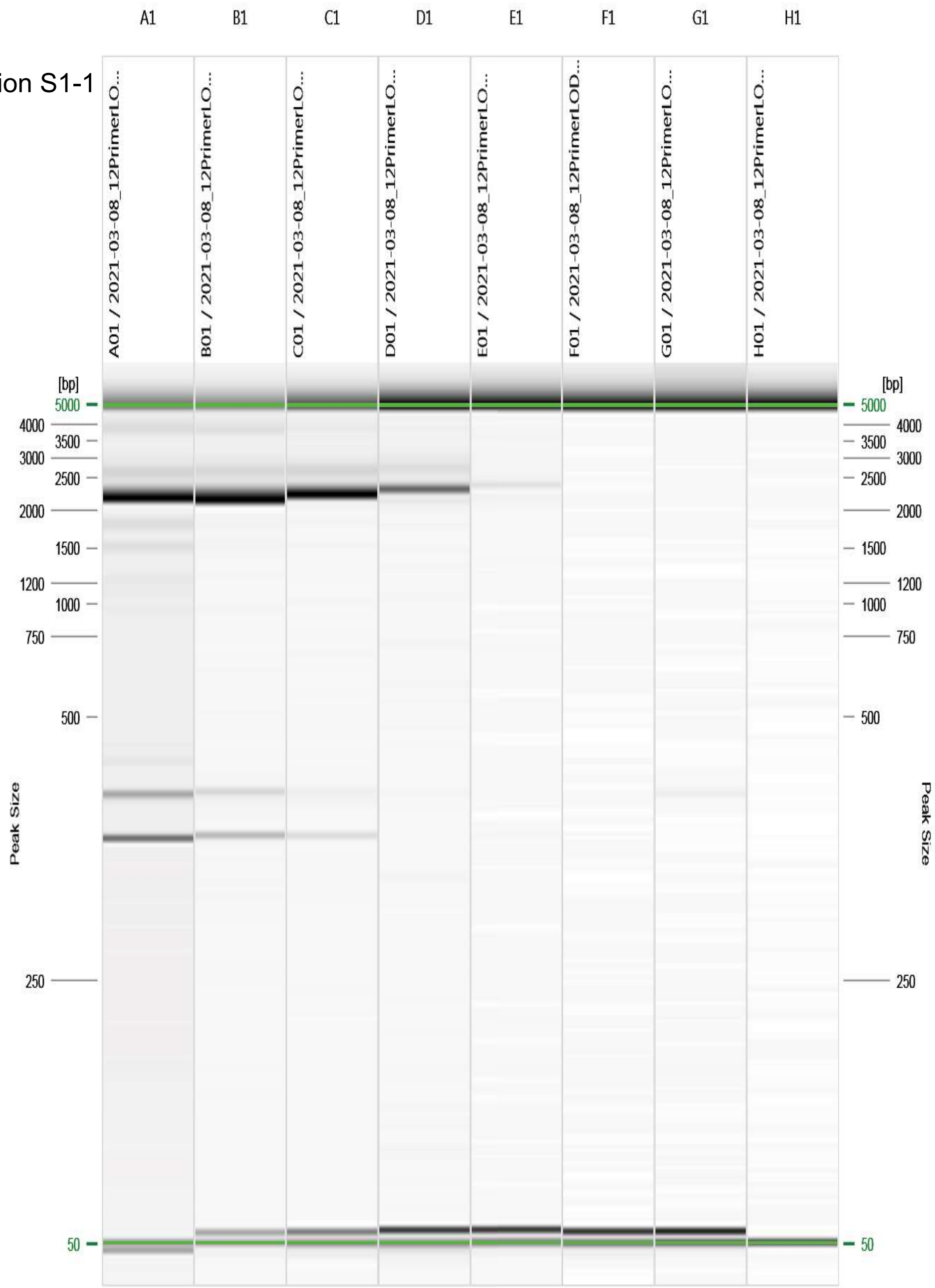


Figure: 1 - 2021-03-08_12PrimerLOD/R:1 E:1 #1

| SARS-CoV-2 Abundance | ct14 | ct16 | ct17 | ct20 | ct22 | ct25 | ct27 | ct30 |
|---|---|---|---|---|---|---|---|---|

Primer Combination S1-2

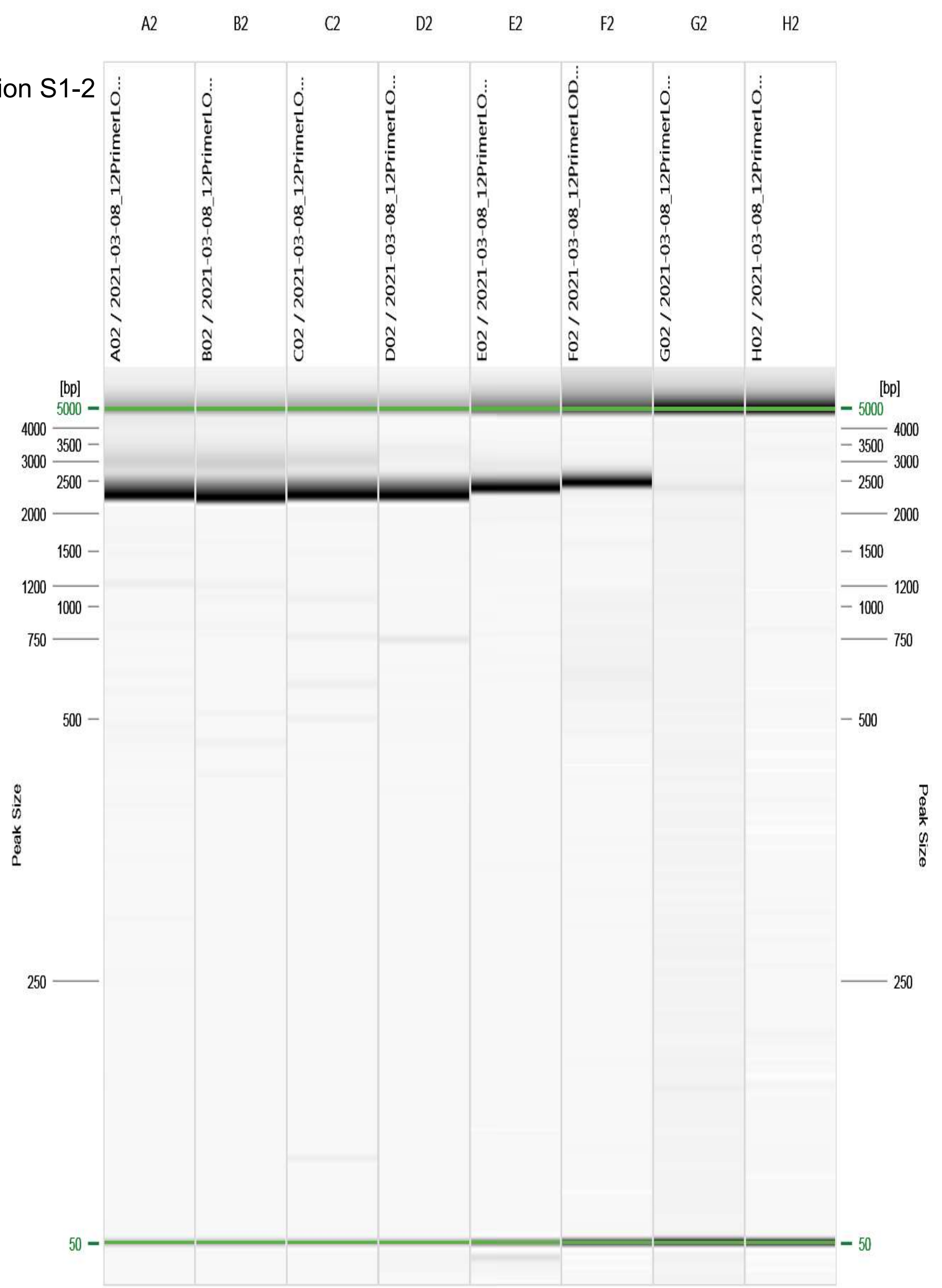


Figure: 2 - 2021-03-08_12PrimerLOD/R:1 E:1 #2

| SARS-CoV-2 Abundance | ct14 | ct16 | ct17 | ct20 | ct22 | ct25 | ct27 | ct30 |
|---|---|---|---|---|---|---|---|---|

Primer Combination S1-3

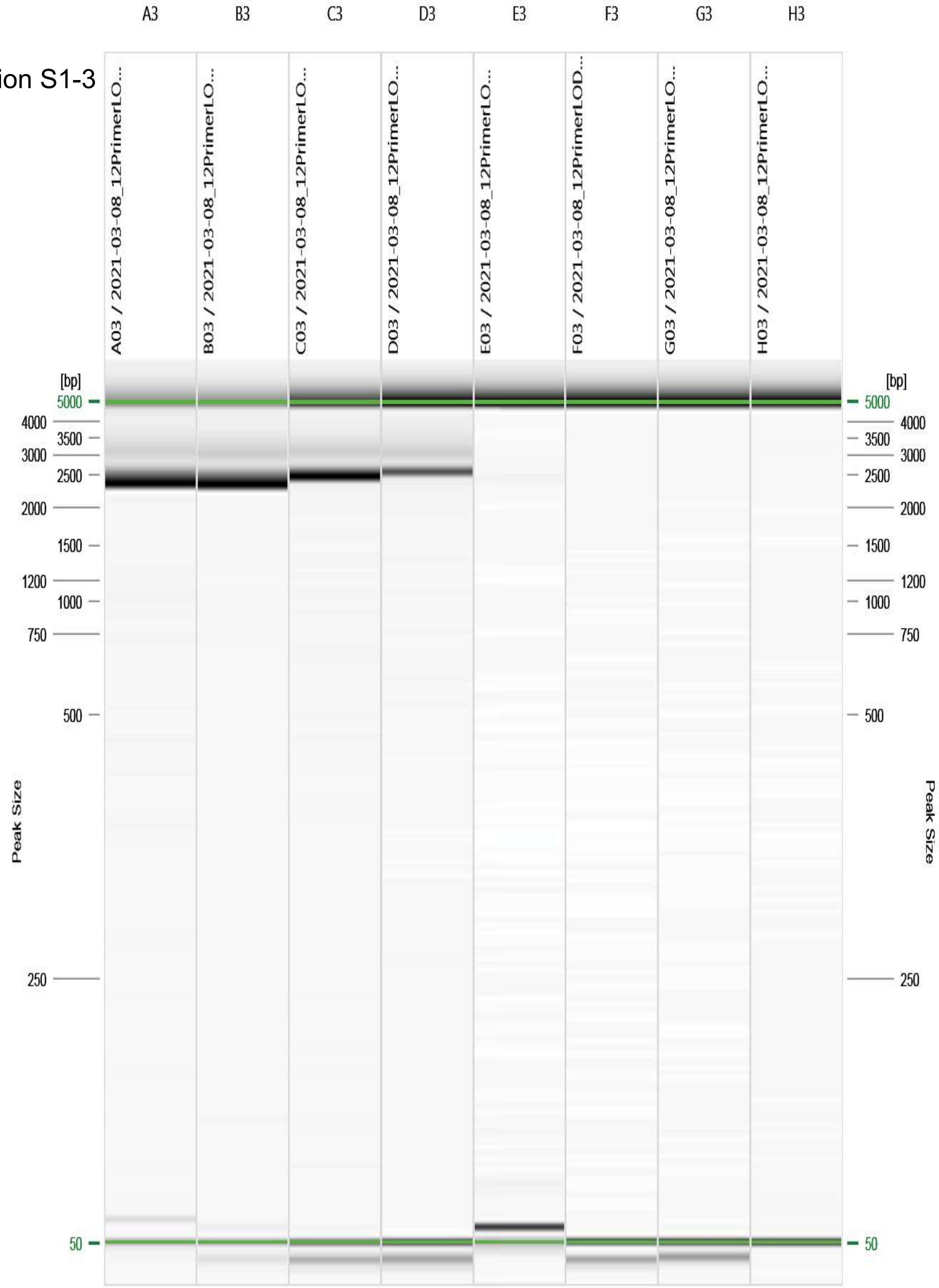


Figure: 3 - 2021-03-08_12PrimerLOD/R:1 E:1 #3

| SARS-CoV-2 Abundance | ct14 | ct16 | ct17 | ct20 | ct22 | ct25 | ct27 | ct30 |
|---|---|---|---|---|---|---|---|---|

Primer Combination S1-8

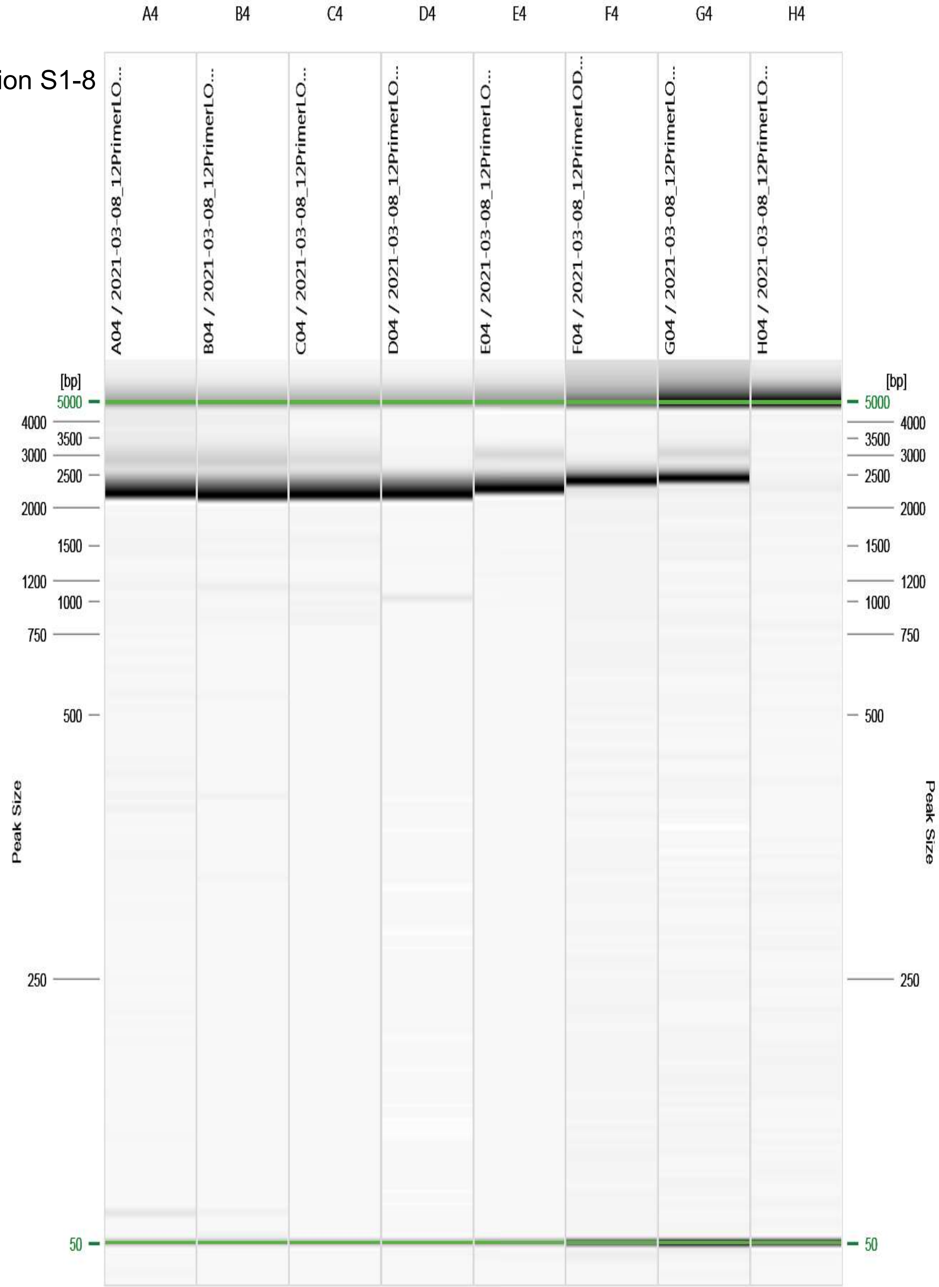


Figure: 4 - 2021-03-08_12PrimerLOD/R:1 E:1 #4

| SARS-CoV-2 Abundance | ct14 | ct16 | ct17 | ct20 | ct22 | ct25 | ct27 | ct30 |
|---|---|---|---|---|---|---|---|---|

Primer Combination S1-9

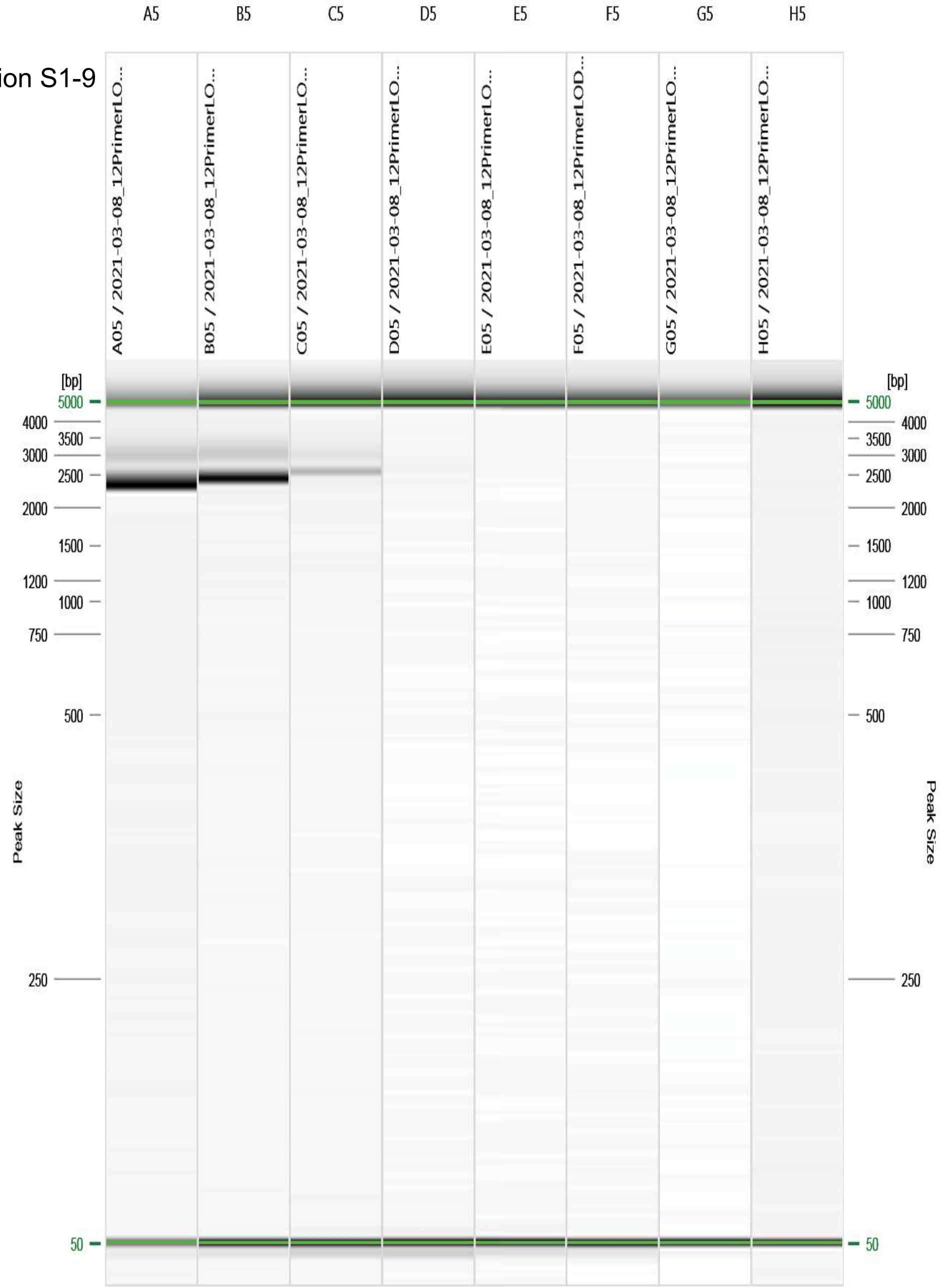


Figure: 5 - 2021-03-08_12PrimerLOD/R:1 E:1 #5

| SARS-CoV-2 Abundance | ct14 | ct16 | ct17 | ct20 | ct22 | ct25 | ct27 | ct30 |
|---|---|---|---|---|---|---|---|---|

Primer Combination S2-1

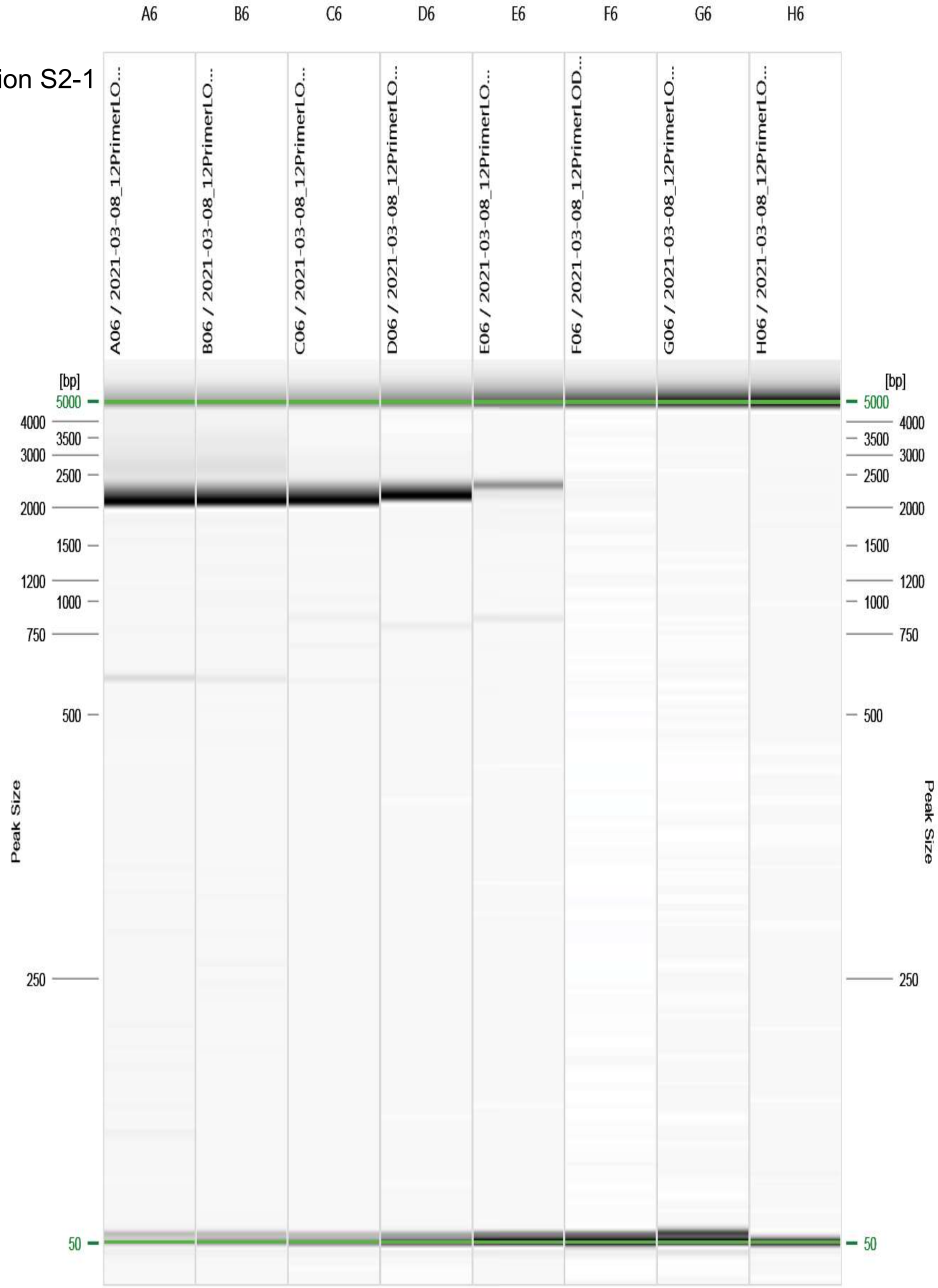


Figure: 6 - 2021-03-08_12PrimerLOD/R:1 E:1 #6

| SARS-CoV-2 Abundance | ct14 | ct16 | ct17 | ct20 | ct22 | ct25 | ct27 | ct30 |
|---|---|---|---|---|---|---|---|---|

Primer Combination S2-2

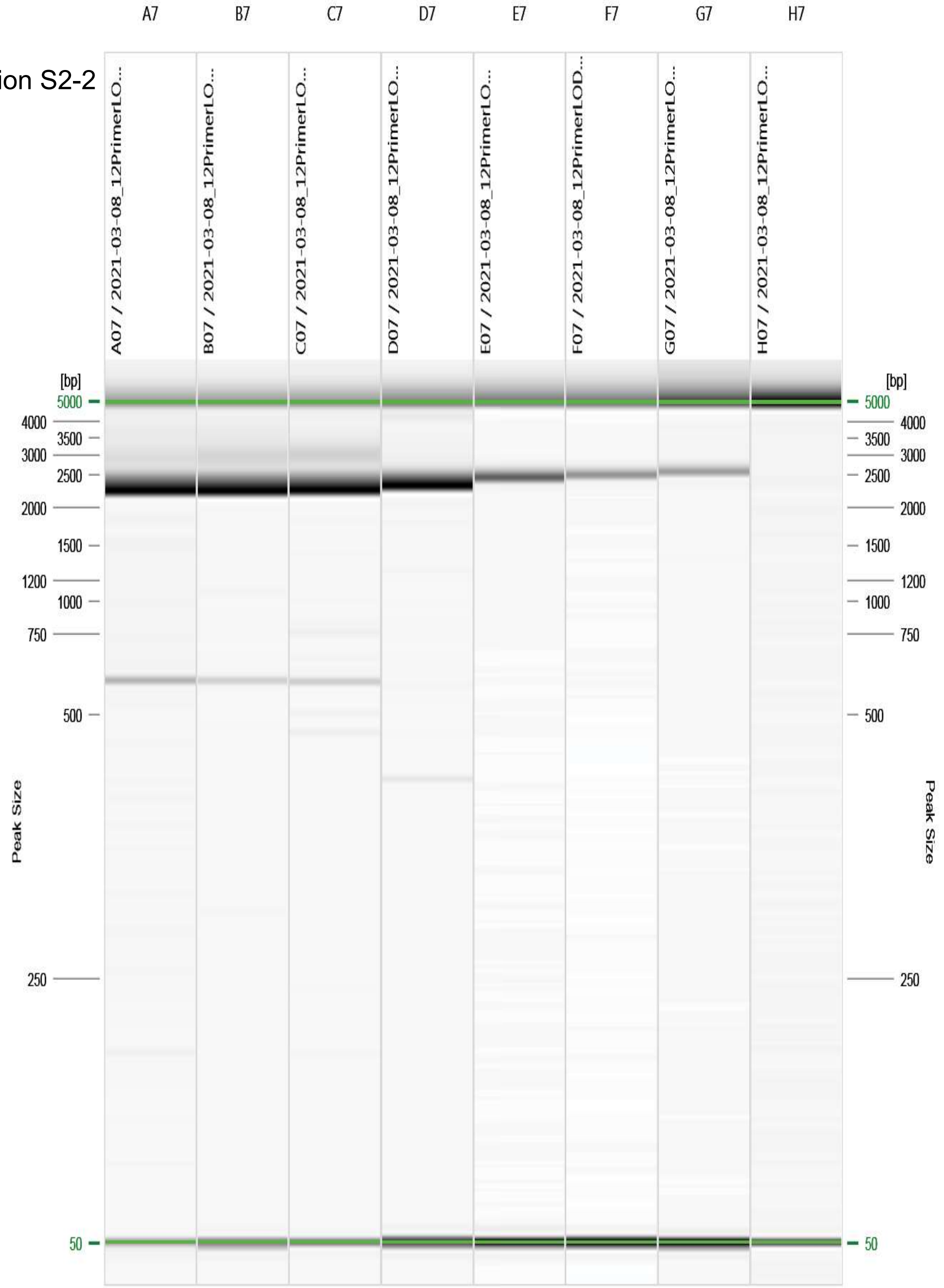


Figure: 7 - 2021-03-08_12PrimerLOD/R:1 E:1 #7

| SARS-CoV-2 Abundance | ct14 | ct16 | ct17 | ct20 | ct22 | ct25 | ct27 | ct30 |
|---|---|---|---|---|---|---|---|---|

Primer Combination S2-4

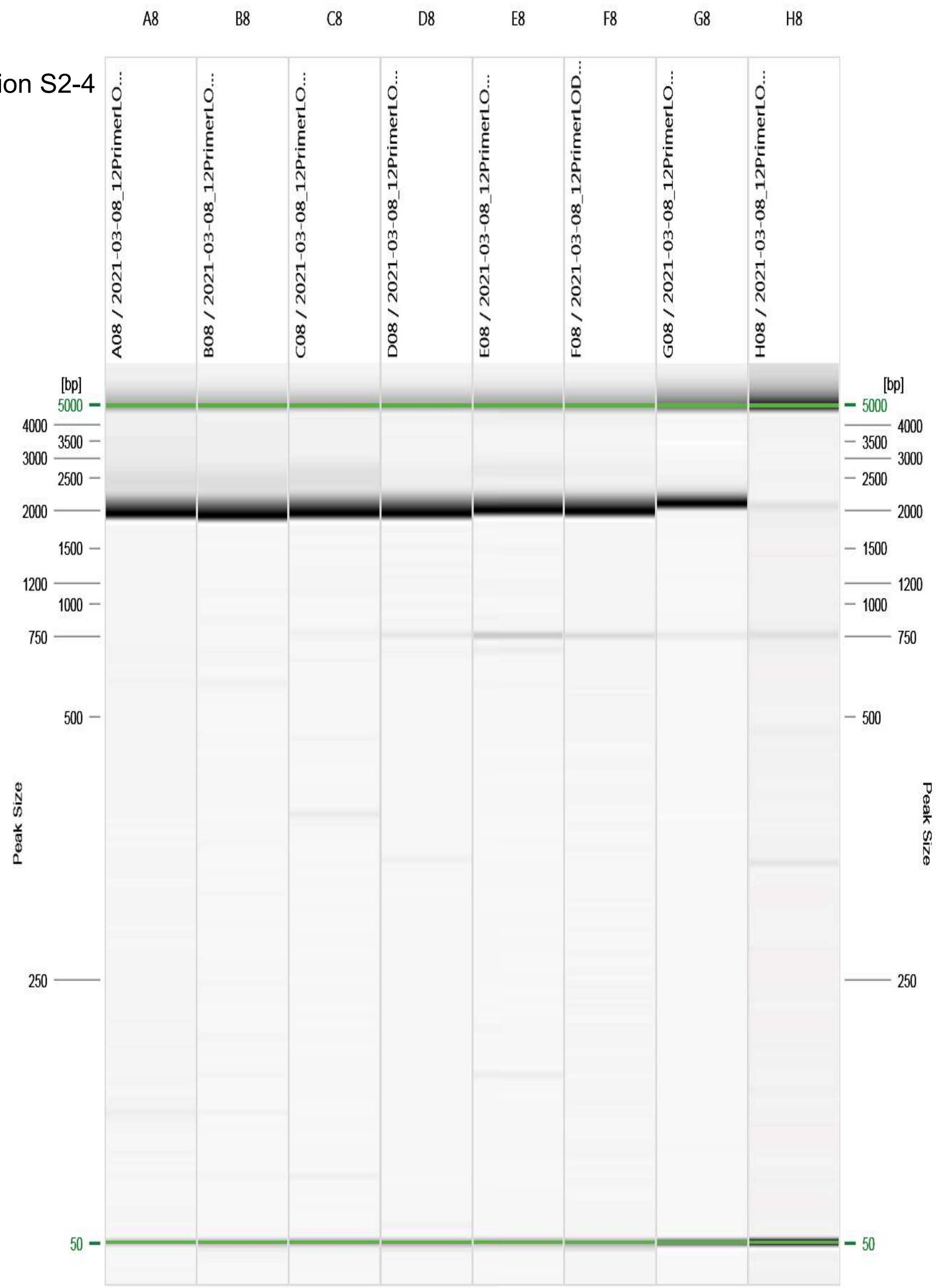


Figure: 8 - 2021-03-08_12PrimerLOD/R:1 E:1 #8

| SARS-CoV-2 Abundance | ct14 | ct16 | ct17 | ct20 | ct22 | ct25 | ct27 | ct30 |
|---|---|---|---|---|---|---|---|---|

Primer Combination S2-5

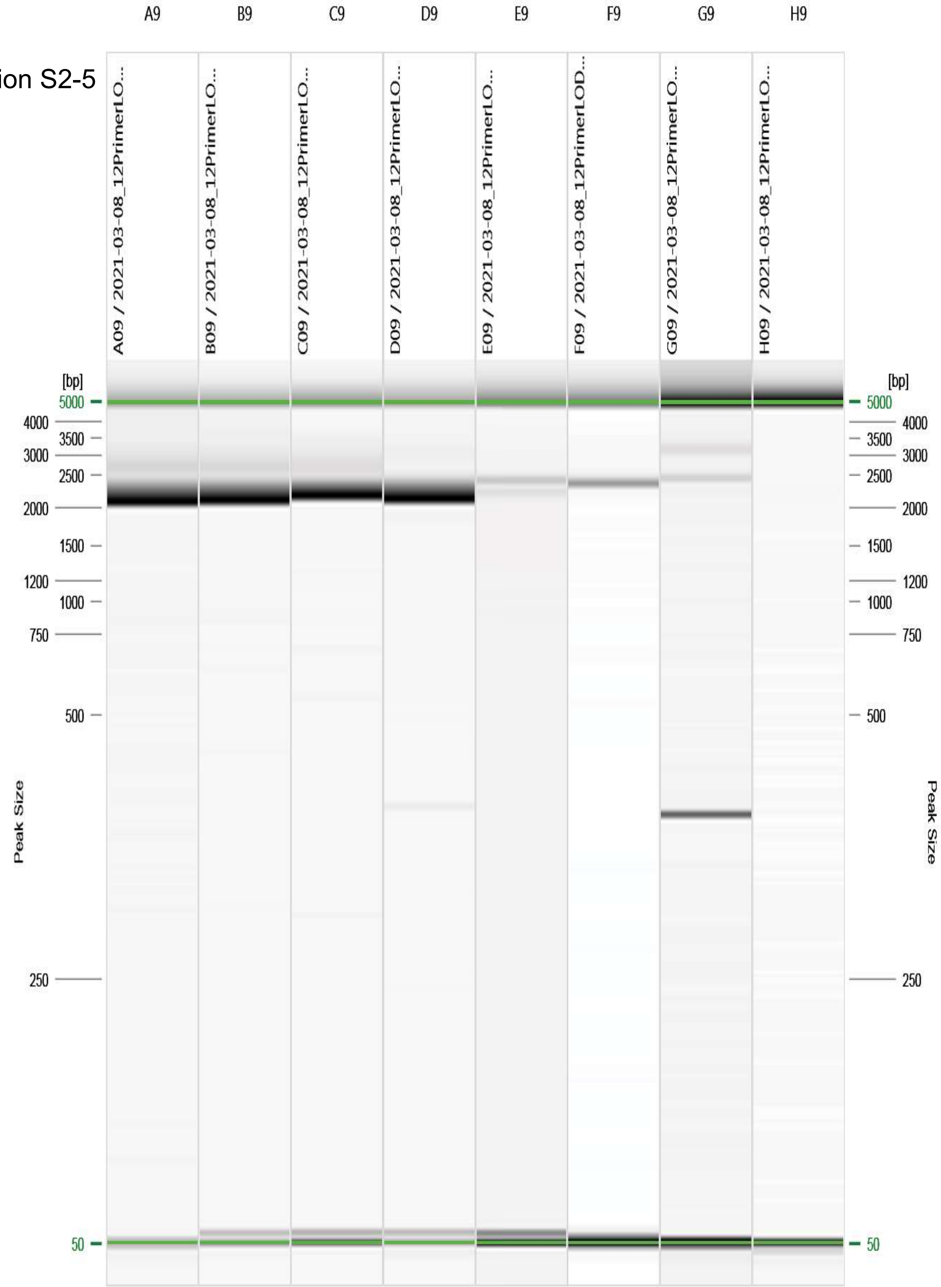


Figure: 9 - 2021-03-08_12PrimerLOD/R:1 E:1 #9

| SARS-CoV-2 Abundance | ct14 | ct16 | ct17 | ct20 | ct22 | ct25 | ct27 | ct30 |
|---|---|---|---|---|---|---|---|---|

Primer Combination S2-6

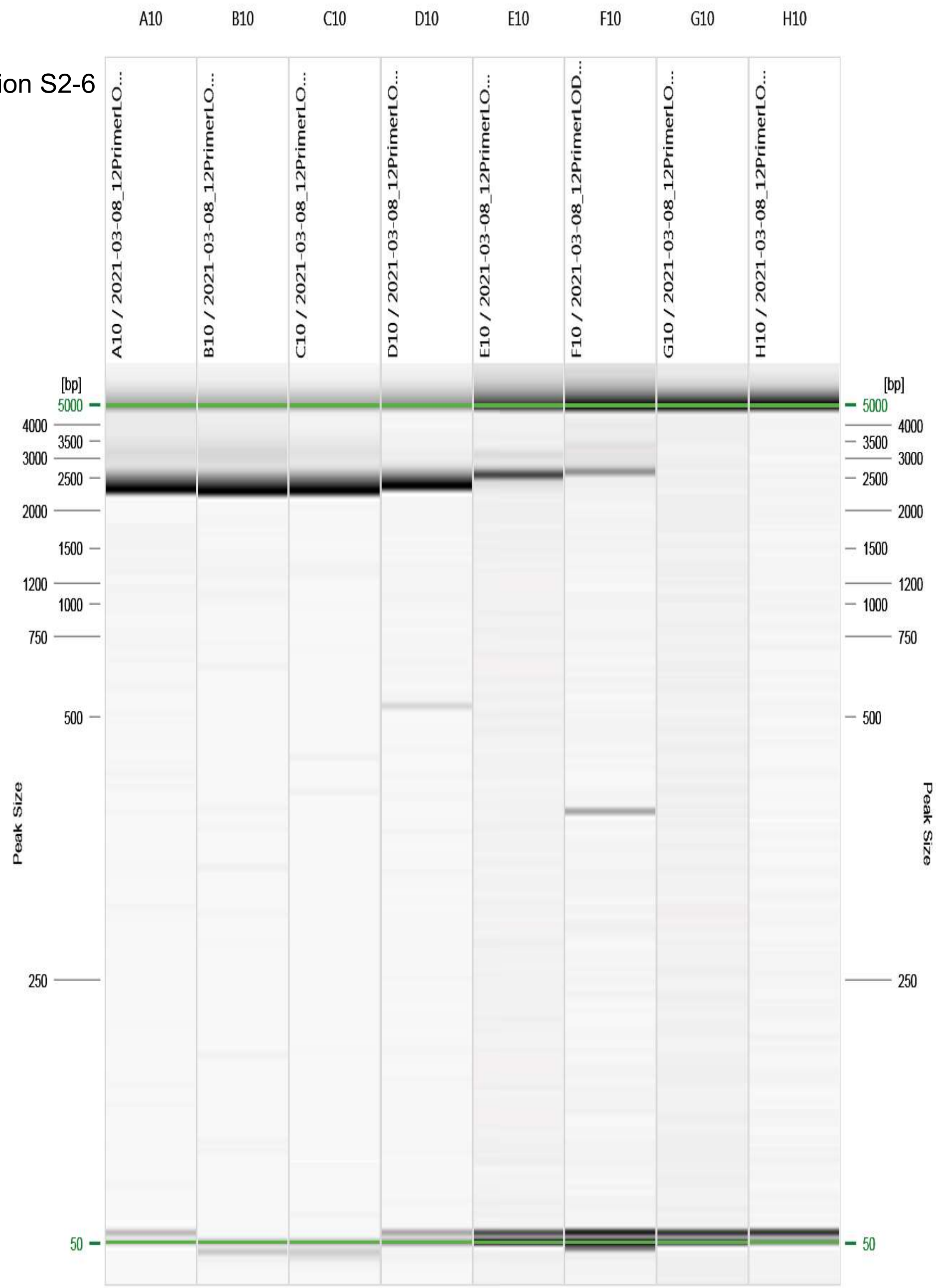


Figure: 10 - 2021-03-08_12PrimerLOD/R:1 E:1 #10

| SARS-CoV-2 Abundance | ct14 | ct16 | ct17 | ct20 | ct22 | ct25 | ct27 | ct30 |
|---|---|---|---|---|---|---|---|---|

Primer Combination S2-7

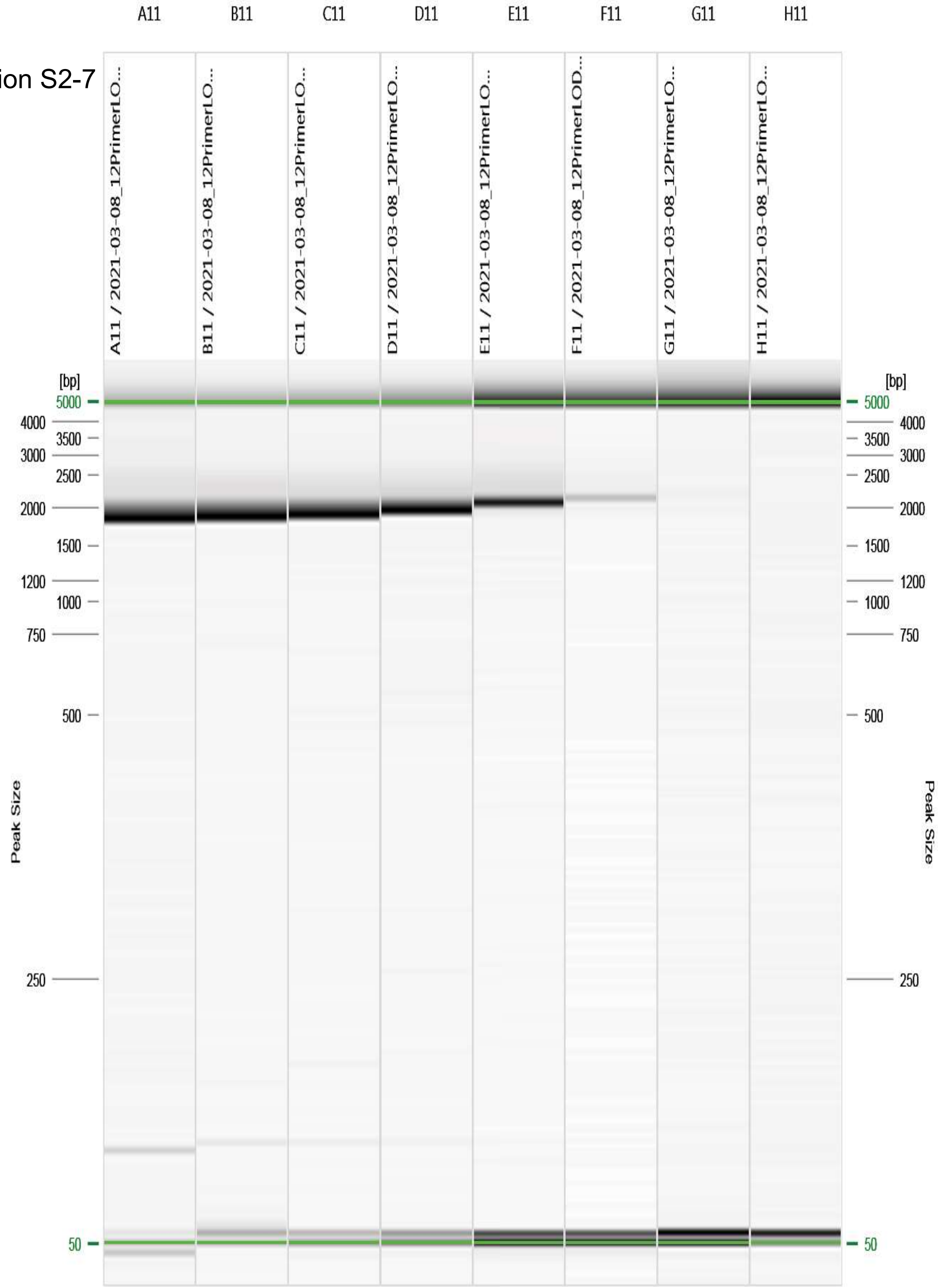


Figure: 11 - 2021-03-08_12PrimerLOD/R:1 E:1 #11

| SARS-CoV-2 Abundance | ct14 | ct16 | ct17 | ct20 | ct22 | ct25 | ct27 | ct30 |
|---|---|---|---|---|---|---|---|---|

Primer Combination S2-9

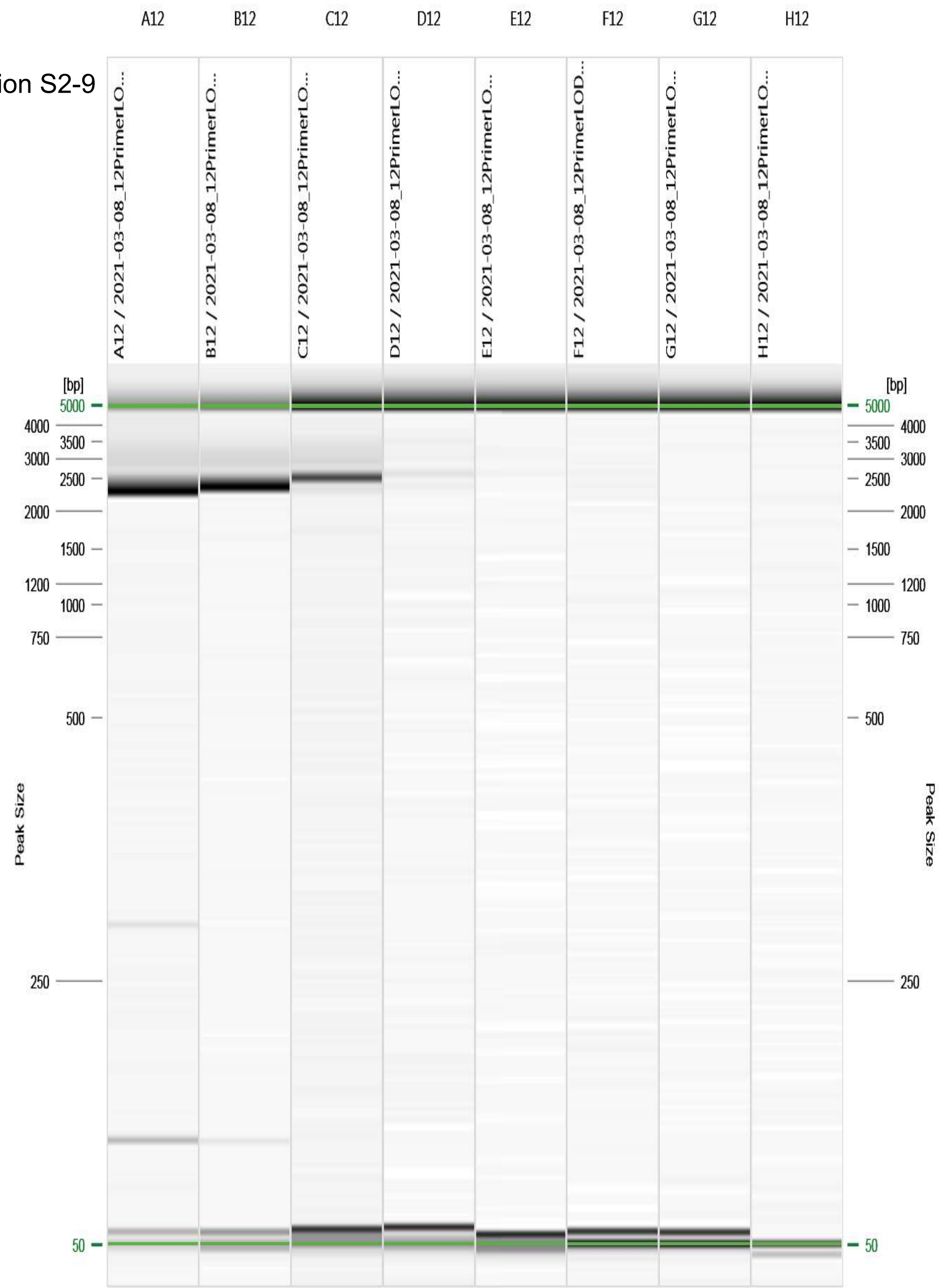


Figure: 12 - 2021-03-08_12PrimerLOD/R:1 E:1 #12
