## SupplementalMaterials for "Targeted Amplification and Genetic Sequencing of the Severe Acute Respiratory Syndrome Coronavirus 2 Surface Glycoprotein": Text_S01_LP-471 - RT-PCR of the SARS-CoV-2 S-gene for Sequencing.pdf

### Reverse Transcription-PCR (RT-PCR) of the SARS-CoV-2 S-gene for Sequencing

*Virology, Surveillance and Diagnosis Branch, Genomics and Diagnostics Team (GDT)*

**NOTE: This procedure is provided for research use only. This document is not intended to be used for commercial development or for-profit testing. Use of trade names and commercial sources is for identification only and does not constitute endorsement by the Public Health Service or by the United States Department of Health and Human Services. Please do not distribute this document to other laboratories or commercial entities.**

#### 1.0 Purpose

1.1 The purpose of this procedure is to describe a fast double-reaction amplification of the SARS-CoV-2 S-gene for subsequent sequencing. Full coverage of the S-gene requires two overlapping amplicons to be produced via separate RT-PCR reactions.

#### 2.0 Definitions

2.1 RT-PCR: Reverse Transcription Polymerase Chain Reaction.

#### 3.0 Equipment (Use Examples or Equivalent)

3.1 Sterile, nuclease-free 1.5 mL micro-centrifuge tubes

3.2 0.2 mL PCR reaction tube strips or plates

3.2.1 PCR 8-tube strips (Brand Tech Scientific Inc. Catalog. No. 781332)

3.2.2 PCR Plate, 96-well, semi-skirted, flat deck (Life Technologies, Catalog No. AB-1400)

3.2.3 TempPlate pierceable sealing foil, sterile (USA Scientific: Catalog No. 2923-0110)

3.2.4 Sealing Roller (BIO-RAD: Catalog No. MSR-0001)

3.2.5 Silicone compression mat (Sigma Aldrich: Catalog No. AXYCMFLAT)

3.3 Vortex

3.4 1.5 mL tube, 0.2 mL strip tube, and 96 well plate compatible centrifuge

3.5 Cold blocks for 0.2 mL and 1.5 mL PCR reaction tubes (ISC BioExpress)

3.6 Pipettes (10 µL, 20 µL, 200 µL, and 1000 µL)

3.6.1 Multichannel pipettes also recommended (20 µL, and 200 µL)

3.6.2 Corresponding aerosol barrier pipette tips

3.7 Disposable reagent reservoir, sterile (Axygen: Catalog No. RES-V-25-S)

3.8 96-well format PCR Thermocycler System (BioRad: T100)

3.9 1.5 mL tube, 8-tube strip, or 96-well format magnetic separation rack (Life Technologies DynaMag-2: Catalog No. 12321D, Alpaqua Magnum FLX: Catalog No. A000400)

3.10 DNA electrophoresis and visualization equipment (QIAxcel advanced: Catalog No. 9001941)

#### 4.0 Reagents

4.1 Nuclease-free water

4.2 SuperScript™ IV One-Step RT-PCR System (Invitrogen: Catalog. No. 12594100 – 100 reactions)

4.3 SPRI beads (Beckman Coulter: Catalog No. A63880, A63881, A63882, or equivalent)

4.4 Molecular biology grade absolute ethanol

#### 5.0 Primers

5.1 SARS-CoV-2 S-gene primers listed in Table 1 are from Integrated DNA Technologies Inc. (IDT) https://www.idtdna.com (or equivalent). Primers must be RNase Free HPLC purified.

5.1.1 Prepare 10 µM stocks of each individual primer.

5.1.2 Pool the 10 µM forward and reverse primers in a 1:1 ratio for both the S1 and S2 primer pools.

5.2 CDC provided primers are premixed at 1x strength and dried.

5.2.1 Reconstitute in 1 mL nuclease-free water.

| **Table 1: S1 and S2 primer pools** | | | |
|---|---|---|---|
| **S1 primer pool** | | | |
| **Oligo** | **# Bases** | **Sequence 5'-3'** | **µM in pool** |
| S1F_21358 | 29 | ACAAATCCAATTCAGTTGTCTTCCTATTC | 5 |
| S1R_23813 | 22 | TGCTGCATTCAGTTGAATCACC | 5 |
| **S2 primer pool** | | | |
| **Oligo** | **# Bases** | **Sequence 5’-3’** | **µM in pool** |
| S2F_23288 | 21 | GTCCGTGATCCACAGACACTT | 5 |
| S2R_25460 | 24 | GCATCCTTGATTTCACCTTGCTTC | 5 |

#### 6.0 Positive and Negative Controls

**Table 2: Positive and Negative Controls**

| Control | Material | Frequency | Expected Outcome |
|---|---|---|---|
| Positive | Twist Bioscience Custom NGS Spike Control<br>e.g. Delta REF: 103885; Omicron REF: 103885<br>Diluted to 20K copies/µL<br>Or<br>Previously sequenced RNA from propagated isolate of currently circulating SARS-CoV-2 variant, Ct <20<br>Or<br>Previously sequenced RNA from clinical sample of currently circulating SARS-CoV-2 variant, Ct <20 | Every run | Amplicons detectable by electrophoresis |
| Negative | Water | Every run | Amplicons not detectable by electrophoresis |

#### 7.0 Safety Precautions

7.1 Personal Protective Equipment (PPE) - Wear lab coats, gloves, and safety glasses.

7.2 Positive controls should be handled in an approved BSL-2 handling area to avoid contamination of laboratory equipment and reagents that could cause false positive results.

7.3 Materials should be handled in accordance with Good Laboratory Practices.

#### 8.0 Comments and Questions

#### 9.0 Gather Reagents

9.1 Thaw and store at the indicated temperature during the procedure.

9.2 Flick/invert the reagent tubes to ensure they are well mixed, and spin down before opening.

9.3 **Keep enzymes at -20°C**

9.3.1 SuperScript IV RT Mix

9.4 **+4°C**

9.4.1 SSIV 2X Reaction Mix

9.4.2 S1 and S2 Primer Pool

9.4.3 Samples and Controls

9.5 **Room temperature (15-25°C).**

9.5.1 Nuclease-free water

9.5.2 SPRI beads

9.5.3 Ethanol

#### 10.0 RT-PCR Procedure

10.1 Combine the components of Table 3 to prepare two reaction master mixes, one for each S1 and S2 reaction, sufficient for all samples and controls.

| Table 3: RT-PCR master mix for each SARS-CoV-2 amplicon | | |
|---|---|---|
| **Reagent** | **Volume (µL) Per Reaction** | **µL Per Master Mix** |
| Nuclease-free Water | 4.25 | |
| SSIV 2X Reaction Mix | 12.5 | |
| SuperScript IV RT Mix | 0.25 | |
| S1 or S2 Primer Pool | 5 | |
| **Subtotal** | **22** | |

10.1.1 Aliquot 22 µL of each reaction mix into respective wells of a 96-well PCR plate or into 0.2 mL PCR tubes.

10.1.2 For each sample, positive control, and negative control add 3 µL of RNA or water to an S1 master mix containing well and a corresponding S2 master mix containing well.

**Figure 1**:

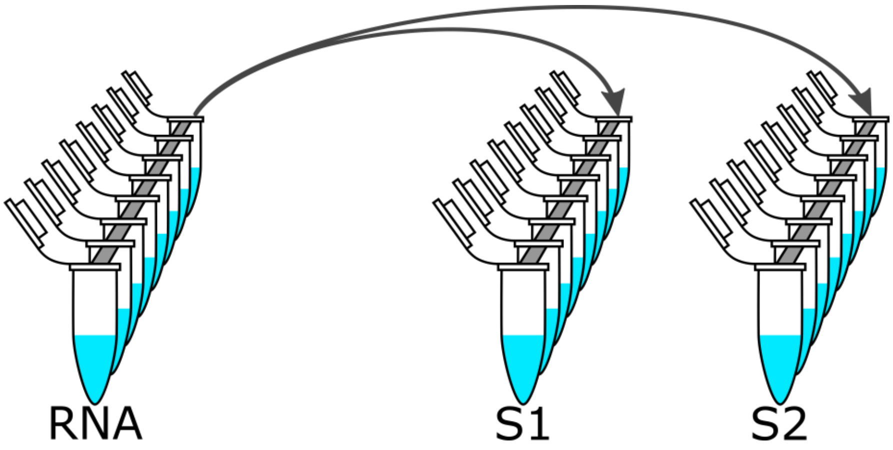

10.2 Seal, gently mix, centrifuge, and incubate.

10.2.1 Securely seal to ensure no evaporation occurs.

| Table 4: Cycling conditions | | |
|---|---|---|
| **Step** | **Temperature (°C)** | **Time (mm:ss)** |
| **1** | 50 | 10:00 |
| **2** | 98 | 2:00 |
| **3** | 98 | 0:10 |
| **4** | 60 | 0:10 |
| **5** | 72 | 1:15 |
| **6** | Repeat steps 3-5 for 40 total cycles | |
| **7** | 72 | 5:00 |
| **8** | 4 | hold |

#### 11.0 Amplicon QC

11.1 QC amplicons via electrophoresis.

11.1.1 S1: 2.5 kb

11.1.2 S2: 2.2 kb

#### 12.0 Combining S1 and S2 Amplicons

12.1 Combine corresponding S1 and S2 reactions into 50 µL amplicon pools.

**Figure 2**:

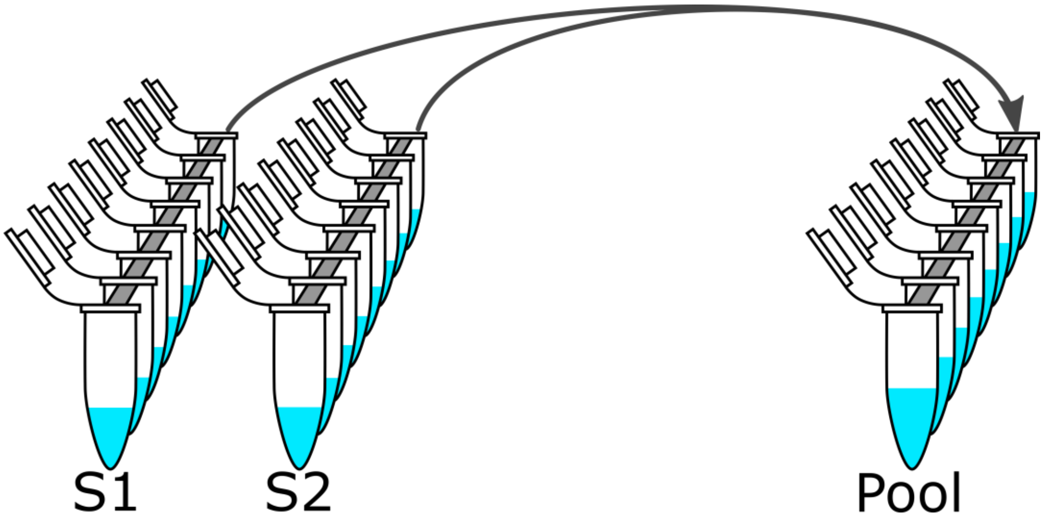

#### 13.0 SPRI bead cleanup with Ethanol wash

13.1 Bring SPRI beads to room temperature (15-25°C) and mix by vortexing.

13.2 Prepare fresh 80% ethanol.

13.3 Add 1x (50 µL) SPRI beads to each sample, mix gently, and incubate at room temperature (15-25°C) for 5 min.

13.3.1 Spin down the samples, pellet on a magnet until the supernatant is clear, then remove and discard the supernatant.

13.4 With the samples on the magnet and without disturbing the pellet:

13.4.1 Add 200 µL of 80% ethanol to each sample.

13.4.2 Remove and discard the ethanol.

13.4.3 Add 200 µL of 80% ethanol to each sample.

13.4.4 Remove and discard the ethanol.

13.5 Spin down the samples, pellet on a magnet, and remove any residual ethanol.

13.5.1 Do not allow the beads to dry to the point of cracking.

13.6 Remove the samples from the magnet, add 15 µL of nuclease-free water, gently resuspend, and incubate at room temperature (15-25°C) for 2 min.

13.6.1 Spin down the samples, pellet on a magnet until the supernatant is clear, then remove and retain the cleaned amplicons in a new plate.

#### 14.0 Proceed to Sequencing

14.1 The amplicons produced here are suitable for nanopore sequencing as described in LP-472 - Library Preparation and Nanopore Sequencing of SARS-CoV-2 S-gene Amplicons. A minimum of 8 samples (including controls) is recommended for library yield.

14.2 The user may decide to sequence the amplicons produced here via other methods of library preparation and or other sequencing platforms that are suitable for 2.2 kb and 2.5 kb amplicons; however, it is the responsibility of that user to validate the chosen sequencing method. It is not recommended to use sequencing data that is of partial or low coverage or low quality.

#### 15.0 Related Procedures

15.1 LP-472 - Library Preparation and Nanopore Sequencing of SARS-CoV-2 S-gene Amplicons

#### 16.0 References

16.1 User Guide: SuperScript™ IV One-Step RT-PCR System (Invitrogen)
