## SupplementalMaterials for "Targeted Amplification and Genetic Sequencing of the Severe Acute Respiratory Syndrome Coronavirus 2 Surface Glycoprotein": Text_S02_LP-472 - Library Preparation and Nanopore Sequencing of SARS-CoV-2 S-gene Amplicons.pdf

DEPARTMENT OF HEALTH & HUMAN SERVICES
Public Health Service
Centers for Disease Control and Prevention (CDC)
Atlanta, GA 30329

### Library Preparation and Nanopore Sequencing of SARS-CoV-2 S-gene Amplicons

***Virology, Surveillance and Diagnosis Branch, Genomics and Diagnostics Team (GDT)***

**NOTE: This procedure is provided for research use only. This document is not intended to be used for commercial development or for-profit testing. Use of trade names and commercial sources is for identification only and does not constitute endorsement by the Public Health Service or by the United States Department of Health and Human Services. Please do not distribute this document to other laboratories or commercial entities.**

#### 1.0 Purpose

1.1 The purpose of this procedure is to describe the library preparation and nanopore sequencing of SARS-CoV-2 S-gene amplicons derived from LP-471 – Reverse Transcription-PCR (RT-PCR) of the SARS-CoV-2 S-gene for Sequencing.

#### 2.0 Definitions

2.1 Library Preparation: The modification of nucleic acids into a suitable state as to be loaded onto a sequencing device.

#### 3.0 Equipment (Use Examples or Equivalent)

3.1 Sterile, nuclease-free 1.5 mL micro-centrifuge tubes

3.2 0.2 mL PCR reaction tube strips or plates

3.2.1 PCR 8-tube strips (Brand Tech Scientific Inc. Catalog. No. 781332)

3.2.2 PCR Plate, 96-well, semi-skirted, flat deck (Life Technologies, Catalog No. AB-1400)

3.2.3 TempPlate pierceable sealing foil, sterile (USA Scientific: Catalog No. 2923-0110)

3.2.4 Sealing Roller (BIO-RAD: Catalog No. MSR-0001)

3.2.5 Silicone compression mat (Sigma Aldrich: Catalog No. AXYCMFLAT)

3.3 Vortex

3.4 1.5 mL tube, 0.2 mL strip tube, and 96 well plate compatible centrifuge

3.5 Cold blocks for 0.2 mL and 1.5 mL PCR reaction tubes (ISC BioExpress)

3.6 1.5 mL tube, 8-tube strip, or 96-well format magnetic separation rack (Life Technologies DynaMag-2: Catalog No. 12321D, Alpaqua Magnum FLX: Catalog No. A000400)

3.7 Pipettes (10 µL, 20 µL, 200 µL, and 1000 µL)

3.7.1 Multichannel pipettes also recommended (10 µL, 20 µL, and 200 µL)

3.7.2 Corresponding aerosol barrier pipette tips

3.8 Disposable reagent reservoir, sterile (Axygen: Catalog No. RES-V-25-S)

3.9 96-well format PCR Thermocycler System (BioRad T100)

3.10 Qubit fluorometer for DNA quantification (Thermo Fischer: Catalog No. Q33238)

3.11 Mk1C or GridION nanopore sequencing device (Oxford Nanopore Technologies)

#### 4.0 Reagents

4.1 Nuclease-free water

4.2 SPRI beads (Beckman Coulter: Catalog No. A63880, A63881, A63882, or equivalent)

4.3 Molecular biology grade absolute ethanol

4.4 Blunt/TA Ligase Master Mix (NEB: Catalog No. M0367)

4.5 NEBNext Ultra II End repair/dA-tailing Module (NEB: Catalog No. E7546)

4.6 NEBNext Quick Ligation Module (NEB: Catalog No. E6056)

4.7 Native barcoding kit (Oxford Nanopore Technologies: Catalog No. EXP-NBD104, EXP-NBD114, or EXP-NBD196)

4.8 Adapter Mix II Expansion (Oxford Nanopore Technologies: Catalog No. EXP-AMII001)

4.9 Sequencing Auxiliary Vials (Oxford Nanopore Technologies: Catalog No. EXP-AUX001)

4.10 Short fragment buffer expansion (Oxford nanopore Technologies: Catalog No. EXP-SFB001)

4.11 Flow Cell Priming Kit (Oxford nanopore Technologies: Catalog No. EXP-FLP002)

4.12 Nanopore flow cells: disposable flongle flow cells and or standard MinION flow cells

4.12.1 Disposable flongle flow cells (Oxford Nanopore Technologies: Catalog No. FLGIntSP and or FLO-FLG001)

4.12.1.1 Flongle Sequencing Expansion kit (EXP-FSE001)

4.12.1.2 Flongle Adapter (Included with FLGIntSP)

4.12.2 Standard MinION flow cells (Oxford Nanopore Technologies: Catalog No. FLO-MIN106D)

4.13 Qubit dsDNA HS assay kit (Thermo Fisher Scientific: Catalog No. Q32851 – 100 assays or Q32854 – 500 assays)

#### 5.0 Positive and Negative Controls

5.1 Controls should be carried over from RT-PCR as described in LP471 – Reverse Transcription-PCR (RT-PCR) of the SARS-CoV-2 S-gene for Sequencing.

**Table 1: Positive and Negative Controls**

| Control | Material | Frequency | Expected Outcome |
|---|---|---|---|
| Positive | Twist Bioscience Custom NGS Spike Control<br>e.g. Delta REF: 103885; Omicron REF: 103885<br>Diluted to 20K copies/µL<br>Or<br>Previously sequenced RNA from propagated isolate of currently circulating SARS-CoV-2 variant, Ct <20<br>Or<br>Previously sequenced RNA from clinical sample of currently circulating SARS-CoV-2 variant, Ct <20 | Every run | Pass Coverage and Quality |
| Negative | Water | Every run | Fail Coverage and Quality |

#### 6.0 Starting Material

6.1 The starting material is anticipated to be final product of LP-471 – Reverse Transcription-PCR (RT-PCR) of the SARS-CoV-2 S-gene for Sequencing. Specifically, this should be SARS-CoV-2 S-gene amplicons where the corresponding S1 and S2 reactions have been combined and cleaned.

#### 7.0 Safety Precautions

7.1 Personal Protective Equipment (PPE) - Wear lab coats, gloves, and safety glasses.

7.2 Positive controls should be handled in an approved BSL-2 handling area to avoid contamination of laboratory equipment and reagents that could cause false positive results.

7.3 Materials should be handled in accordance with Good Laboratory Practices.

#### 8.0 Comments and Questions

#### 9.0 Gather Reagents

9.1 Thaw and store at the indicated temperature during the procedure.

9.2 Flick/invert the reagent tubes to ensure they are well mixed, and spin down before opening.

9.3 The Ultra II End Prep Buffer may contain some precipitate. Bring to room temperature, pipette to break up the precipitate, and vortex to ensure the reagent is thoroughly mixed.

9.4 **Keep enzymes at -20°C**

9.4.1 Ultra II End Prep Enzyme Mix

9.4.2 Blunt TA MM

9.4.3 Quick T4 DNA Ligase

9.5 **+4°C**

9.5.1 Cleaned Amplicons

9.5.2 Ultra II EP Buffer

9.5.3 EXP-NBD196 plate of barcodes

9.5.4 SFB

9.5.5 5x Quick Ligation Buffer

9.5.6 AMII

9.5.7 EB

9.5.8 SBII (glass vial) for flongle or SQB (plastic tubes) for standard MinION flow cells

9.5.9 LBII (glass vial) for flongle or LB (plastic tubes) for standard MinION flow cells

9.5.10 FB (glass vial) for flongle or FB (plastic tubes) for standard MinION flow cells

9.5.11 FLT

9.6 **Room temperature (15-25°C).**

9.6.1 Nuclease-free water

9.6.2 SPRI beads

9.6.3 Ethanol

#### 10.0 End-Prep

10.1 Prepare a master mix:

| Table 1: End-Prep master mix | | |
|---|---|---|
| **Reagent** | **Volume (µL) Per Reaction** | **µL Per Master Mix** |
| Nuclease-Free Water | 3.3 | |
| Ultra II End-Prep Buffer | 1.2 | |
| Ultra II End-Prep Enzyme | 0.5 | |
| **Total** | **5** | |

10.2 For each sample, transfer 5 µL of the end-prep master mix to the wells of a new plate.

10.3 For each sample add 5 µL of cleaned amplicons to the end-prep master mix containing wells.

10.3.1 Seal and retain the remaining the clean amplicons.

10.4 Seal, mix, centrifuge, and incubate the end-prep plate.

10.4.1 Securely seal to ensure no evaporation occurs.

| Table 2: End-Prep incubation conditions | |
|---|---|
| **Temperature (°C)** | **Time (mm:ss)** |
| 20 | 15:00 |
| 65 | 15:00 |
| 4 | hold |

#### 11.0 Native Barcoding

11.1 Prepare a master mix of diluted Blunt TA Master Mix for each sample:

| Table 3: Diluted Blunt TA Master Mix | | |
|---|---|---|
| **Reagent** | **Volume (µL) Per Reaction** | **µL Per Master Mix** |
| Nuclease-Free Water | 3 | |
| Blunt TA Master Mix | 5 | |
| **Total** | **8** | |

11.2 Add to each well of a new plate:

| Table 4: Native barcoding reaction | |
|---|---|
| **Reagent** | **Volume (µL)** |
| Diluted Blunt TA Master Mix | 8 |
| Unique Native Barcodes | 1.25 |
| End prepped samples | 0.75 |
| **Total** | **10** |

11.2.1 Seal and retain the remaining material in end-prep plate.

11.3 Seal, mix, centrifuge, and incubate the barcoding plate.

11.3.1 Securely seal to ensure no evaporation occurs.

**Table 5: Native barcoding incubation conditions**

| Temperature (°C) | Time (mm:ss) |
|---|---|
| 20 | 20:00 |
| 65 | 10:00 |
| 4 | hold |

11.4 Pool the samples:

11.4.1 8-24 samples: entire 10 µL reactions.

11.4.2 25-48 samples: 5 µL from each reaction.

11.4.3 49-96 samples: 2.5 µL from each reaction.

11.4.4 Seal and retain any remaining material in barcoding plate.

#### 12.0 SPRI Bead Clean Up with SFB and Ethanol Wash

12.1 Calculate or measure the volume of the pool with a pipette and add 0.4x of SPRI beads to the sample, mix gently, and incubate at room temperature (15-25°C) for 5 min.

12.1.1 e.g., for 240 µL of pooled sample, add 96 µL of beads.

12.1.2 Spin down the sample, pellet on a magnet until the supernatant is clear, then remove and discard the supernatant.

12.2 Remove from the magnet, add 250 µL short fragment buffer (SFB), flick to resuspend.

12.2.1 Spin down the sample, pellet on a magnet until the supernatant is clear, then remove and discard the supernatant.

12.3 With the samples on the magnet and without disturbing the pellet:

12.3.1 Add 250 µL of 80% ethanol to the sample.

12.3.2 Remove and discard the ethanol.

12.4 Spin down the sample, pellet on a magnet, and remove any residual ethanol.

12.4.1 Do not allow the beads to dry to the point of cracking.

12.5 Remove from the magnet, add 30 µL of water, gently resuspend, and incubate at room temperature (15-25°C) for 2 min.

12.5.1 Spin down the sample, pellet on a magnet until the supernatant is clear, then remove and retain the cleaned barcoded amplicons.

#### 13.0 Adapter Ligation

13.1 Mix:

| Table 6: Adapter ligation reaction | |
|---|---|
| **Reagent** | **Volume (µL)** |
| Barcoded, pooled, and cleaned amplicons from previous step | 30 |
| NEBNext Quick Ligation Reaction Buffer (5x) | 10 |
| Adaptor Mix (AMII) | 5 |
| Quick T4 DNA Ligase | 5 |
| **Total** | **50** |

13.2 Incubate at room temperature for 20 minutes.

#### 14.0 SPRI Bead Clean Up with SFB Wash and EB Elution

14.1 Add 1x (50 µL) of SPRI beads the sample, mix gently, and incubate at room temperature (15-25°C) for 5 min.

14.1.1 Spin down the sample, pellet on a magnet until the supernatant is clear, then remove and discard the supernatant.

14.2 Remove from the magnet, add 250 µL short fragment buffer (SFB), flick to resuspend.

14.2.1 Spin down the sample, pellet on a magnet until the supernatant is clear, then remove and discard the supernatant.

14.3 Remove from the magnet, add 250 µL short fragment buffer (SFB), flick to resuspend.

14.3.1 Spin down the sample, pellet on a magnet until the supernatant is clear, then remove and discard the supernatant.

14.3.2 Spin down the sample and remove any residual SFB.

14.4 Remove from the magnet, add 15 µL of elution buffer (EB), gently resuspend, and incubate at room temperature (15-25°C) for 10 min.

14.4.1 Spin down the sample, pellet on a magnet until the supernatant is clear, then remove and retain the eluant.

14.4.2 Quantify 2 µL of the eluant.

#### 15.0 Prepare the Flow Cell

15.1 If applicable and possible, power cycle the GridION.

15.2 Insert the flow cell and perform a flow cell check.

#### 16.0 Sequencing on Flongle Flow Cells: FLO-FLG001 (for FLO-MIN106, skip this section)

16.1 Mix the final library using the following calculations and table:

16.1.1 $(80\, ng\ target) \div (quantified\ DNA\ in \frac{ng}{\mu L}) = \mu L\ of\ DNA\ to\ add\ to\ final\ library$

16.1.1.1 A maximum of 5 µL can be added.

16.1.2 Make up the remaining 5 µL with EB.

16.2 Use SBII and LBII from glass vials provided by Flongle Sequencing Expansion kit (EXP-FSE001) that ships with FLO-FLG001.

**Table 7: FLO-FLG001 final DNA library mixture**

| Reagent | Volume (µL) |
|---|---|
| Sequencing Buffer II (SBII) | 15 |
| Resuspended Library Loading Beads II (LBII) | 10 |
| DNA in EB: ≤80 ng | 5 |
| **Total** | **30** |

16.3 Prepare for priming.

16.3.1 Use FB from the glass vial provided by Flongle Sequencing Expansion kit (EXP-FSE001) that ships with FLO-FLG001.

16.3.2 Prepare the priming mix by adding 3 µL of mixed Flush Tether (FLT) to 117 µL of Flush Buffer (FB).

16.3.3 Peel back the plastic cover.

16.3.4 Aspirate any storage solution from the tape or outside the loading port.

16.4 Priming with 120 µL of priming mix.

16.4.1 Carefully avoid introducing air bubbles.

16.4.2 Pipette priming mix into the loading port by rolling or slowly depressing.

16.4.3 Remove the pipette tip before air is dispensed into the flow cell.

16.4.4 Do not allow the pipette to draw the solution backwards.

16.5 Loading 30 µL library.

16.5.1 Carefully avoid introducing air bubbles.

16.5.2 Mix the final library by pipetting to resuspend the loading beads.

16.5.3 The library may be loaded by inserting the pipette into the loading port or dropwise onto the loading port

16.5.3.1 Pipette library into the loading port by rolling or slowly depressing.

16.5.3.2 Remove the pipette tip before air is dispensed into the flow cell.

16.5.3.3 Do not allow the pipette to draw the solution backwards.

16.5.4 Gently seal the adhesive plastic over the loading port and vents.

#### 17.0 Sequencing on Standard MinION Flow Cells: FLO-MIN106

17.1 Priming

17.1.1 Prepare the priming mix by adding 30 µL of mixed Flush Tether (FLT) to a new plastic tube of Flush Buffer (FB).

17.1.2 Open the priming port.

17.1.3 Carefully avoid introducing air bubbles.

17.1.4 Prime with 800 µL of priming mix into the priming port > 5 min before loading.

17.2 Mix the final library using the following calculations and table:

17.2.1 $(80\, ng\ target) \div (quantified\ DNA\ in \frac{ng}{\mu L}) = \mu L\ of\ DNA\ to\ add\ to\ final\ library$

17.2.1.1 A maximum of 12 µL can be added.

17.2.2 Make up the remaining 12 µL with EB.

| Table 8: FLO-MIN final DNA library mixture | |
|---|---|
| **Reagent** | **Volume (µL)** |
| Sequencing buffer (SQB) | 37.5 |
| Resuspended library loading beads (LB) | 25.5 |
| DNA library: ≤80 ng in elution buffer (EB) | 12 |
| **Total** | **75** |

17.3 Loading

17.3.1 With the priming port still open.

17.3.2 Open the spot-on port.

17.3.3 Add 200 µL of priming mix into the priming port.

17.3.3.1 Carefully avoid introducing air bubbles.

17.3.3.2 Roll or slowly depress the pipette to dispense the priming mix into the flow cell.

17.3.3.3 Remove the pipette tip before air is dispensed into the flow cell.

17.3.3.4 Do not allow the pipette to draw the solution backwards as this will draw air into the now open spot-on port.

17.3.4 Immediately load the 75 µL final library into the spot-on port.

17.3.4.1 Pipette up and down to resuspend the loading beads.

17.3.4.2 In a drop-wise fashion, roll or slowly depress the pipette to dispense the library onto the spot-on port.

17.3.5 Close all ports and lids.

#### 18.0 Starting the Sequencing Run

18.1 No spaces in experiment or sample name.

18.2 Kit Selection:

18.2.1 SQK-LSK109, EXP-NBD196

18.3 Run options: Run time

18.3.1 FLO-FLG001: 12-24 hours

18.3.2 FLO-MIN106: ~30 min/barcode used

18.3.2.1 12 hours is an acceptable default

18.4 Basecalling:

18.4.1 GridION: Live – super accuracy basecalling enabled

18.4.2 MinION Mk1C: Live – high accuracy basecalling enabled

18.4.2.1 For lower yield flongle flow cells, the MinION Mk1C basecalling may lag during the run.

18.4.2.2 For higher yield standard MinION flow cells, the MinION Mk1C basecalling will lag significantly behind raw read production. If live or faster data is required from the Mk1C, use the fast basecalling model.

18.4.3 Barcoding - enabled

18.4.3.1 Trim barcodes

18.4.3.2 Require barcodes on both ends

18.5 Output:

18.5.1 Reads per FAST5 and FASTQ

18.5.1.1 Defaults cannot be changed on Mk1C

18.5.1.1.1 FLO-FLG001: 1K reads/file

18.5.1.1.2 FLO-MIN106: 4K reads/file

18.5.1.2 GridION

18.5.1.2.1 FLO-FLG001: 5K reads/file

18.5.1.2.2 FLO-MIN106: 10K reads/file

18.5.2 Filtering

18.5.2.1 Qscore filtering at default value

18.5.2.2 Min read length 1 kb

#### 19.0 Related Procedures

19.1 LP-471 – Reverse Transcription-PCR (RT-PCR) of the SARS-CoV-2 S-gene for Sequencing

#### 20.0 References

20.1 Native barcoding amplicons (with EXP-NBD104, EXP-NBD114, and SQK-LSK109): Oxford Nanopore Technologies

20.1.1 https://store.nanoporetech.com/us/productDetail/?id=native-barcoding-expansion-96

20.1.2 https://community.nanoporetech.com/protocols/amplicon-barcoding-with-native-barcoding-expansion-96-exp-nbd196-and-sqk/v/nba_9102_v109_revf_09jul2020

20.2 Flow cell Loading

20.2.1 https://community.nanoporetech.com/protocols/lambda-control-sqk-lsk109/v/cde_9062_v109_revaf_14aug2019/priming-and-loading-the-sp?devices=flongle

20.2.2 https://community.nanoporetech.com/protocols/lambda-control-sqk-lsk109/v/cde_9062_v109_revaf_14aug2019/priming-and-loading-the-sp?devices=minion

20.2.3 https://community.nanoporetech.com/protocols/lambda-control-sqk-lsk109/v/cde_9062_v109_revaf_14aug2019/priming-and-loading-the-spoton-gridion?devices=gridion

20.2.4 https://community.nanoporetech.com/docs/prepare/library_prep_protocols/minion-mk1c-user-manual/v/mkc_2005_v1_revw_27nov2019
