## SupplementalMaterials for "Targeted Amplification and Genetic Sequencing of the Severe Acute Respiratory Syndrome Coronavirus 2 Surface Glycoprotein": Text_S03_Supplemental_Legends.docx

### Texts

#### Text S01: LP-471

Reverse Transcription-PCR (RT-PCR) of the SARS-CoV-2 S-gene for Sequencing

#### Text S02: LP-472

Library Preparation and Nanopore Sequencing of SARS-CoV-2 S-gene Amplicons

#### Text S03: Supplemental Legends

### Tables

#### Table S01: Primer Candidates

Fourteen individual primers were considered for use with this method. Initially, three primers candidates were identified for each position: S1F, S1R, S2F, S2R. Later, an additional candidate for S2R was added which corresponded to an area of homology between SARS-CoV and SARS-CoV-2, along with a degenerate version at the same location.

#### Table S02: Primer Conservation as of 2021-03-24

Primer candidate conservation was evaluated against SARS-CoV-2 genomic data: all data, last 3 months, and for major variants. This was conservation analysis was used in the primer selection process.

#### Table S03: Primer Combinations

Twenty primer combination were evaluated for their ability to amplify either the S1 or S2 fragment. The results of the iterative testing and selection process is summarized. Initially, nine combinations were tested for each amplicon: S1 or S2. After primer mixtures 1-8 and 2-4 were selected for S1 and S2, respectively, S2v2 and S2v2-R7. S2v2 was retained as an alternate and S2v2-R7 was eliminated.

#### Table S04: 12 Primer LOD Ct and Copy Number

Flu SC2 multiplex and copy number results are given for the LOD of B.1.351 (Beta) RNA used in the primer selection process.

#### Table S05: SpikeSeq Primer Conservation against XBB

SARS-CoV-2 XBB surveillance data as of 2023-01-18.

#### Table S06: SpikeSeq Runs

We performed 14 nanopore sequencing runs to validate and characterize SpikeSeq. A summary of those runs is presented here.

#### Table S07: LOD MinION Flow Cell

A synthetic and viral limit of detection using standard MinION (MIN) flow cells.

#### Table S08: LOD Flongle Flow Cell

A synthetic and viral limit of detection using disposable flongle (FLG) flow cells.

#### Table S09: LOD-MIN Water NTCs

Plate layout, barcode assignment, and read mapping of the LOD-MIN sequencing results, along with the 48 additional water NTCs.

#### Table S10: LOD-FLG A549 NTCs

Plate layout, barcode assignment, and read mapping of the LOD-FLG sequencing results, along with the 24 additional A549 RNA (Rp Ct 22) NTCs.

#### Table S11: Validation Results

Ct values (where available), SpikeSeq coverage results, and a comparison of Nextclade results for 321 samples with both SpikeSeq and full genome sequencing data.

#### Table S12: Coverage vs Ct

We compared the coverage results and Ct values for 277 clinical specimens. Samples were grouped by Ct range and assigned a pass if they met the coverage threshold of requiring ≥ 50x coverage at every position. For samples with a Ct value less than 25, 98-99% of samples passed. For samples with Ct values between 25 and 30, 89% of samples passed. For samples with Ct values over 30, 81% of samples passed.

#### Table S13: MinION vs Flongle Coverage

A coverage comparison between libraries that were split and sequenced on both types of flow cells. These samples were limited to 277 NS3 samples through Delta and include all attempted samples, pass or fail.

#### Table S14: Accuracy Comparison

For samples that generated full length alignments (n =251), a three-way blast comparison of each sample was used to compare consensus level accuracy of SpikeSeq amplification and nanopore sequencing (MIN), illumina sequencing of those same amplicons (ILL), and NS3 surveillance results (NS3).

### Figures

#### Figure S01: Receptor Binding Domain Coverage

Sequence completion of the spike protein RBD is displayed from global SC2 surveillance data across different time frames.

#### Figure S02: S1 Temperature Gradient

Fragment analysis of S1 primer candidates across an annealing temperature gradient.

#### Figure S03: S2 Temperature Gradient

Fragment analysis of S2 primer candidates across an annealing temperature gradient.

#### Figure S04: Primer Selection LOD

Fragment analysis of 12 remaining primer candidates across a limit of detection.

#### Figure S05: S1F_21358 US 3mo summary plot

Primer S1F_21358 conservation is visualized for the latest (3 months) United States SARS-CoV-2 surveillance data. Data as of 2023-07-28.

#### Figure S06: S1R_23813 US 3mo summary plot

Primer S1R_23813 conservation is compared to the latest (3 months) of United States SARS-CoV-2 surveillance data. Data as of 2023-07-28.

#### Figure S07: S2F_23288 US 3mo summary plot

Primer S2F_23288 conservation is compared to the latest (3 months) of United States SARS-CoV-2 surveillance data. Data as of 2023-07-28.

#### Figure S08: S2R_25460 US 3mo summary plot

Primer S2R_25460 conservation is visualized for the latest (3 months) United States SARS-CoV-2 surveillance data. Data as of 2023-07-28.

#### Figure S09: S1F_21358 global 3mo summary plot

Primer S1F_21358 conservation is visualized for the latest (3 months) Global SARS-CoV-2 surveillance data. Data as of 2023-07-28.

#### Figure S10: S1R_23813 global 3mo summary plot

Primer S1R_23813 conservation is visualized for the latest (3 months) Global SARS-CoV-2 surveillance data. Data as of 2023-07-28.

#### Figure S11: S2F_23288 global 3mo summary plot

Primer S2F_23288 conservation is visualized for the latest (3 months) Global SARS-CoV-2 surveillance data. Data as of 2023-07-28.

#### Figure S12: S2R_25460 global 3mo summary plot

Primer S2R_25460 conservation is visualized for the latest (3 months) Global SARS-CoV-2 surveillance data. Data as of 2023-07-28.

#### Figure S13: S1F_21358 global all summary plot

Primer S1F_21358 conservation is visualized for the Global SARS-CoV-2 surveillance data. Data as of 2023-07-28.

#### Figure S14: S1R_23813 global all summary plot

Primer S1R_23813 conservation is visualized for all Global SARS-CoV-2 surveillance data. Data as of 2023-07-28.

#### Figure S15: S2F_23288 global all summary plot

Primer S2F_23288 conservation is visualized for all Global SARS-CoV-2 surveillance data. Data as of 2023-07-28.

#### Figure S16: S2R_25460 global all summary plot

Primer S2R_25460 conservation is visualized for all Global SARS-CoV-2 surveillance data. Data as of 2023-07-28.

#### Figure S17: Related Virus Alignment

The SpikeSeq primers are aligned to SARS-CoV-2 and related coronaviruses.

#### Figure S18: LOD MinION Flow Cell

A synthetic and viral limit of detection using standard MinION (MIN) flow cells.

#### Figure S19: LOD Flongle Flow Cell

A synthetic and viral limit of detection using disposable flongle (FLG) flow cells.

#### Figure S20: LOD-MIN Water NTCs

Plate layout, barcode assignment, and read mapping of the LOD-MIN sequencing results, along with the 48 additional water NTCs.

#### Figure S21: LOD-FLG A549 NTCs

Plate layout, barcode assignment, and read mapping of the LOD-FLG sequencing results, along with the 24 additional A549 RNA (Rp Ct 22) NTCs.

#### Figure S22: Pass Rate vs Ct

We compared the coverage results and Ct values for 277 clinical specimens. Samples were grouped by Ct range and assigned a pass if they met the coverage threshold of requiring ≥ 50x coverage at every position. For samples with a Ct value less than 25, 98-99% of samples passed. For samples with Ct values between 25 and 30, 89% of samples passed. For samples with Ct values over 30, 81% of samples passed.

#### Figure S23: MinION vs Flongle Flow Cells

A coverage comparison between libraries that were split and sequenced on both types of flow cells. These samples were limited to the 277 retrospective NS3 samples.

#### Figure S24: SpikeSeq-NS3 Tanglegram

Pairwise phylogenetics of matching samples (n = 321) that passed both SpikeSeq (circles left) and full genome sequencing (circles right). Phylogenetics were exported from Nextclade and visualized using Auspice.

#### Figure S25: Sequencing Completion

This example shows one use case for SpikeSeq to complete full genome sequencing data that is missing important regions. Here, an early BA.2 lineage Omicron virus was missing sequencing data that was necessary to initiate isolate propagation.

### Sequences

#### Seq S01: Primers

Primer sequences used in this study.

#### Seq S02: Related Virus Alignment

The SpikeSeq primers are aligned to SARS-CoV-2 and related coronaviruses.

#### Seq S03: Twist Delta Fragment

The custom synthetic positive control used in this assay.

#### Seq S04: SpikeSeq Data Set

SpikeSeq derived consensus sequences of the 321 samples with available SpikeSeq and full genome results.

#### Seq S05: Full Genome Data Set

Full genome sequencing derived consensus sequences of the 321 samples with available SpikeSeq and full genome results.

#### Seq S06: SpikeSeq Amplification and Illumina Sequencing

SpikeSeq amplification and illumina sequencing derived consensus sequences of 251 samples.
