## Supplementary figures and images for "Targeted Amplification and Genetic Sequencing of the Severe Acute Respiratory Syndrome Coronavirus 2 Surface Glycoprotein"

### Figure_S01_RDB_coverage.pdf

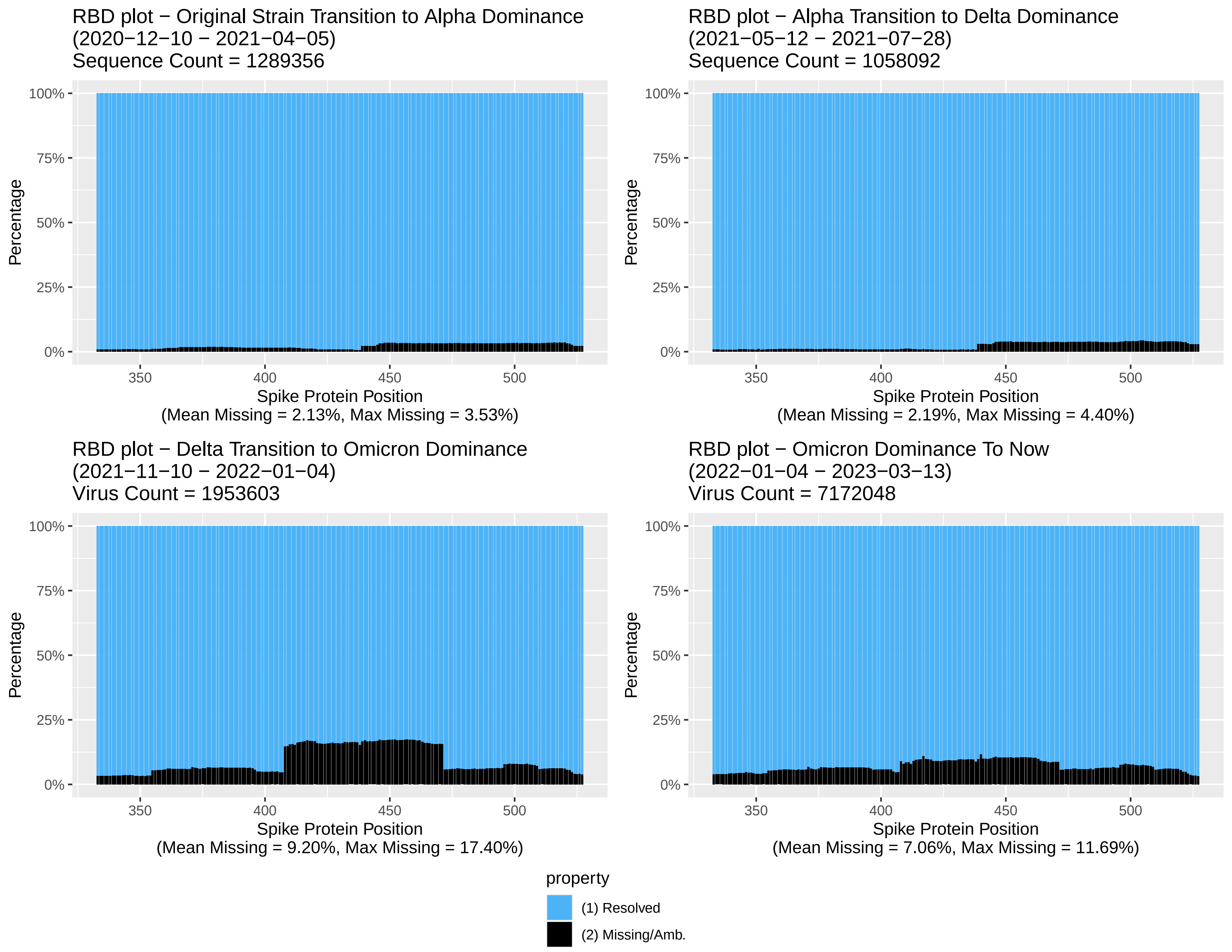

### Figure_S05_SARS_CoV_2_US_3mo_S1F_21358_summary_plot.png

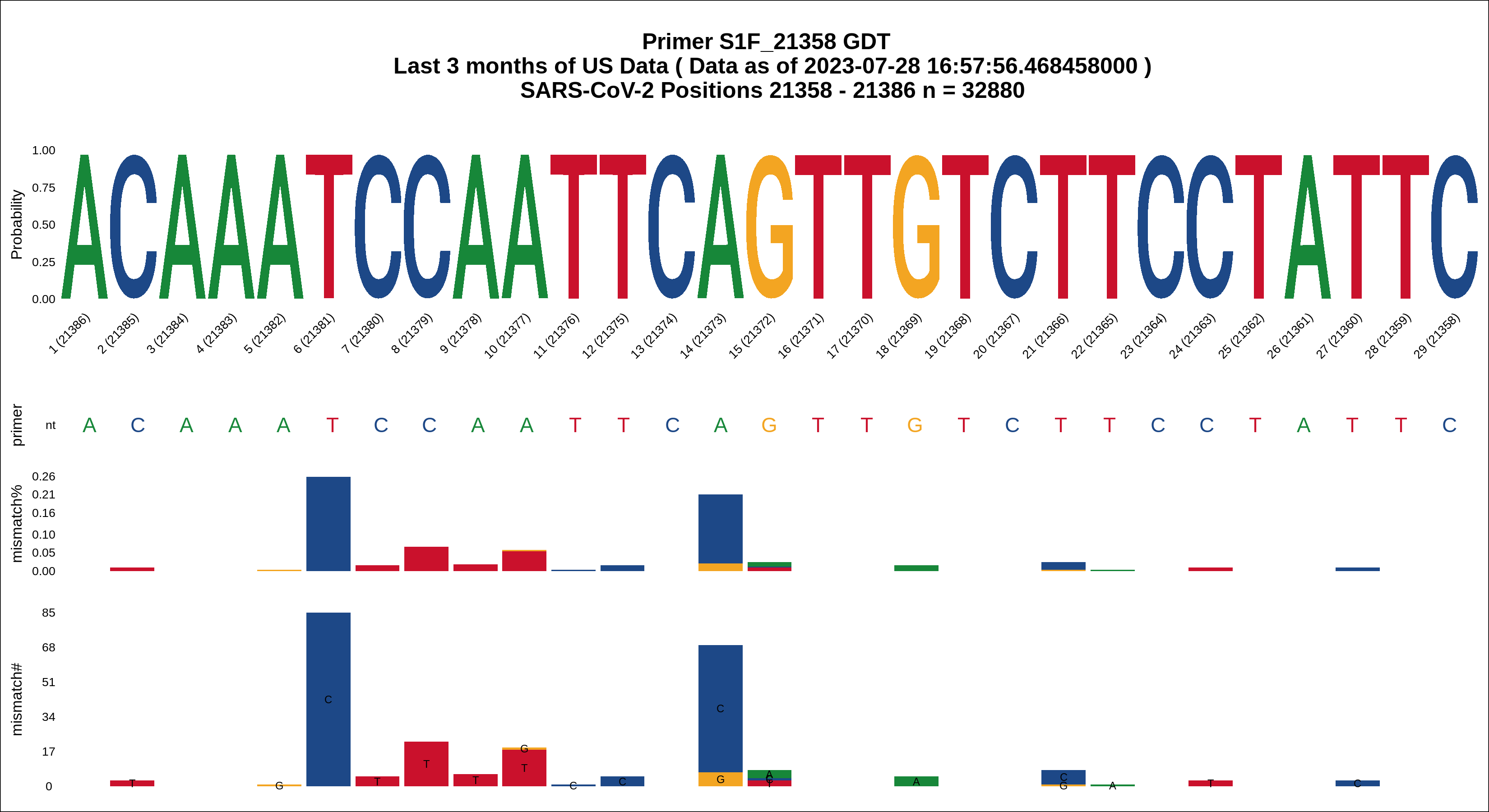

### Figure_S06_SARS_CoV_2_US_3mo_S1R_23813_summary_plot.png

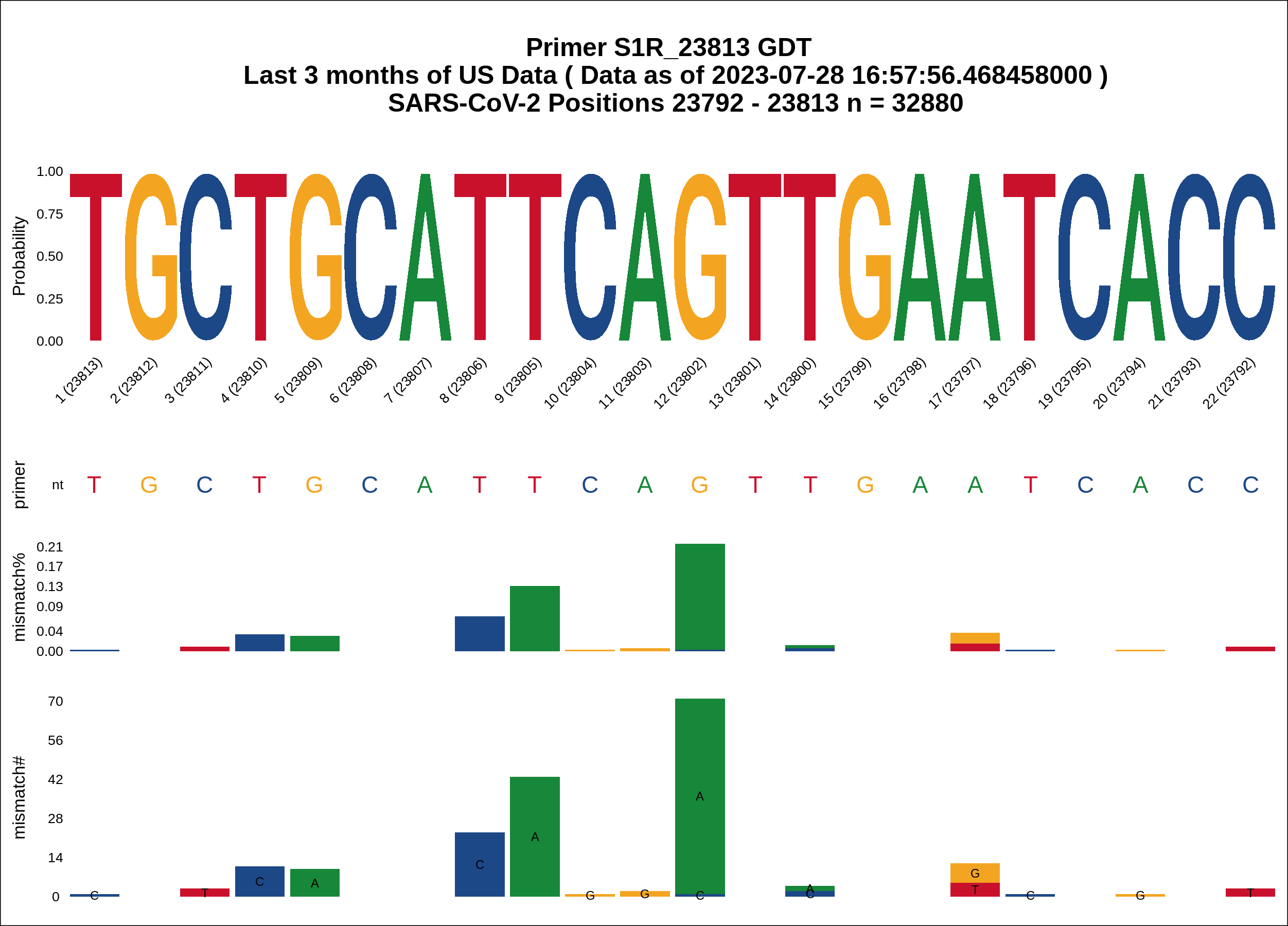

### Figure_S07_SARS_CoV_2_US_3mo_S2F_23288_summary_plot.png

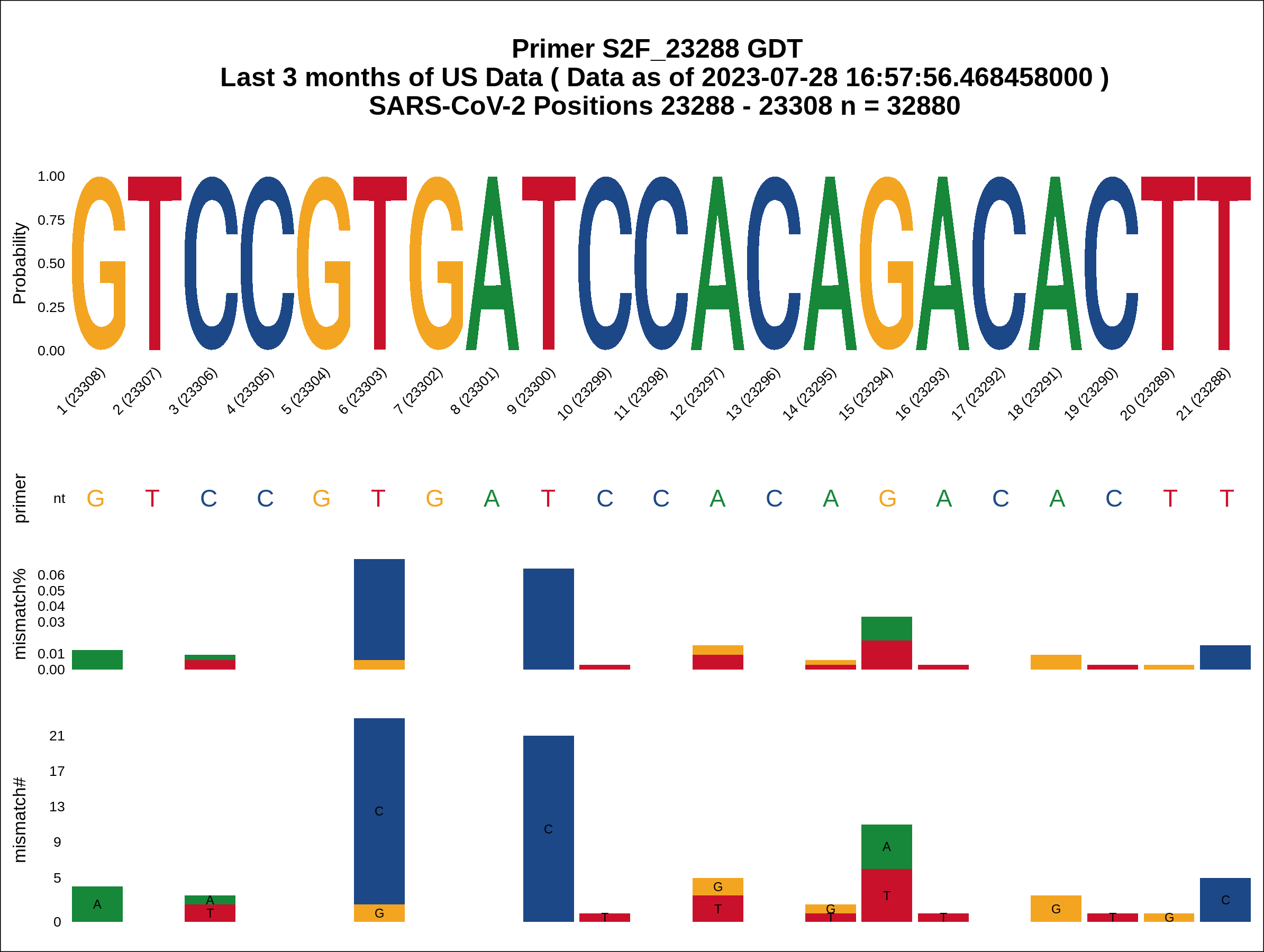

### Figure_S08_SARS_CoV_2_US_3mo_S2R_25460_summary_plot.png

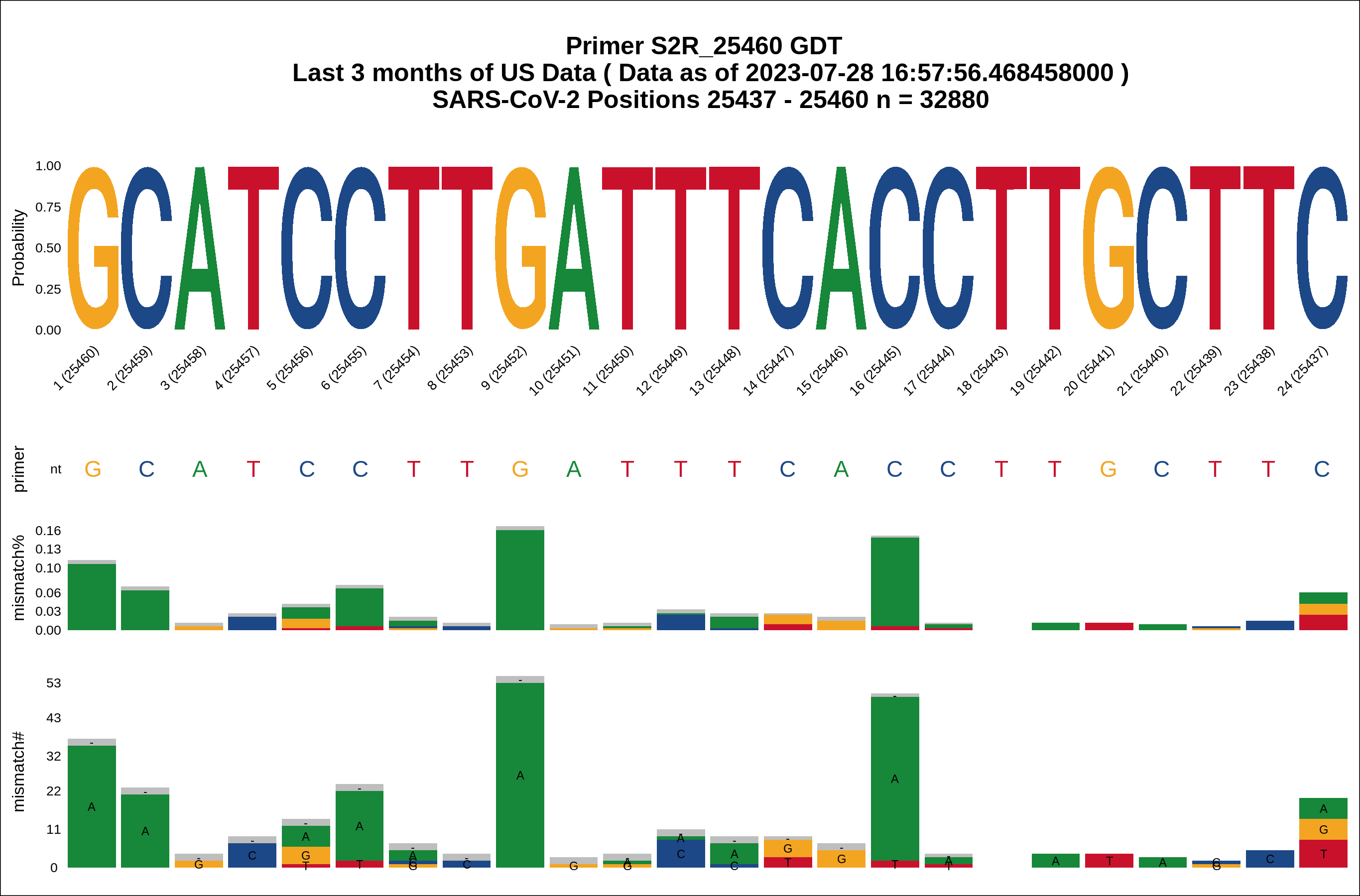

### Figure_S09_SARS_CoV_2_Global_3mo_S1F_21358_summary_plot.png

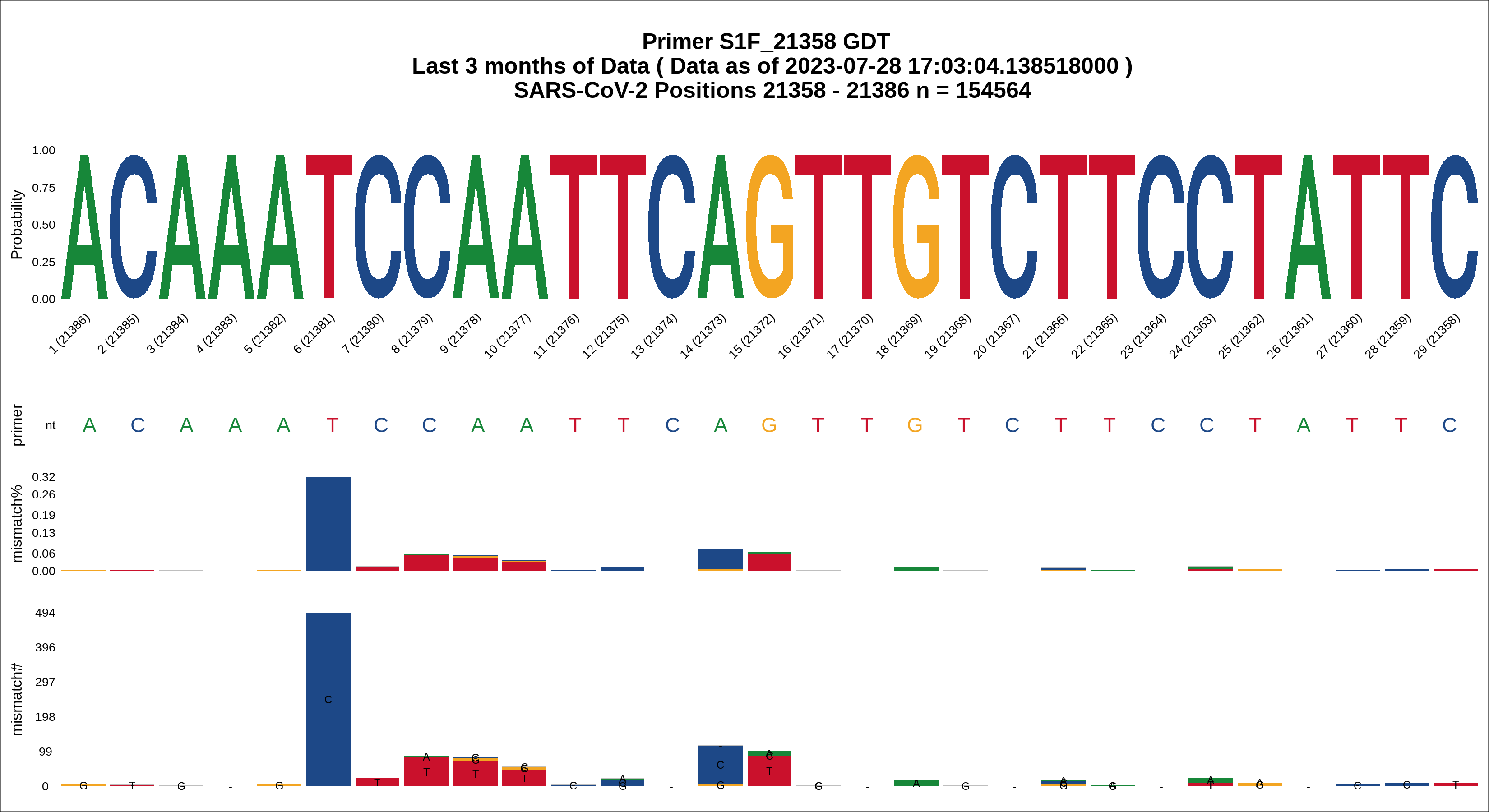

### Figure_S10_SARS_CoV_2_Global_3mo_S1R_23813_summary_plot.png

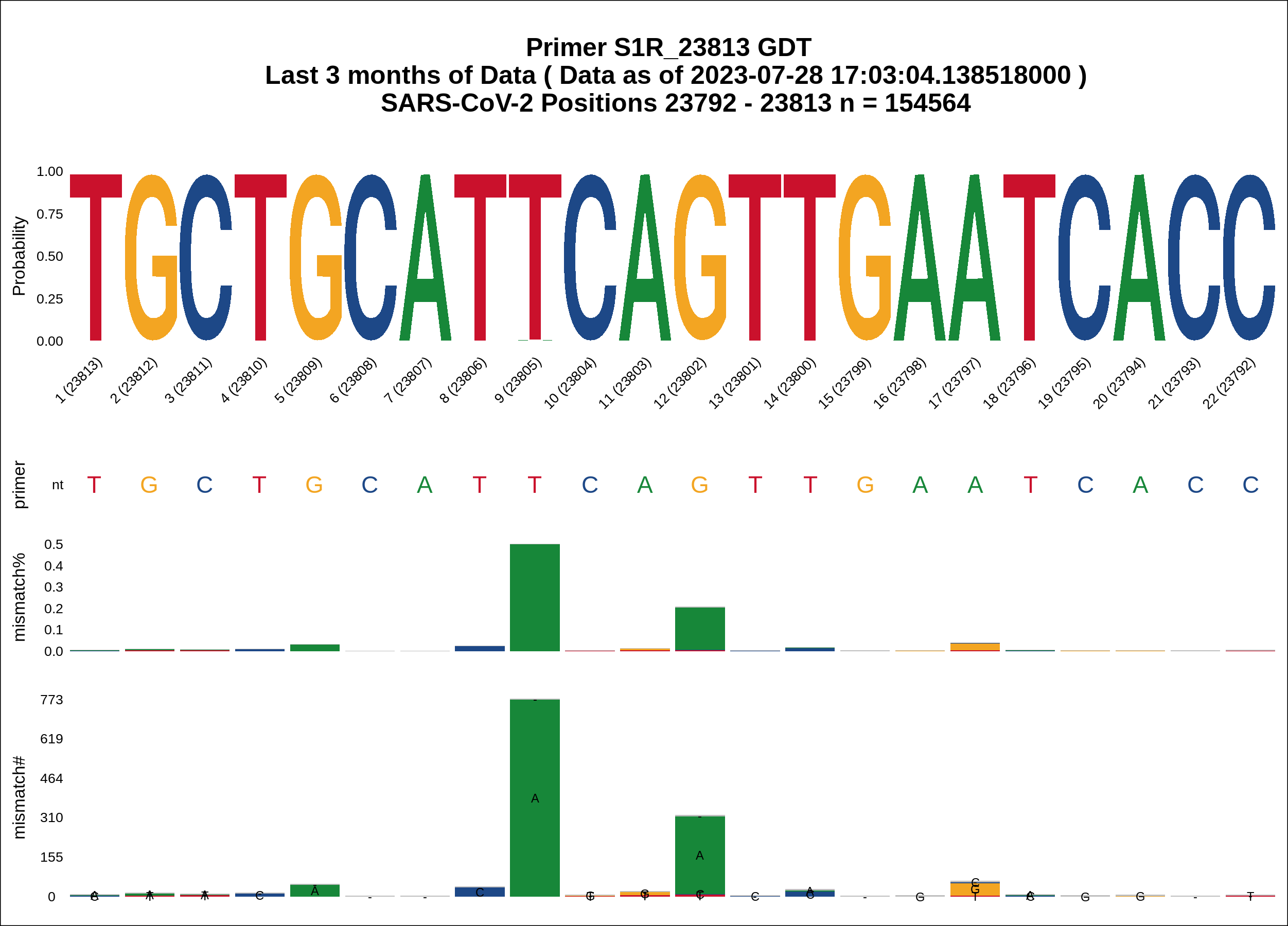

### Figure_S11_SARS_CoV_2_Global_3mo_S2F_23288_summary_plot.png

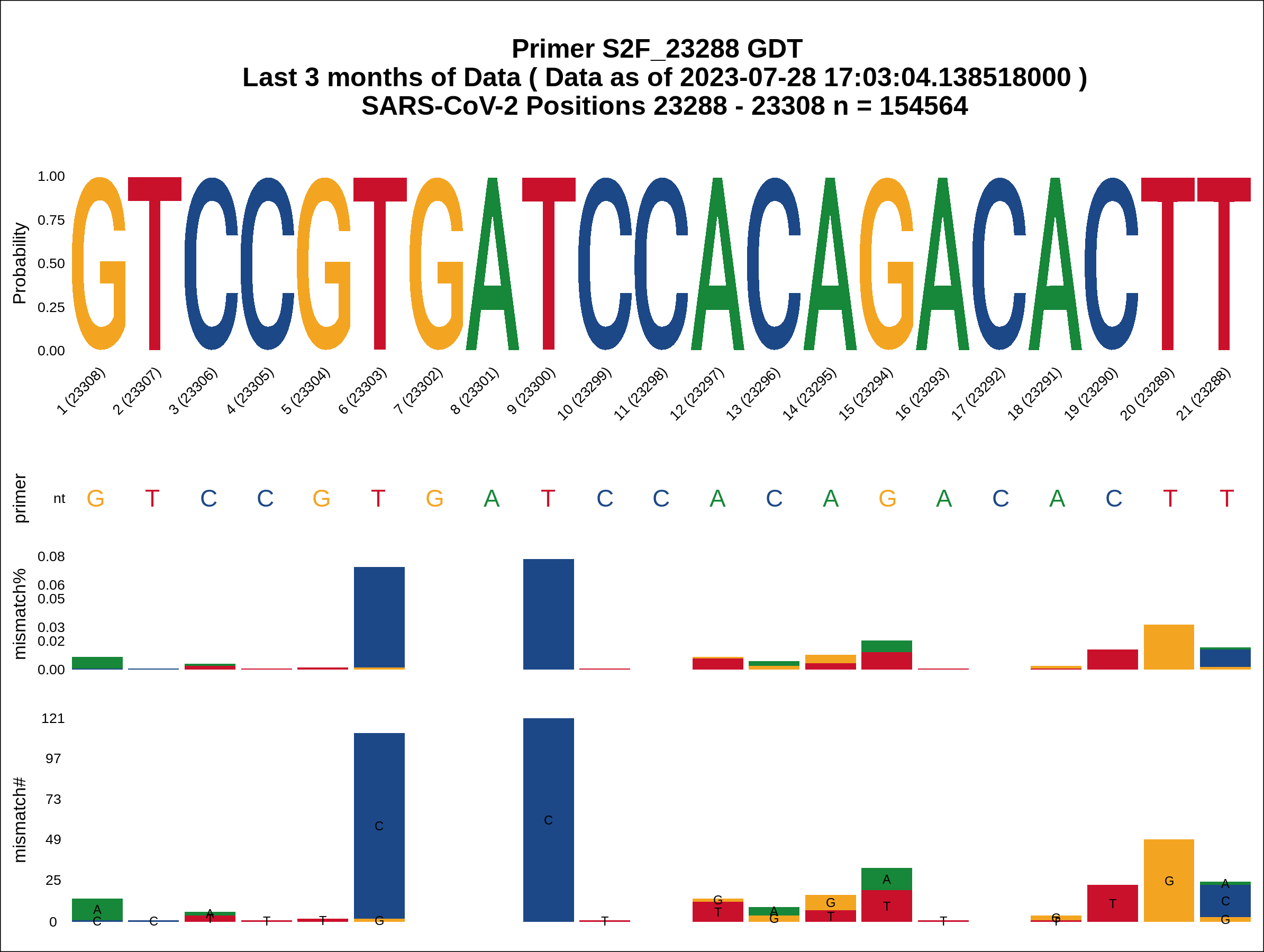

### Figure_S12_SARS_CoV_2_Global_3mo_S2R_25460_summary_plot.png

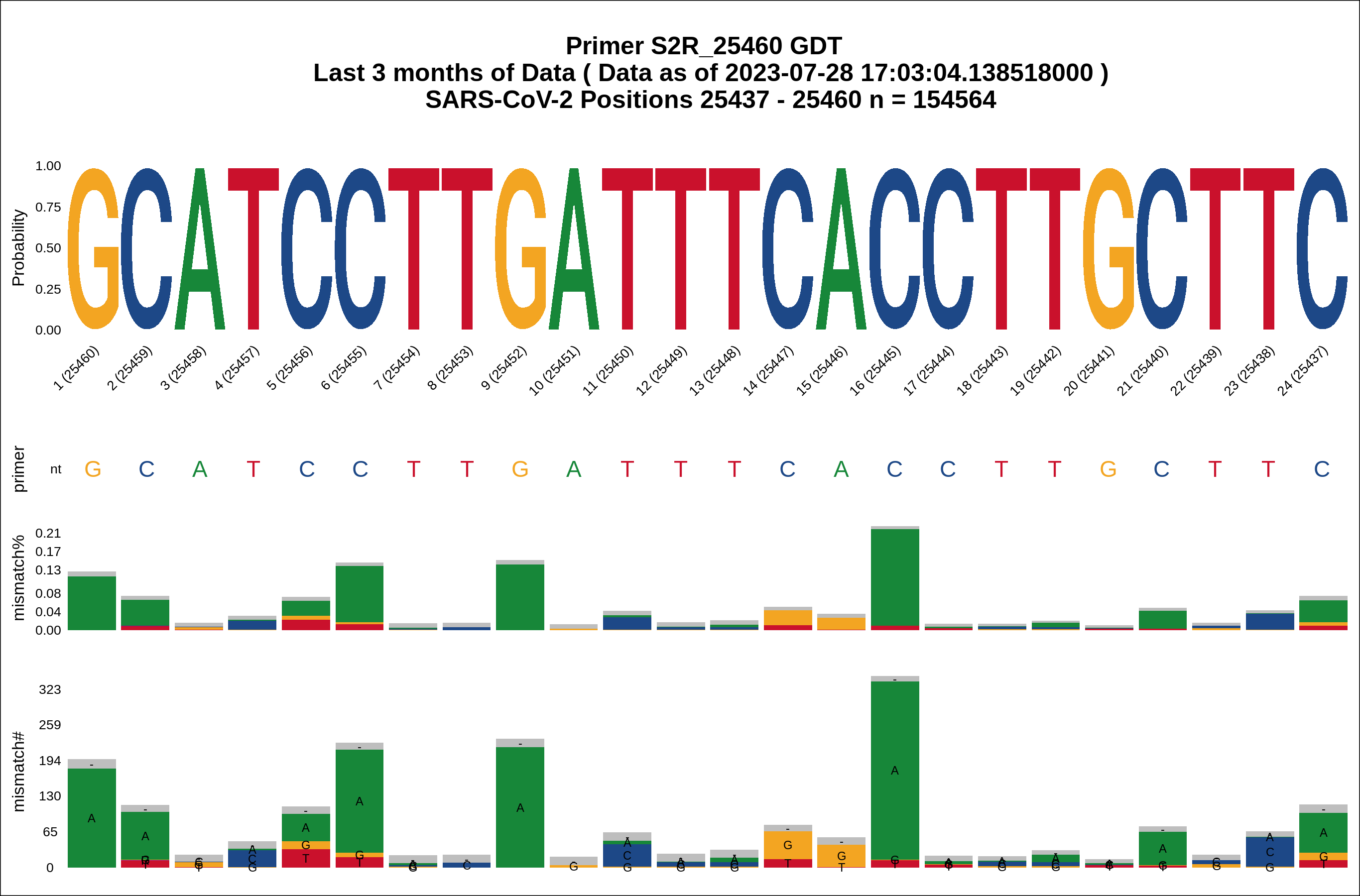

### Figure_S13_SARS_CoV_2_Global_all_S1F_21358_summary_plot.png

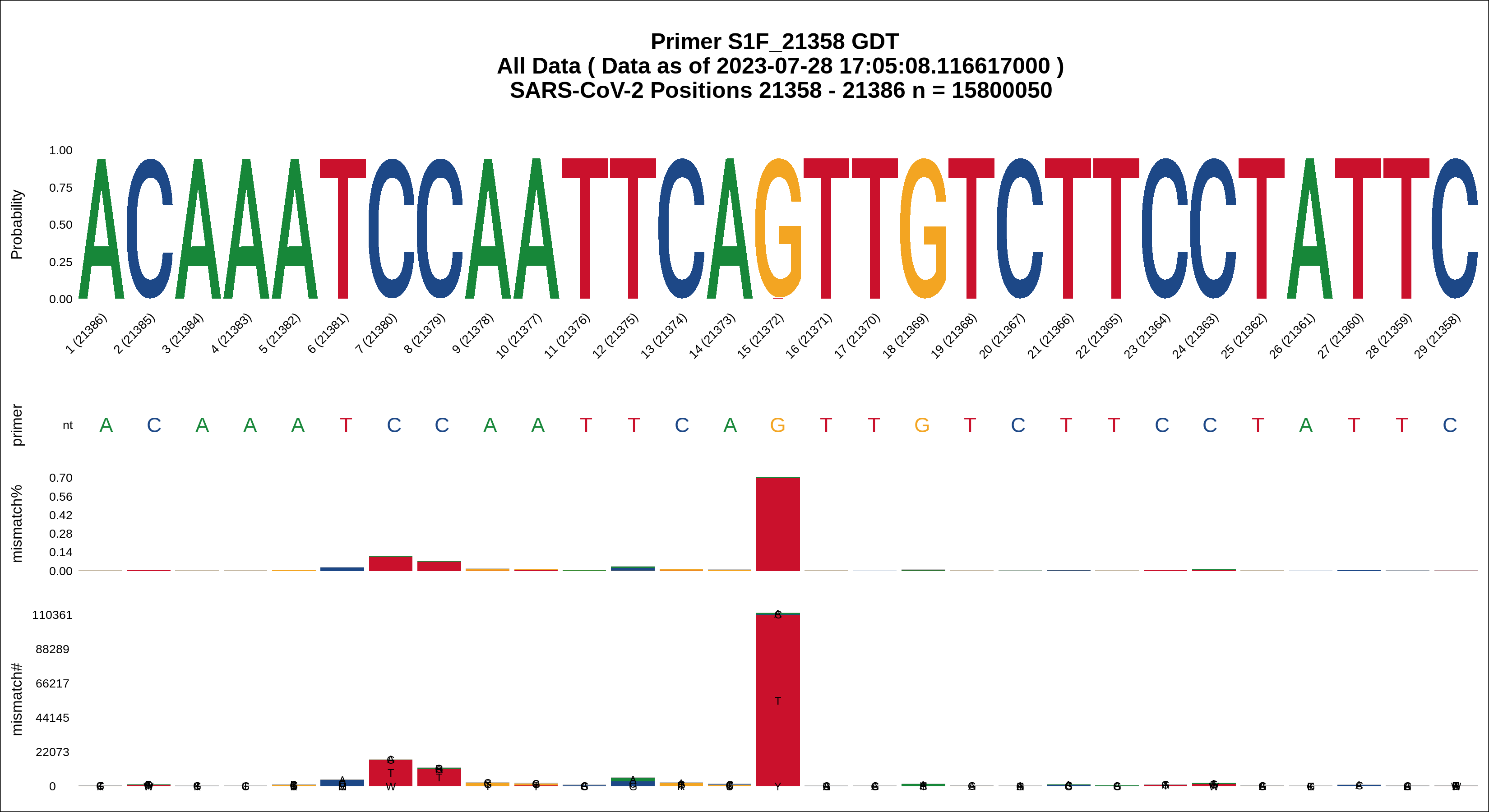

### Figure_S14_SARS_CoV_2_Global_all_S1R_23813_summary_plot.png

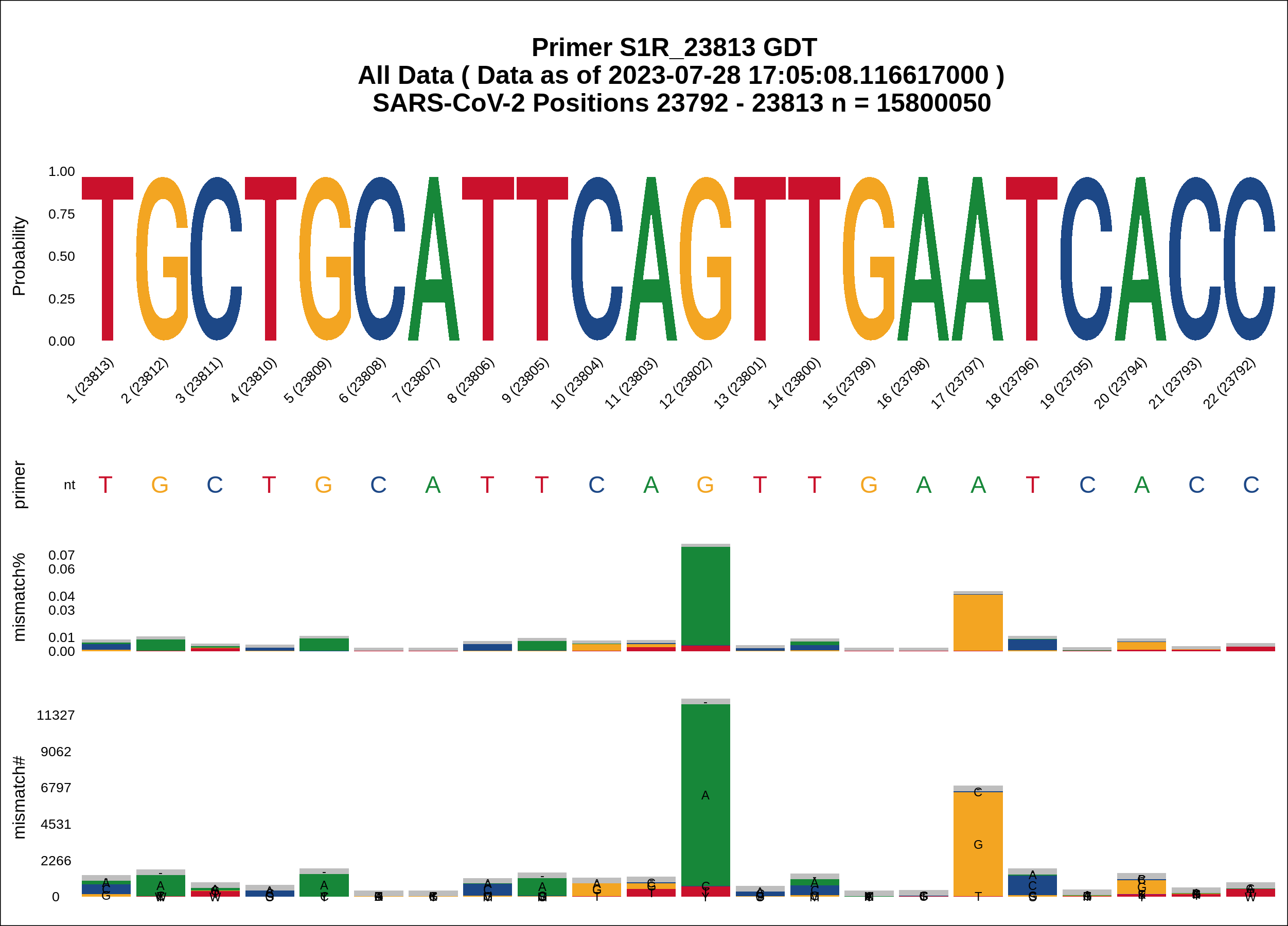

### Figure_S15_SARS_CoV_2_Global_all_S2F_23288_summary_plot.png

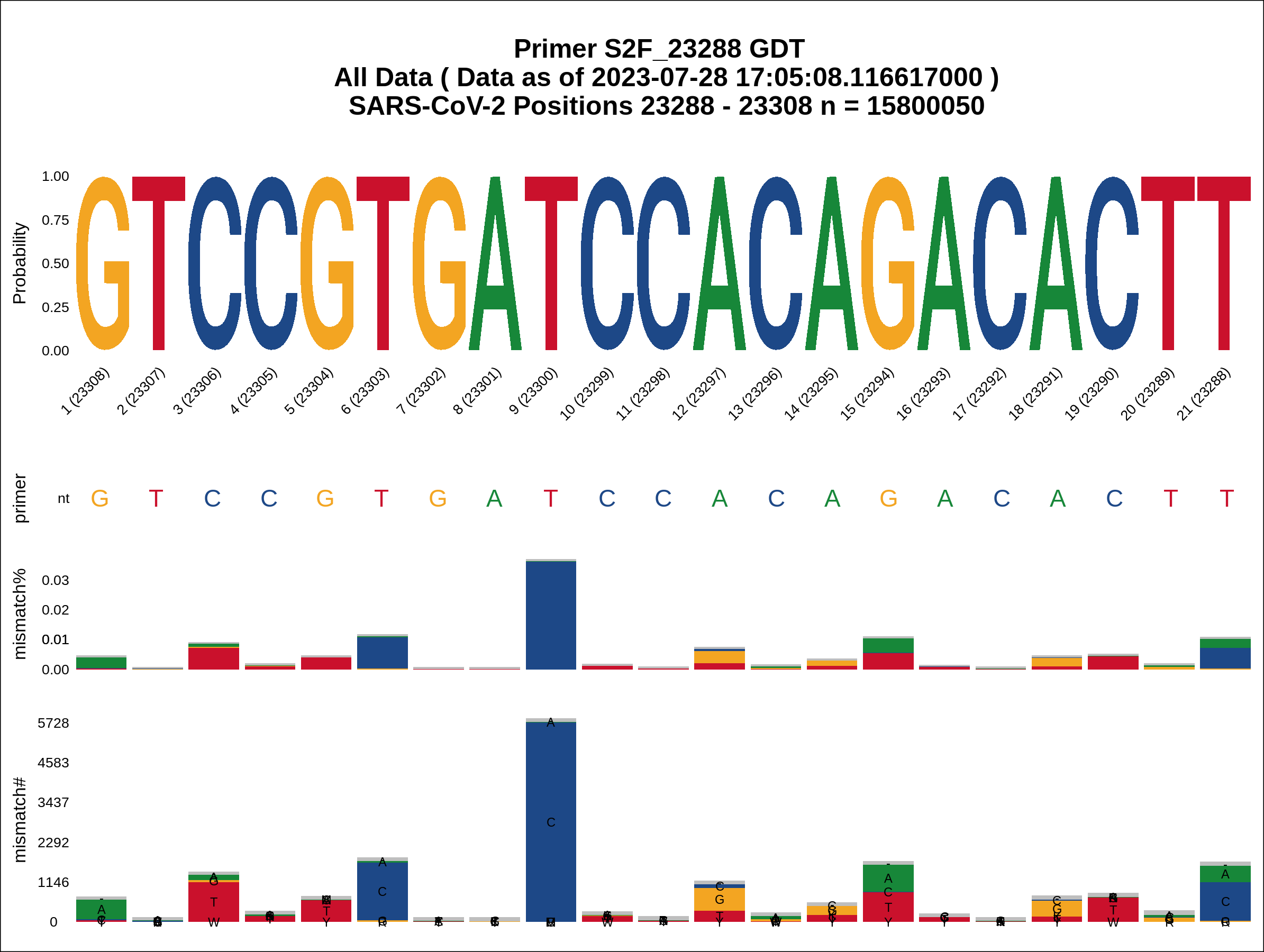

### Figure_S16_SARS_CoV_2_Global_all_S2R_25460_summary_plot.png

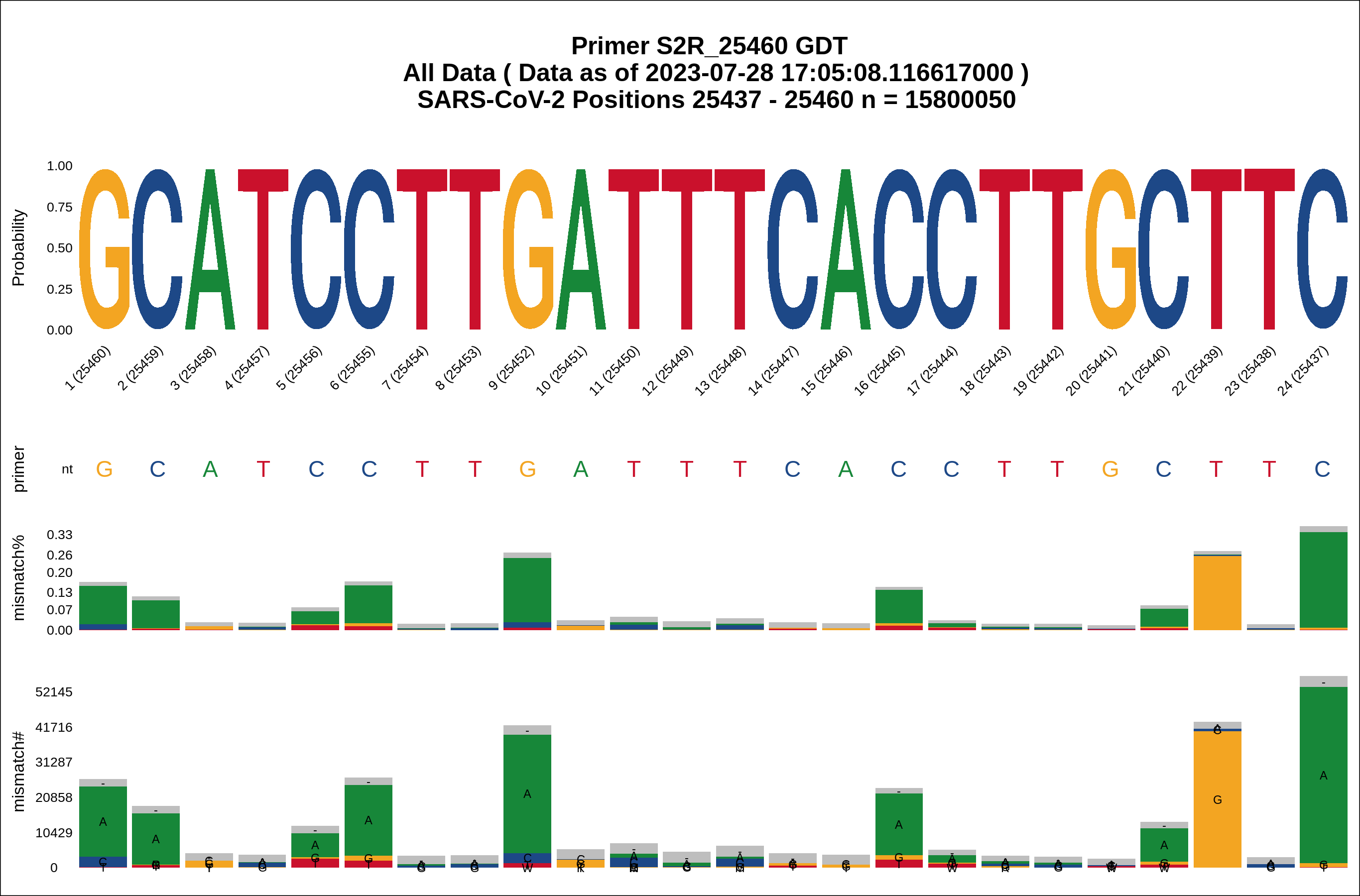

### Figure_S17_RelatedVirusAlignment.png

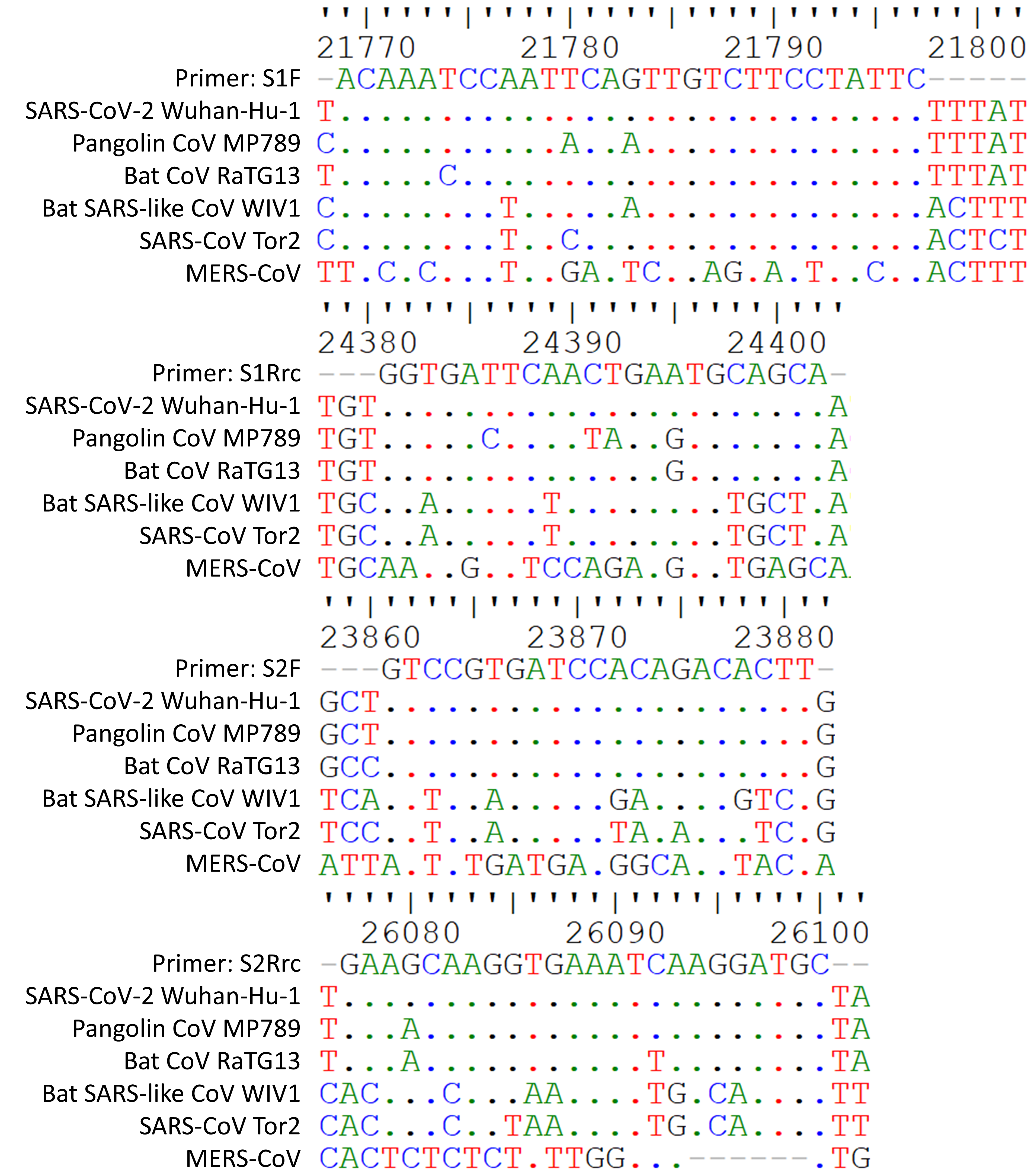

### Figure_S18_LOD-MIN.PNG

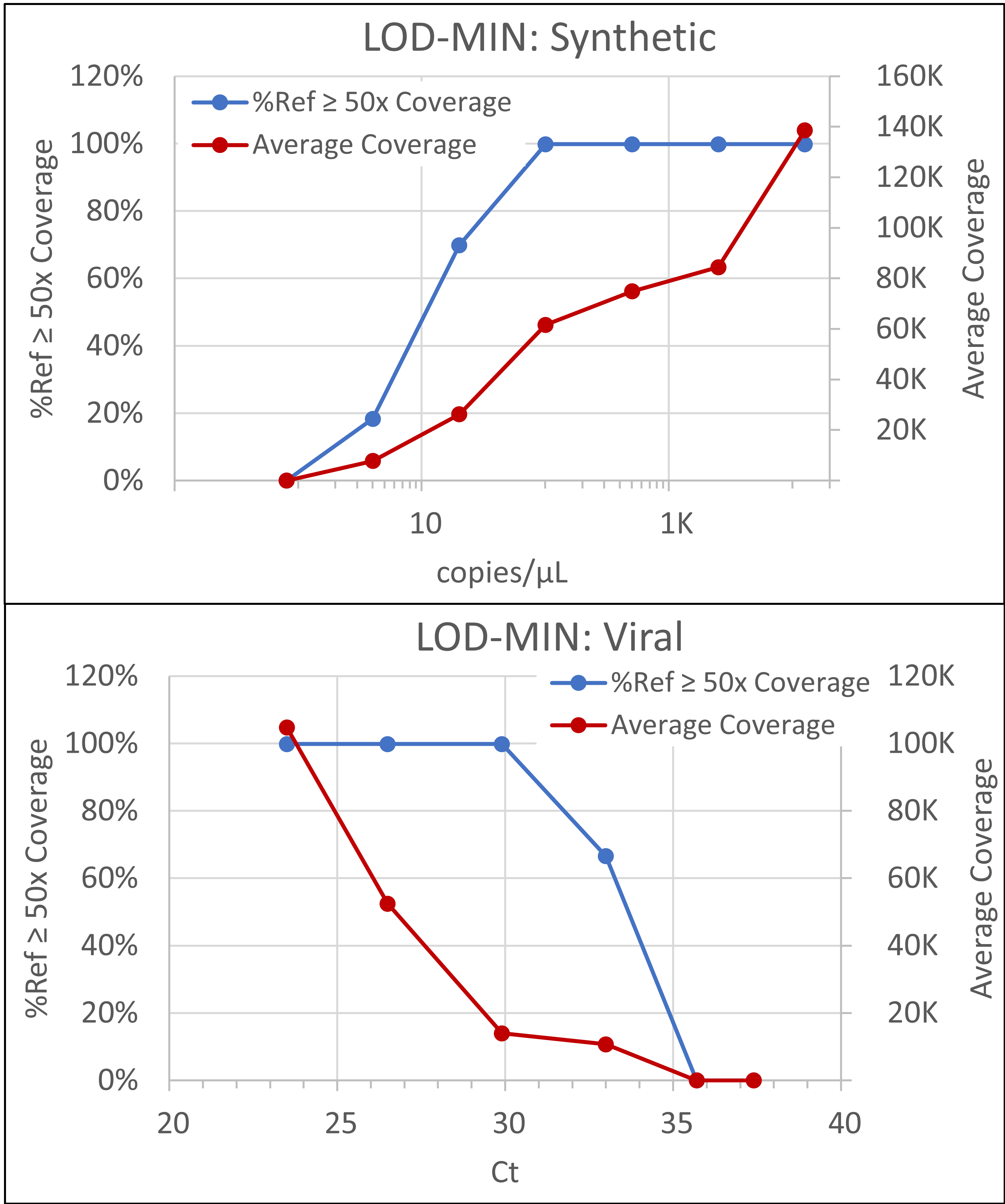

### Figure_S19_LOD-FLG.PNG

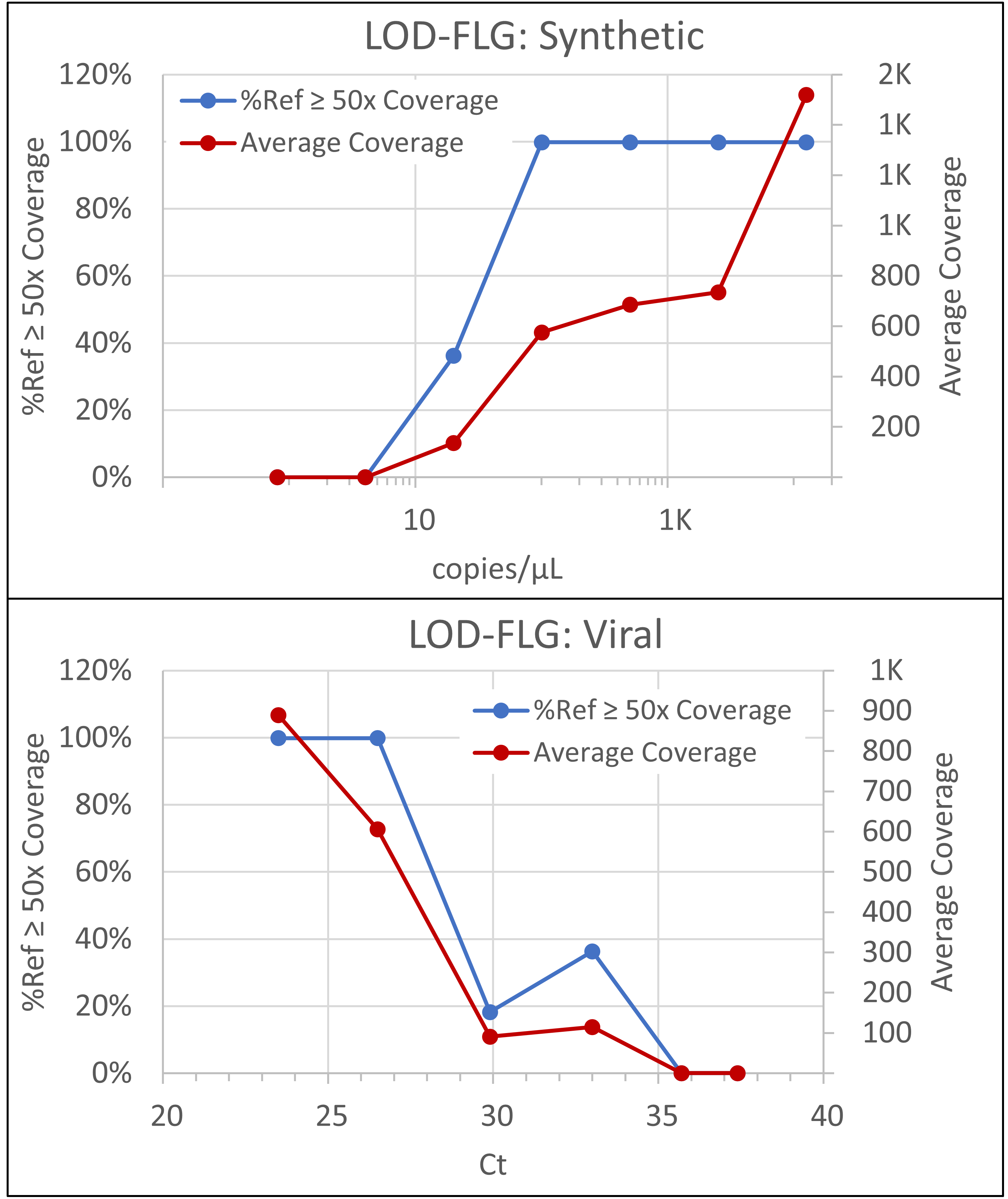

### Figure_S20_WaterNTCs.PNG

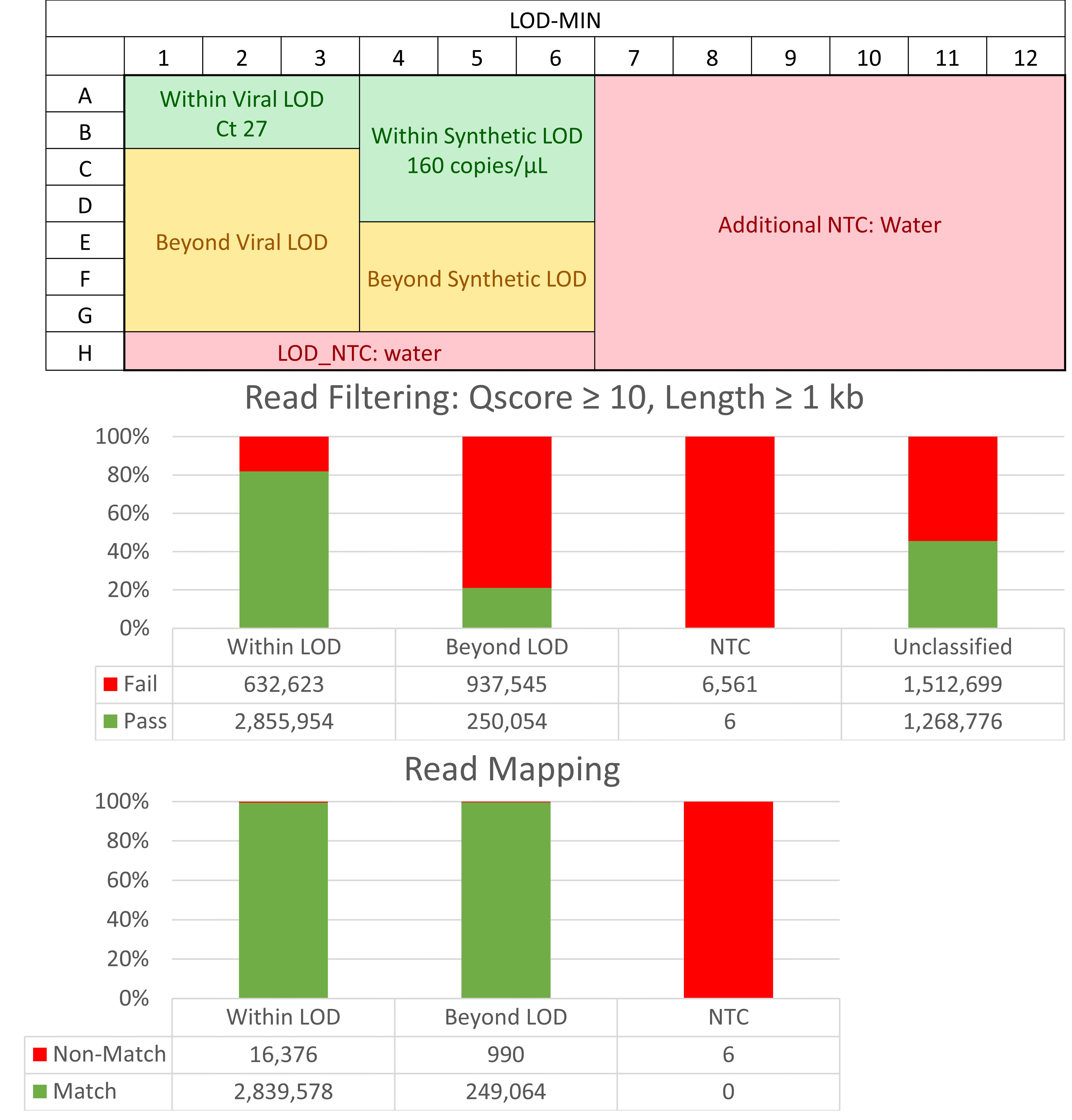

### Figure_S21_A549NTCs.PNG

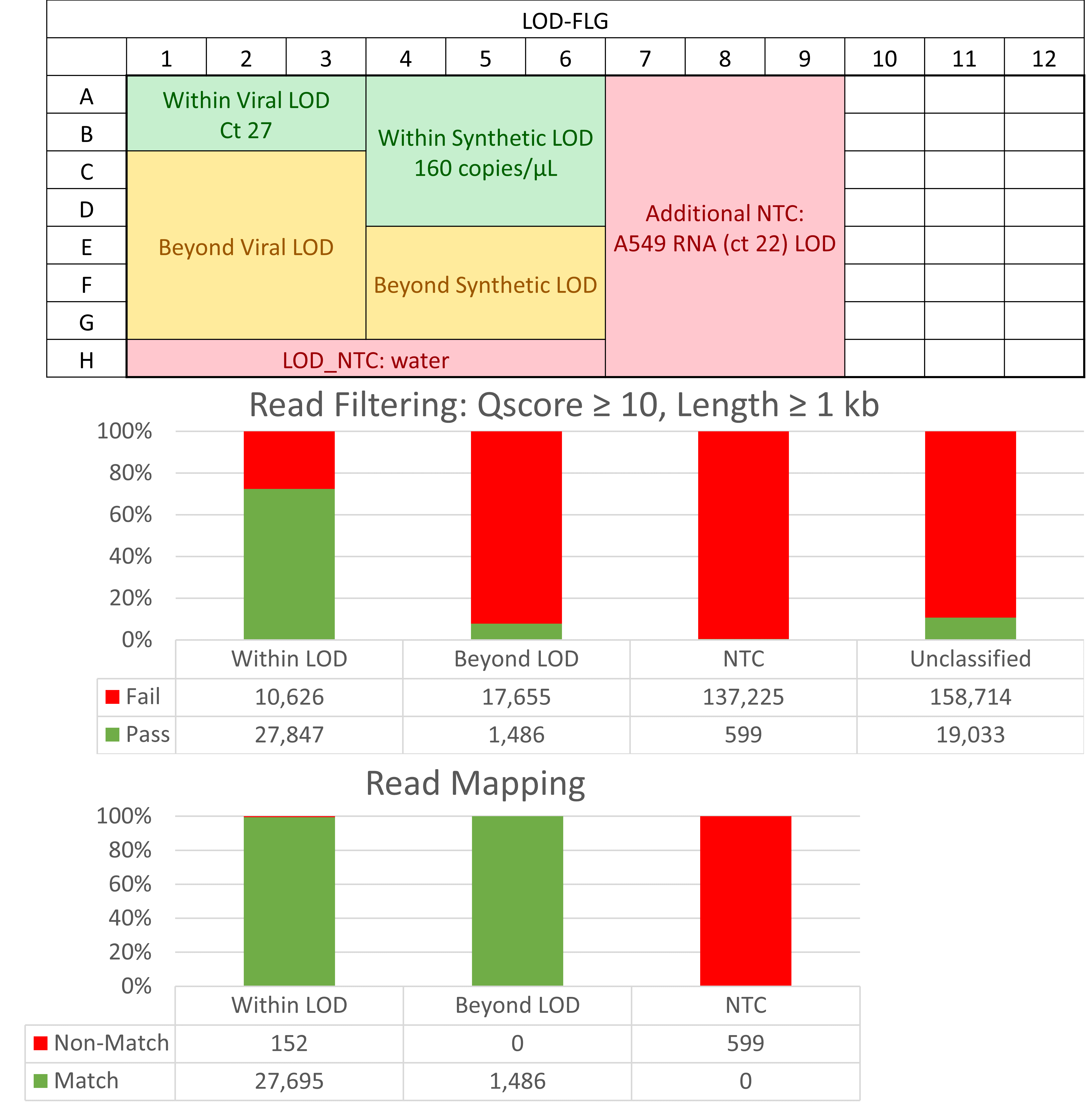

### Figure_S22_PassRateVsCt.png

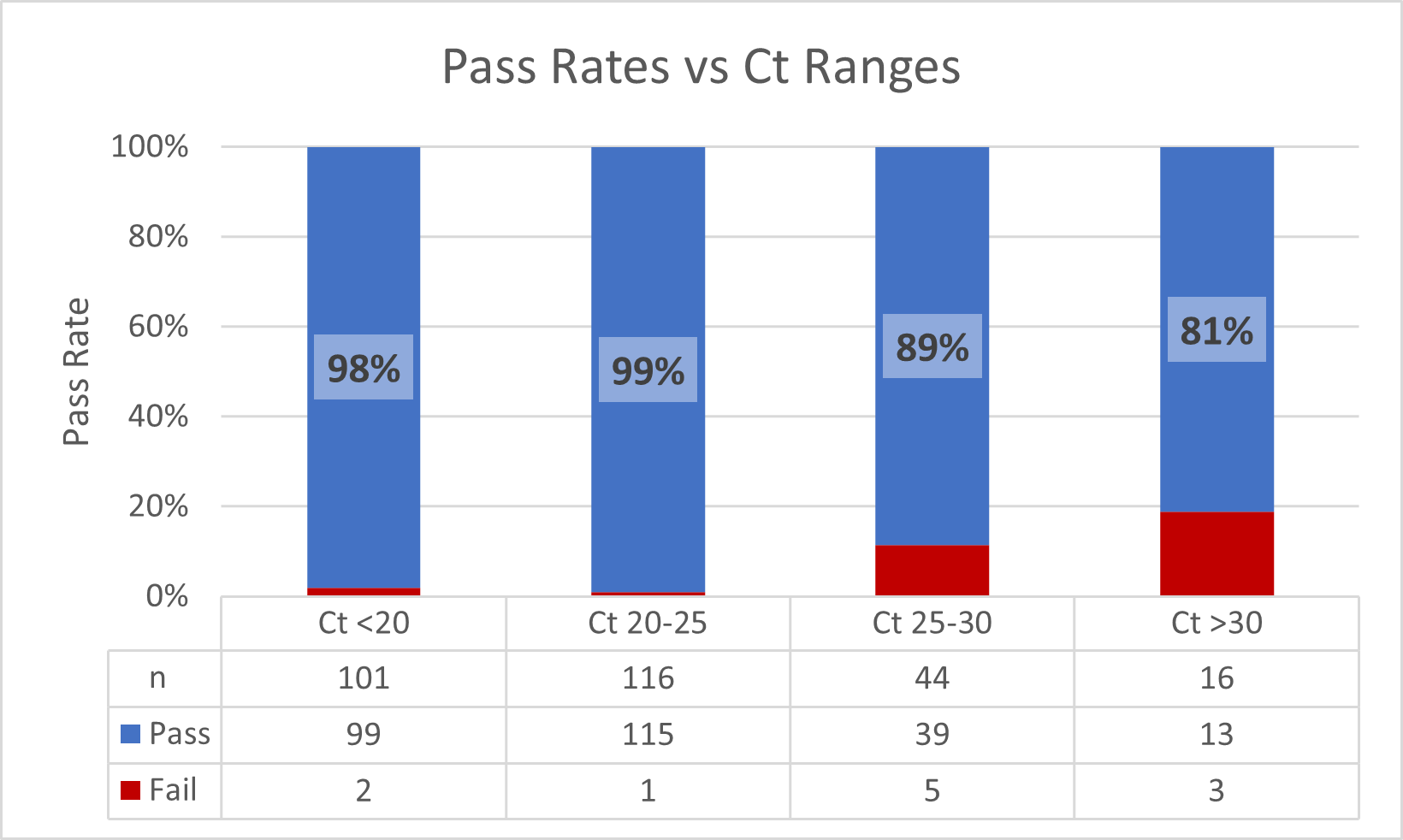

### Figure_S23_MINvFLG.png

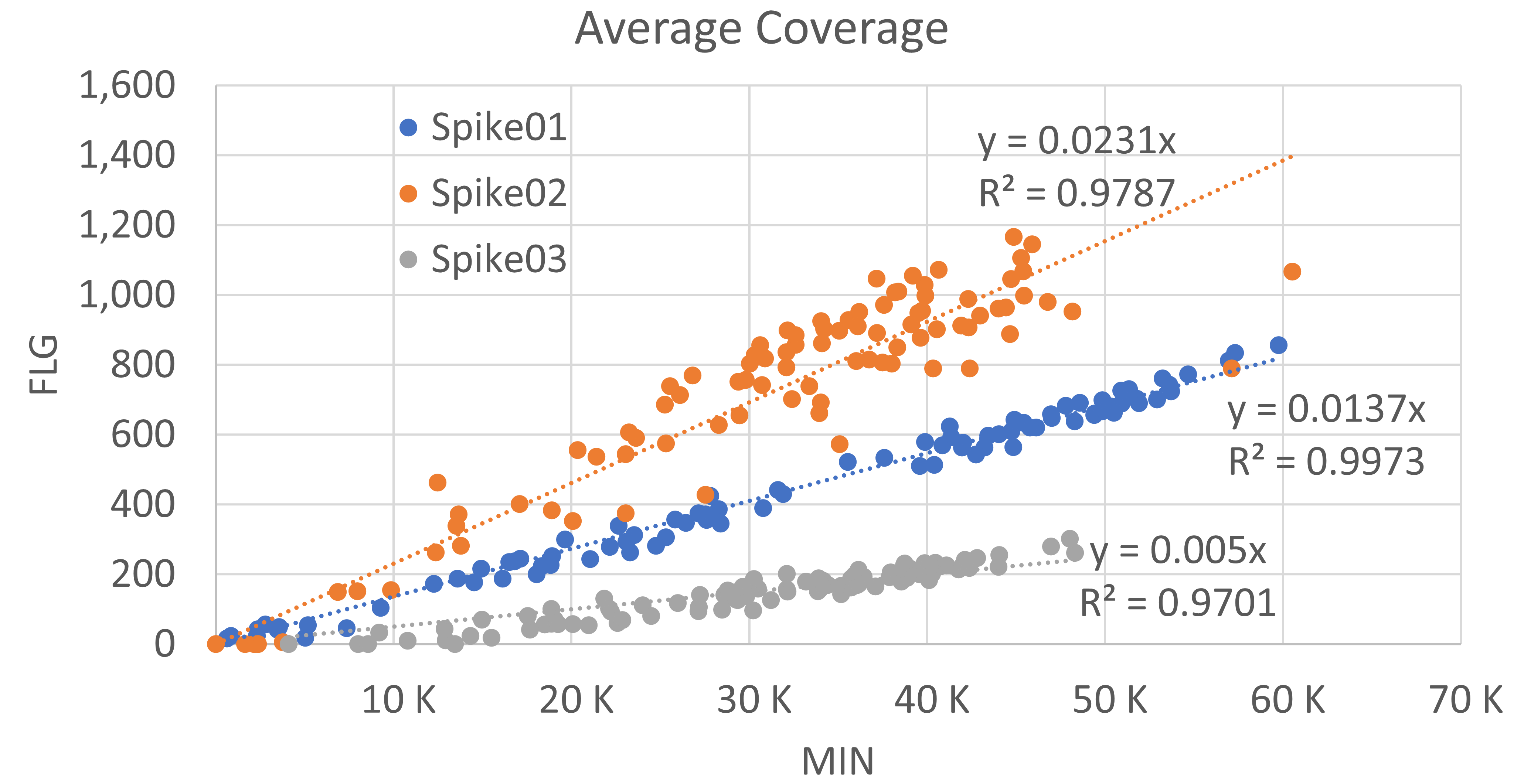

### Figure_S24_SpikeSeq-NS3-Tanglegram.png

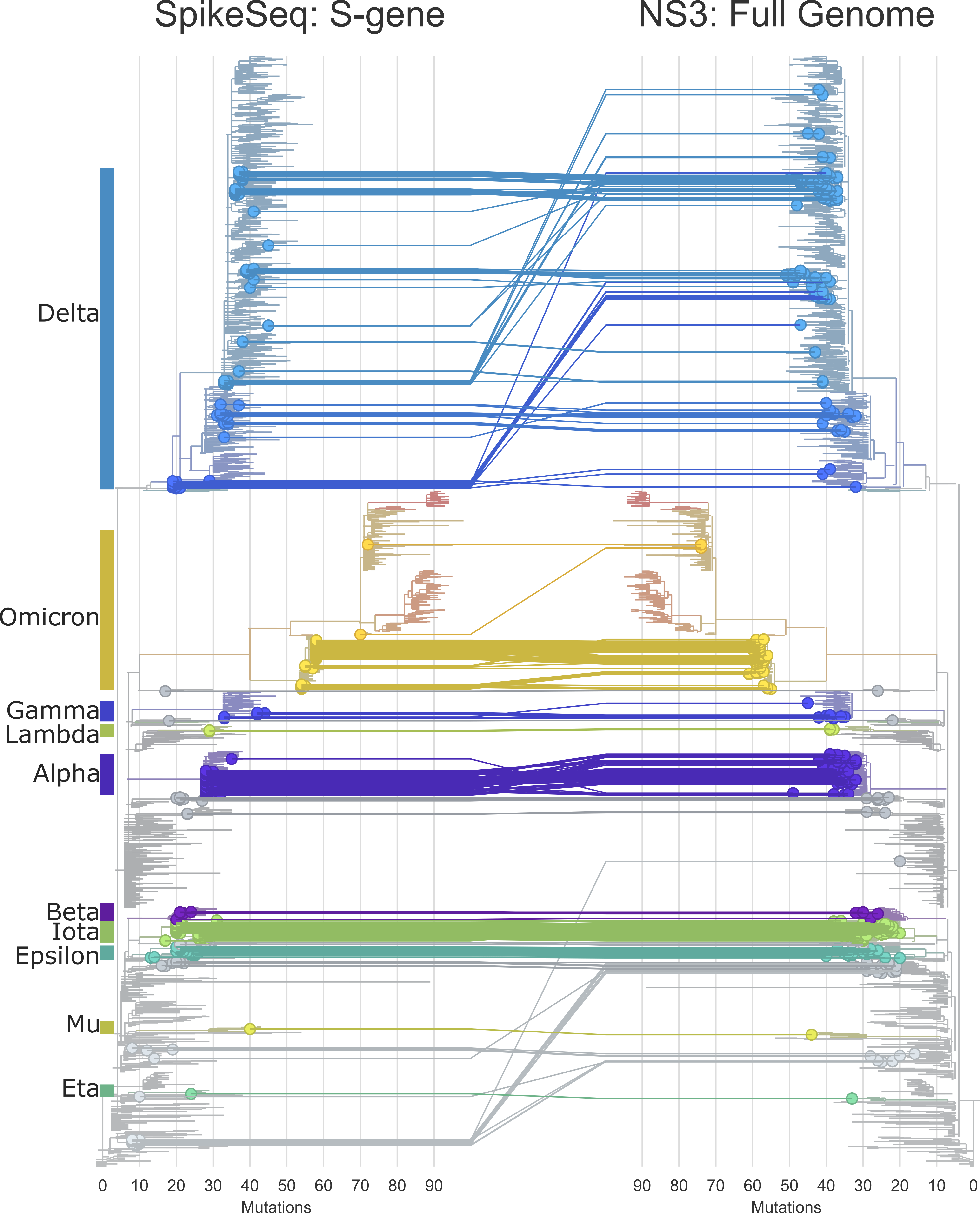

### Figure_S25_SeqCompletion.PNG

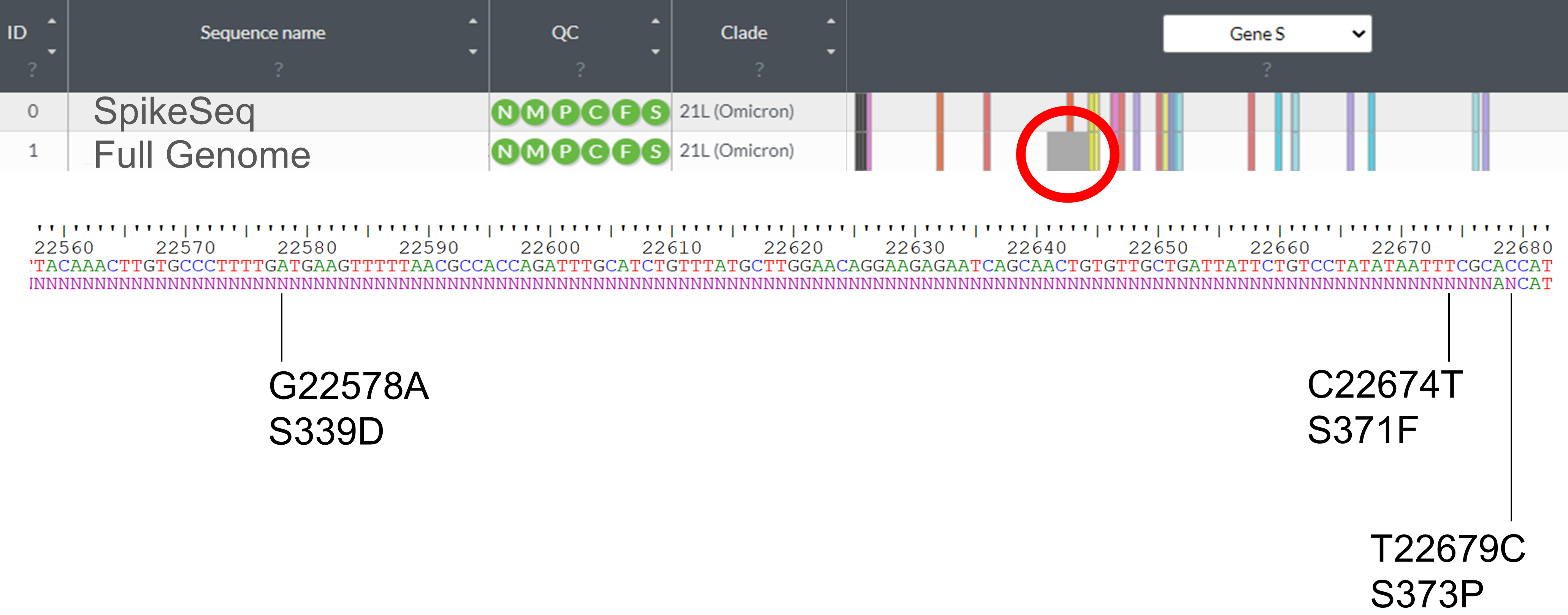
